# Cryo-EM Structures of Respiratory *bc*_1_-*cbb*_3_ type CIII_2_CIV Supercomplex and Electronic Communication Between the Complexes

**DOI:** 10.1101/2020.06.27.175620

**Authors:** Stefan Steimle, Trevor VanEeuwen, Yavuz Ozturk, Hee Jong Kim, Merav Braitbard, Nur Selamoglu, Benjamin A. Garcia, Dina Schneidman-Duhovny, Kenji Murakami, Fevzi Daldal

## Abstract

The respiratory electron transport complexes convey electrons from nutrients to oxygen and generate a proton-motive force used for energy (ATP) production in cells. These enzymes are conserved among organisms, and organized as individual complexes or combined forming large super-complexes (SC). Bacterial electron transport pathways are more branched than those of mitochondria and contain multiple variants of such complexes depending on their growth modes. The Gram-negative species deploy a mitochondrial-like cytochrome *bc*_1_ (Complex III, CIII_2_), and may have bacteria-specific *cbb*_3_-type cytochrome *c* oxidases (Complex IV, CIV) in addition to, or instead of, the canonical *aa*_3_-type CIV. Electron transfer between these complexes is mediated by two different carriers: the soluble cytochrome *c*_2_ which is similar to mitochondrial cytochrome *c* and the membrane-anchored cytochrome *c*_y_ which is unique to bacteria. Here, we report the first cryo-EM structure of a respiratory *bc*_1_-*cbb*_3_ type SC (CIII_2_CIV, 5.2Å resolution) and several conformers of native CIII_2_ (3.3Å resolution) from the Gram-negative bacterium *Rhodobacter capsulatus*. The SC contains all catalytic subunits and cofactors of CIII_2_ and CIV, as well as two extra transmembrane helices attributed to cytochrome *c*_y_ and the assembly factor CcoH. Remarkably, some of the native CIII_2_ are structural heterodimers with different conformations of their [2Fe-2S] cluster-bearing domains. The unresolved cytochrome *c* domain of *c*_y_ suggests that it is mobile, and it interacts with CIII_2_CIV differently than cytochrome *c*_2_. Distance requirements for electron transfer suggest that cytochrome *c*_y_ and cytochrome *c*_2_ donate electrons to heme *c*_p1_ and heme *c*_p2_ of CIV, respectively. For the first time, the CIII_2_CIV architecture and its electronic connections establish the structural features of two separate respiratory electron transport pathways (membrane-confined and membrane-external) between its partners in Gram-negative bacteria.

## Introduction

Mitochondrial and bacterial respiratory chains couple exergonic electron transport from nutrients to the terminal acceptor oxygen (O_2_) through a set of enzyme complexes. Concomitantly, they generate a proton motive force used for ATP synthesis and other energy-dependent cellular processes. The mitochondrial respiratory chain consists of four complexes. Complex I (NADH dehydrogenase) and Complex II (succinate dehydrogenase) are the entry points of reducing equivalents (NADH and FADH_2_) derived from nutrients into the chain. They reduce the hydrophobic electron carrier quinone (Q). Reduced quinone (QH_2_) moves rapidly within the membrane to Complex III (cytochrome (cyt) *bc*_1_ or CIII_2_) which oxidizes it and reduces the electron carrier cyt *c*. The reduced cyt *c* diffuses to Complex IV (cyt *c* oxidase or CIV) which oxidizes it and subsequently reduces the terminal electron acceptor oxygen to water (Nicholls and Ferguson, 2013) (**Fig. 1A**).

**Figure 1.**
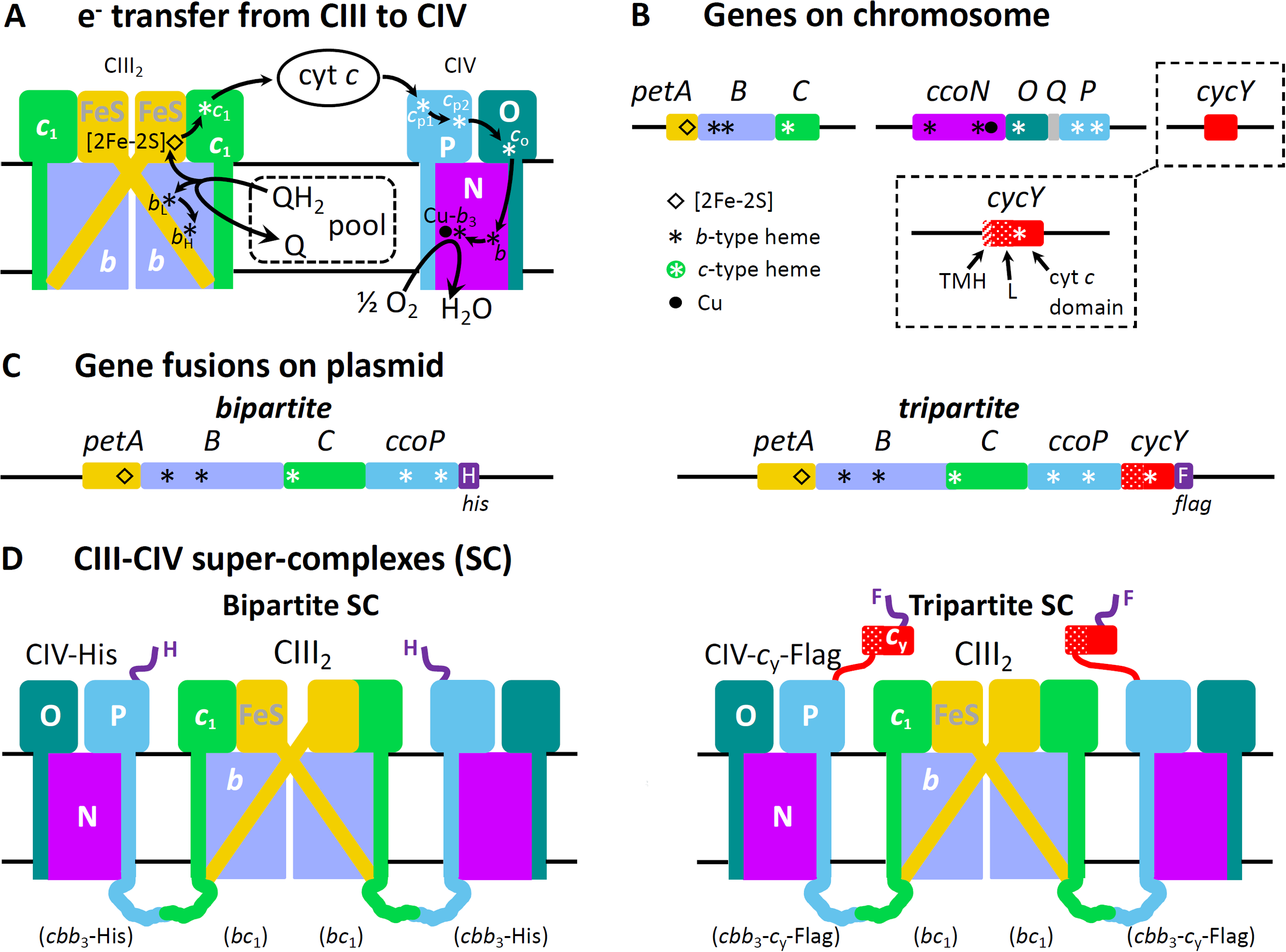
Schematic representation of fused SCs. **A.** Oxidation of QH_2_ to Q by CIII_2_ and reduction of O_2_ to H_2_O by CIV. A bifurcated electron transfer reaction conveys one electron from QH_2_ to the [2Fe-2S] cluster of the FeS protein (FeS, yellow), and another electron to hemes *b*_L_ and *b*_H_ of cyt *b* (periwinkle). The FeS protein transfers the electron from its [2Fe-2S] cluster to heme *c*_1_ on cyt *c*_1_ (green). The movement of reduced FeS protein from heme *b*_L_ to heme *c*_1_ and the electron transfer from heme *b*_H_ to Q from the pool to form a SQ (semiquinone) are not shown for clarity. An electron carrier cyt *c* (*c*_2_ or *c*_y_) receives the electron from heme *c*_1_ and delivers it to CIV. The electron arriving to CIV reaches the heme-Cu (Cu*-b*_3_) site, where O_2_ is reduced to H_2_O, via the hemes *c*_p1_ and *c*_p2_ of CcoP (P, light blue), *c*_o_ of CcoO (O, dark green) and heme *b* of CcoN (N, purple). **B.** *R. capsulatus* genes relevant to the construction of fused SCs. The *petABC* encodes the structural genes of the *bc*_1_- type CIII_2_ subunits, the FeS protein (*petA*, yellow), cyt *b* (*B*, periwinkle) and cyt *c*_1_ (*C*, green). The *ccoNOQP* encodes the structural genes of the *cbb*_3_-type CIV subunits, the CcoN (*ccoN*, purple), CcoO (*O*, dark green), CcoQ (*Q*, grey) and CcoP (*P*, light blue). The *cycY* gene (red) encodes the membrane-anchored cyt *c*_y_, and its 30-residue transmembrane helix (TMH), 69-residue linker (L) and 100- residue cyt *c* (cyt *c*) domain are indicated. Heme cofactors of *b*- and *c*-type cyts are indicated by black and white asterisks, respectively, and diamond and dot designate the [2Fe-2S] cluster and Cu atom, respectively. **C.** Plasmid-borne genetic fusions. The bipartite fusion (left) is formed by linking in-frame the 3’-end of *petC* to the 5’-end of *ccoP*, and the tripartite fusion (right) is obtained by adding in-frame the linker and cyt *c* domain of *cycY* to the 3’-end of *ccoP*. Colors and cofactor symbols are as in **A**, and the His (H) and Flag (F) affinity tags (dark purple) are added at the 3’-end of the bipartite and tripartite fusion subunits, respectively. **D.** Schematic depiction of bipartite (left) and tripartite (right) super-complexes (SC). The bipartite SC encodes a *bc*_1_-type CIII_2_ dimer fused on each side to a His-tagged *cbb*_3_-type CIV. The tripartite SC also contains the Flag-tagged cyt *c* domain of *c*_y_ (red) at the end of CcoP (blue).

Respiratory complexes are evolutionarily conserved among organisms, but bacterial enzymes are structurally simpler than their mitochondrial counterparts, consisting mainly of the catalytic subunits. However, bacterial respiratory chains are more elaborate than those of mitochondria, since they contain various complexes forming branched pathways to accommodate their diverse growth modes (Melo and Teixeira, 2016). In facultative phototrophs, the mitochondrial-like *bc*_1_-type CIII_2_ is central to respiratory and photosynthetic electron transport pathways. CIII_2_ is a dimer with each monomer comprised of three subunits: the Rieske FeS (FeS) protein with a [2Fe-2S] cluster, cyt *b* with hemes *b*_H_ and *b*_L_, and cyt *c*_1_ with heme *c*_1_ cofactors (**Fig. 1A,B**). The FeS protein external domain (FeS-ED) is mobile between the **b** (close to heme *b*_L_) and **c** (close to heme *c*_1_) positions (Darrouzet et al., 2001; Esser et al., 2006). Some species such as *Rhodobacter sphaeroides* contain a mitochondrial-like *aa*_3_-type and a bacteria-specific *cbb*_3_-type CIV, a monomer comprised of four subunits: CcoN with heme *b* and heme *b*_3_-Cu binuclear center, CcoO with heme *c*_o_, CcoQ, and CcoP with hemes *c*_p1_ and *c*_p2_ cofactors (**Fig. 1A,B**). Other species such as *Rhodobacter capsulatus* (Khalfaoui-Hassani et al., 2016) and pathogens like *Helicobacter pylori* and *Campylobacter jejuni* (Smith et al., 2000), *Neisseria* (Aspholm et al., 2010) have only a high oxygen affinity *cbb*_3_-type CIV to support their micro-aerophilic growth.

Besides the mitochondrial-like soluble and diffusible cyt *c*, many Gram-negative bacteria contain additional electron carriers that are membrane-anchored via transmembrane domains (*e.g*., *Rhodobacter capsulatus* cyt *c*_y_ (Jenney and Daldal, 1993), *Paracoccus denitrificans* cyt *c*_552_ (Turba et al., 1995), and *Bradyrhizobium japonicum* CycM (Bott et al., 1991)) or fatty acids (*e.g*., *Blastochloris viridis* tetraheme cyt *c* (Weyer et al., 1987) and *Helicobacterium gestii* cyt *c*_553_ (Albert et al., 1998)). Conversely, Gram-positive bacteria are devoid of freely diffusing electron carriers. Instead, they may have additional cyt *c* domains fused to their CIII_2_ (*i.e*., *bcc*-type) such as in *Mycobacterium smegmatis* (Kim et al., 2015) and *Corynebacterium glutamicum* (Niebisch and Bott, 2003) or CIV (*i.e*., *caa*_3_-type) such as in *Bacillus subtilis* (Winstedt and von Wachenfeldt, 2000) and *Bacillus stearothermophilus* (Sakamoto et al., 1996). Bacterial electron carrier cyts *c* are involved in multiple metabolic pathways. Both the diffusible cyt *c* (*e.g*., *R. capsulatus* cyt *c*_2_ or its homologs) and the membrane-anchored cyt *c* (*e.g*., *R. capsulatus c*_y_ or its homologs) electronically connect CIII_2_ to the photochemical reaction center in photosynthesis (Daldal et al., 1986), and to CIV in respiration (Hochkoeppler et al., 1995). In species like *R. sphaeroides*, cyt *c*_2_ functions in both photosynthesis and respiration, while cyt *c*_y_ is restricted to respiration (Myllykallio et al., 1999).

In recent years, the co-occurrence of individual complexes together with multi-enzyme supercomplexes (SCs) in energy-transducing membranes has become evident (Acin-Perez and Enriquez, 2014; Melo and Teixeira, 2016). However, the regulation and physiological role of this heterogeneity are debated (Brzezinski, 2019; Letts and Sazanov, 2017; Milenkovic et al., 2017). SCs may stabilize individual complexes, enhance catalytic efficiency through substrate/product channeling, or minimize production of harmful intermediates (*e.g*., reactive oxygen species) to decrease cellular distress (Enriquez, 2016; Letts et al., 2019; Quintana-Cabrera and Soriano, 2019). Mitochondrial SCs, such as CICIII_2_CIV (respirasomes) or their smaller variants containing only CICIII_2_ (Sousa and Vonck, 2019) or CIII_2_CIV (Letts et al., 2016) have established molecular architectures (Gu et al., 2016; Hartley et al., 2019; Wu et al., 2016). Bacterial SCs, including these of *P. denitrificans* (Berry and Trumpower, 1985; Stroh et al., 2004), *Geobacillus stearothermophilus* (Bergdoll et al., 2016), *Bacillus PS3* (Sone et al., 1987), *M. smegmatis* (Kim et al., 2015), and *C. glutamicum* (Kao et al., 2016), have been characterized biochemically, but only the structure of the Gram positive *M. smegmatis* SC (CIII_2_CIV_2_) has been reported (Gong et al., 2018; Wiseman et al., 2018).

As of yet, no respiratory SC structure has been determined for Gram-negative bacteria, the evolutionary precursors of mitochondria. Furthermore, SCs containing ancient forms of CIV (*i.e*., *cbb*_3_-type) representing primordial features of respiratory chains with multiple electron carriers are unknown (Ducluzeau et al., 2008). Structural studies of such SCs have been hampered due to unstable interaction between CIII_2_ and CIV, hence their trace amounts in nature. We have overcome this hurdle using a genetic approach, yielding large amounts of SCs from the Gram negative facultative phototroph *R. capsulatus*. Here, we report the first cryo-EM structure of a respiratory *bc*_1_-*cbb*_3_ type SC (CIII_2_CIV, at 5.2Å resolution), as well as several conformers of native CIII_2_ (at 3.3-4.2Å resolution). We define the interaction regions of cyt *c*_2_ and cyt *c*_y_ within the SC by combining cryo-EM, cross-linking mass spectrometry (XL-MS) and integrative structure modeling. We propose that the membrane-bound cyt *c*_y_ donates electrons to heme *c*_p1_, while the diffusible cyt *c*_2_ transfers them to heme *c*_p2_, of CcoP subunit of CIV. For the first time, this work establishes the structural features of CIII_2_CIV and its two distinct respiratory electron transport pathways (membrane-confined and membrane-peripheral) connecting its partners in Gram-negative bacteria.

## Results

### Stabilization, isolation, and composition of functional fused SCs

Earlier studies on soluble cyt *c*-independent electron transport pathways have indicated that in some species (*e.g*., *R. capsulatus* (Myllykallio et al., 2000)), CIII_2_, CIV, and the membrane-anchored cyt *c*_y_ are in close proximity to each other. BN-PAGE of membranes from a wild type strain of *R. capsulatus*, overstained for CIV-specific in-gel activity, showed barely detectable bands around ∼450 kDa M_r_ (**Fig. S1A**). The masses of these bands were larger than that of the CIV monomer (∼100 kDa, running as ∼230 kDa on BN-PAGE) or CIII_2_ dimer (∼200 kDa, running as >250 kDa on BN-PAGE), suggesting the occurrence of large SCs. However, these entities were of low abundance and highly unstable, rendering their study difficult. In our earlier work, translationally fusing cyt *c*_1_ of CIII_2_ to cyt *c*_y_ had produced an active *bcc*-type CIII_2_ (*i.e*., cyt *bc*_1_-*c*_y_ fusion) (Lee et al., 2008), suggesting that this approach might also be used to stabilize the interactions between CIII_2_ and CIV.

During the assembly processes of CIII_2_ and CIV, cyt *c*_1_ interacts with cyt *b* to form a cyt *b*-*c*_1_ subcomplex (Davidson et al., 1992), and CcoP associates with CcoNOQ subcomplex to yield an active CIV (Kulajta et al., 2006). We thought that translationally fusing the C-terminus (C-ter) of cyt *c*_1_ to the N-terminus (N-ter) of CcoP, which are on the inner (*n*) side of the membrane, forming a bipartite cyt *c*_1_-CcoP fusion protein might produce a stable bipartite *bc*_1_-*cbb*_3_ type SC (left panels of **Fig. 1C,D**). Furthermore, adding the natural 69-residue linker (L) and the 100-residue cyt *c* domain of *c*_y_ to the C-ter of cyt *c*_1_-CcoP, which is on the outer (*p*) side of the membrane, to form a tripartite cyt *c*_1_-CcoP-*c*_y_ fusion protein might yield a tripartite *bc*_1_-*ccbb*_3_ type SC with an attached electron carrier (right panels of **Fig. 1C,D**). This approach (see Supplemental Information, Methods, for details) yielded fusion constructs (**Fig. S1B**) that functionally complemented a mutant lacking CIII_2_ and CIV for photosynthesis-proficiency (*i.e*., CIII_2_ activity) and CIV activity (**Fig. S1C**).

The His-tagged bipartite and Flag-tagged tripartite SCs were purified from detergent-dispersed membranes by tag-affinity and size exclusion chromatography (SEC) (SI, Methods) (**Fig. 2A,B**). BN-PAGE of isolated proteins showed that the A-1 and B-1 fractions contained mostly the large entities of M_r_ ∼450 kDa range (**Fig. 2A,B**, insets), and SDS-PAGE revealed that they had the cyt *c*_1_-CcoP (∼65 kDa) or cyt *c*_1_-CcoP-*c*_y_ fusion proteins (∼80 kDa) (**Fig. 2C**). All protein bands seen in **Fig. 2C** were identified by mass spectrometry (MS) (**Table S3**) and assigned to the subunits of CIII_2_ and CIV. The fusion proteins also contained covalently-attached heme cofactor(s), as shown by 3,3’,5,5’-*tetramethyl-benzidine* (TMBZ) staining, which is specific to covalent heme containing *c*-type cyts **(Fig. 2D**). CcoQ (M_r_ ∼7kDa) of CIV was absent in both SC preparations.

**Figure 2.**
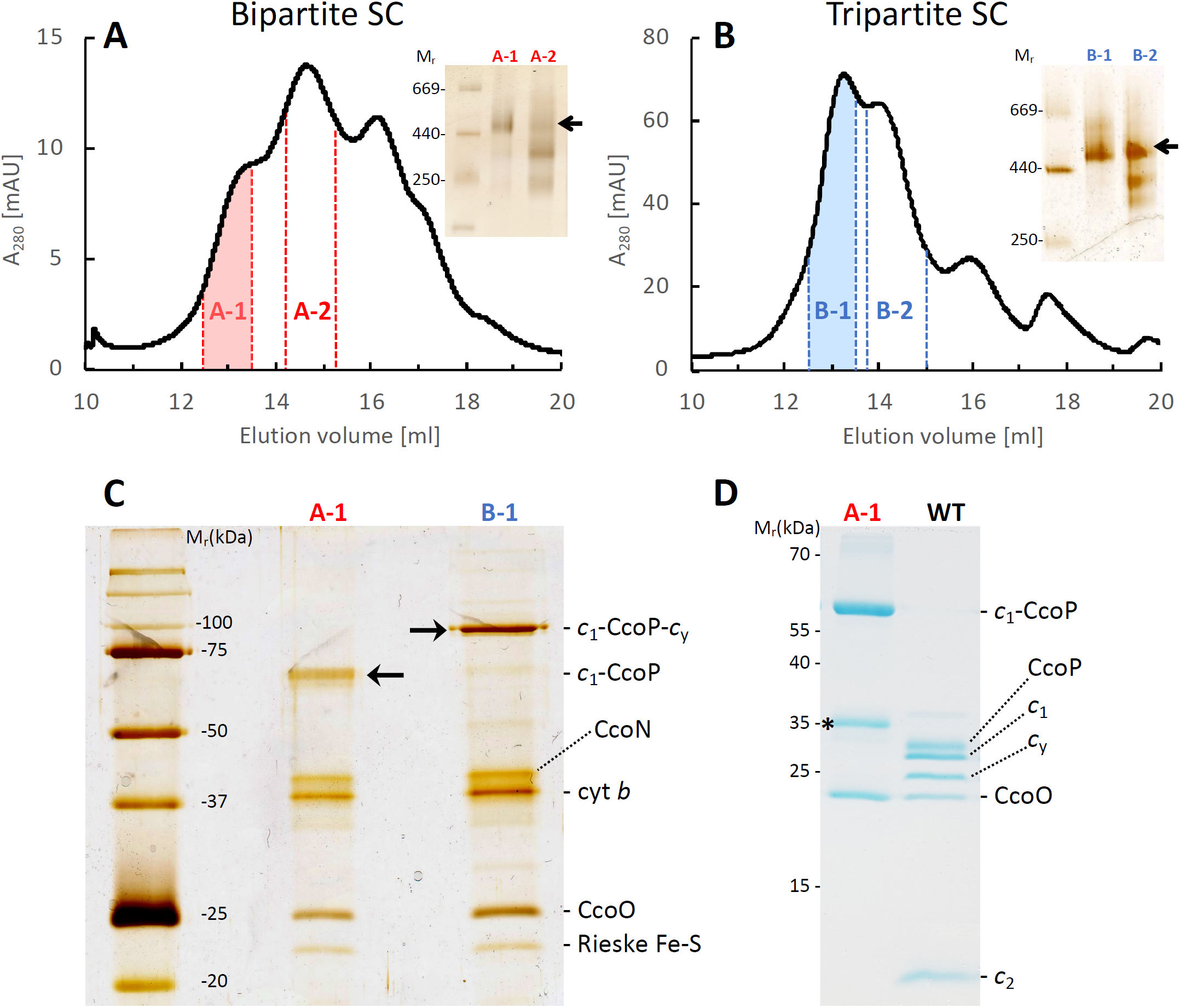
Purification and characterization of bipartite and tripartite SCs. The SEC elution profiles of the bipartite SC (**A**) and tripartite SC (**B**) are shown. In each case, the fractions 1 and 2 were analyzed by 4-16% Native PAGE (insets) and silver staining, and only the fractions A-1 and B-1 were used for cryo-EM studies. The bands at ∼450/480 kDa corresponding to bipartite and tripartite SCs are indicated by arrows. **C.** Fractions A-1 and B-1 were separated by SDS-PAGE, silver stained, and protein bands identified by mass spectrometry. The fused bipartite *c*_1_-CcoP (A-1) and tripartite *c*_1_-CcoP-*c*_y_ (B-1) subunits are indicated by arrows. Note the absence of *c*_1_ and CcoP subunits in both cases. **D**. Peak A-1 and DDM-dispersed membranes from wild type *R. capsulatus* (WT) were analyzed by SDS-PAGE/TMBZ to reveal the covalently attached heme cofactors. The tripartite construct (B-1) is virtually identical to A-1, except that the *c*_1_-CcoP band is replaced by *c*_1_-CcoP-*c*_y_. All *c*-type cyts are labeled, and the additional band indicated by * corresponds to a proteolytic cleavage product of the c_1_-CcoP fusion subunit.

### Spectral and functional characterization of SCs

Purified SCs were characterized by optical redox difference spectra for their total cyt *b* and cyt *c* contents. The spectra were distinct from those of CIII_2_ (Valkova-Valchanova et al., 1998) or CIV (Gray et al., 1994), and the tripartite SC contained more heme *c* than the bipartite SC, due to the additional cyt *c* domain of *c*_y_ (**Fig. S2A**). Both SC preparations exhibited 2,3-dimethoxy-5-methyl-6-*decyl*-1,4-*benzoquinone* (DBH_2_):cyt *c* reductase activity (12.4 +/-1.8 μmol/mg of protein/min and 7.2 +/-2.1 μmol/mg of protein/min for bipartite and tripartite SCs, respectively), which is specific to *bc*_1_-type CIII_2_ (**Fig. S2B)**. They also had cyt *c*:O_2_ reductase activity (0.46 +/-0.14 μmol/mg of protein/min and 2.9 +/-0.7 μmol/mg of protein/min for bipartite and tripartite SCs, respectively), which is specific to *cbb*_3_-type CIV (**Fig. S2C**). Importantly, the tripartite SC exhibited DBH_2_ and O_2_ dependent DBH_2_:O_2_ reductase (*i.e*., coupled CIII_2_+CIV) activity (0.143 +/-0.025 μmol of O_2_ consumed/mg of protein/min) without addition of a soluble electron carrier (*e.g*., horse heart cyt *c* or *R. capsulatus* cyt *c*_2_) (**Fig. S2D**), unlike the bipartite SC that required it. Remarkably, the cyt *c* domain of *c*_y_ fused to cyt *c*_1_-CcoP transferred electrons from CIII_2_ to CIV.

### Structures of the tripartite SCs

We first focused on cryo-EM analysis of the tripartite SC preparations that were more stable and abundant than the bipartite SCs (**Fig. 2B**, fraction B-1). Initial 3D classes were of primarily two different sizes (**Fig. S3**, Box 1, left). The smaller (∼180Å length) particles were asymmetrical, and their size and shape suggested that they may correspond to a dimeric CIII_2_ associated with a single CIV. Focused classification and processing of the subclass containing ∼62,000 particles with the highest initial resolution, and best discernable features, led to a tripartite CIII_2_CIV map (SC-1A, EMD-22228) at 6.1Å resolution (**Fig. S3A**, see SI Methods for details), while another dataset yielded a slightly lower resolution map (SC-1B, EMD-22230) at 7.2Å (**Fig. S3B**) (**Table 1**). The larger particles (∼250Å length, **Fig. S3**, Box 1) were more symmetrical and represented a dimeric CIII_2_ flanked by two CIV (*i.e*., CIII_2_CIV_2_), as expected based on two *c*_1_-CcoP-*c*_y_ subunits *per* CIII_2_. However these particles were rare (∼5,000) and their map (SC-1C) could not be refined beyond ∼10Å resolution (**Fig. S3C**).

**Table 1:** Statistics of data collection and 3D reconstruction of CIII_2_CIV SC. Six and two individual datasets were combined for the tripartite (∼c_y_, fused cyt *c* domain of cyt *c*_y_) and the bipartite (+ c_y_, supplemented with native cyt *c*_y_) SCs, respectively. For the combined datasets, the parameters for data collection were identical, except the exposure time. For all combined datasets, the range of exposure times and the corresponding dose rates are provided.

| Data Collection |  |  |  |
| --- | --- | --- | --- |
| Sample | Tripartite<br>CIII<sub>2</sub>CIV~c<sub>y</sub> | Bipartite<br>CIII<sub>2</sub>CIV + c<sub>y</sub> |  |
| Number of micrographs | 17390 | 17680 |  |
| Electron microscope | FEI Titan Krios | FEI Titan Krios |  |
| Voltage (kV) | 300 | 300 |  |
| Electron detector | K2 | K3 |  |
| Pixel size (Å) | 1.32 | 1.36 |  |
| Defocus range (µm) | 0.4 - 4.0 | 0.4 - 4.0 |  |
| Frames per movie | 40 | 80 |  |
| Total dose (e-/Å<sup>2</sup>) | 40 | 40 |  |
| Exposure time (s) | 9 - 15 | 3.0 - 3.2 |  |
| Dose rate (e-/Å<sup>2</sup>·s<sup>-1</sup>) | 2.7 - 4.4 | 12.5 - 13.2 |  |
| Dose/frame (e-/pix) | 1.74 | 0.93 |  |
| 3D Reconstruction |  |  |  |
| Map Name | SC-1A | SC-1B | SC-2A |
| Particles | 61934 | 87026 | 14978 |
| B-factor | -184 | -684 | -87 |
| Resolution range (Å) | 4.5 - 9 | 4.8 - 9 | 4.3 – 8 |
| Final resolution (Å)<br>at FSC 0.143 (0.5) | 6.09 (7.14) | 7.20 (8.85) | 5.18 (7.35) |
| EMDB map entry | EMD-22228 | EMD-22230 | EMD-22230 |
| Model Refinement |  |  |  |
| PDB coordinate entry | 6XKX | 6XKZ | 6XKW |
| Phenix CC<sub>box/mask/volume</sub> | 0.75/0.64/0.64 | 0.71/0.59/0.58 | 0.74/0.61/0.63 |
| Refinement statistics | Models were obtained by fitting the high-resolution model of CIII<sub>2</sub> (PDB: 6XI0) and the homology model of CIV (**Table S7**) into the maps. See these tables for refinement statistics. |  |  |

The *R. capsulatus cbb*_3_-type CIV is highly homologous to that of *P. stutzeri* but not identical (see Methods for details). Thus, a homology model of CIV was built using the *P. stutzeri* structure (PDB: 3MK7; 3.2Å resolution) as a template and validated (**Table S7**) (SI, Methods). In addition, the existing CIII_2_ model (PDB: 1ZRT; 3.5Å resolution) was further refined (PDB: 6XI0; 3.3Å resolution) using our cryo-EM data (see below and **Table 2**). These models were fitted as rigid bodies into the maps SC1-A with a correlation coefficient CC_box_ of 0.75 and SC-1B with a correlation coefficient CC_box_ of 0.71 (**Fig. S4A**) (**Table 1**). The [2Fe-2S] clusters of the FeS proteins of CIII_2_ could be recognized closer to heme *b*_L_ (b position) than to heme *c* (c position), but had lower occupancy and resolution likely due to conformational heterogeneity (**Fig. S4B**). In particular, the heterogeneity of the FeS-ED in monomer A (*i.e*., adjacent to CIV) was more pronounced than that in monomer B (*i.e*., away from CIV) of CIII_2_. Lower resolutions of the FeS-ED portions were anticipated because of their mobility (Darrouzet et al., 2001; Esser et al., 2006). Details of the tripartite CIII_2_CIV structure are described below together with the bipartite SC, which has a higher resolution.

**Table 2:**
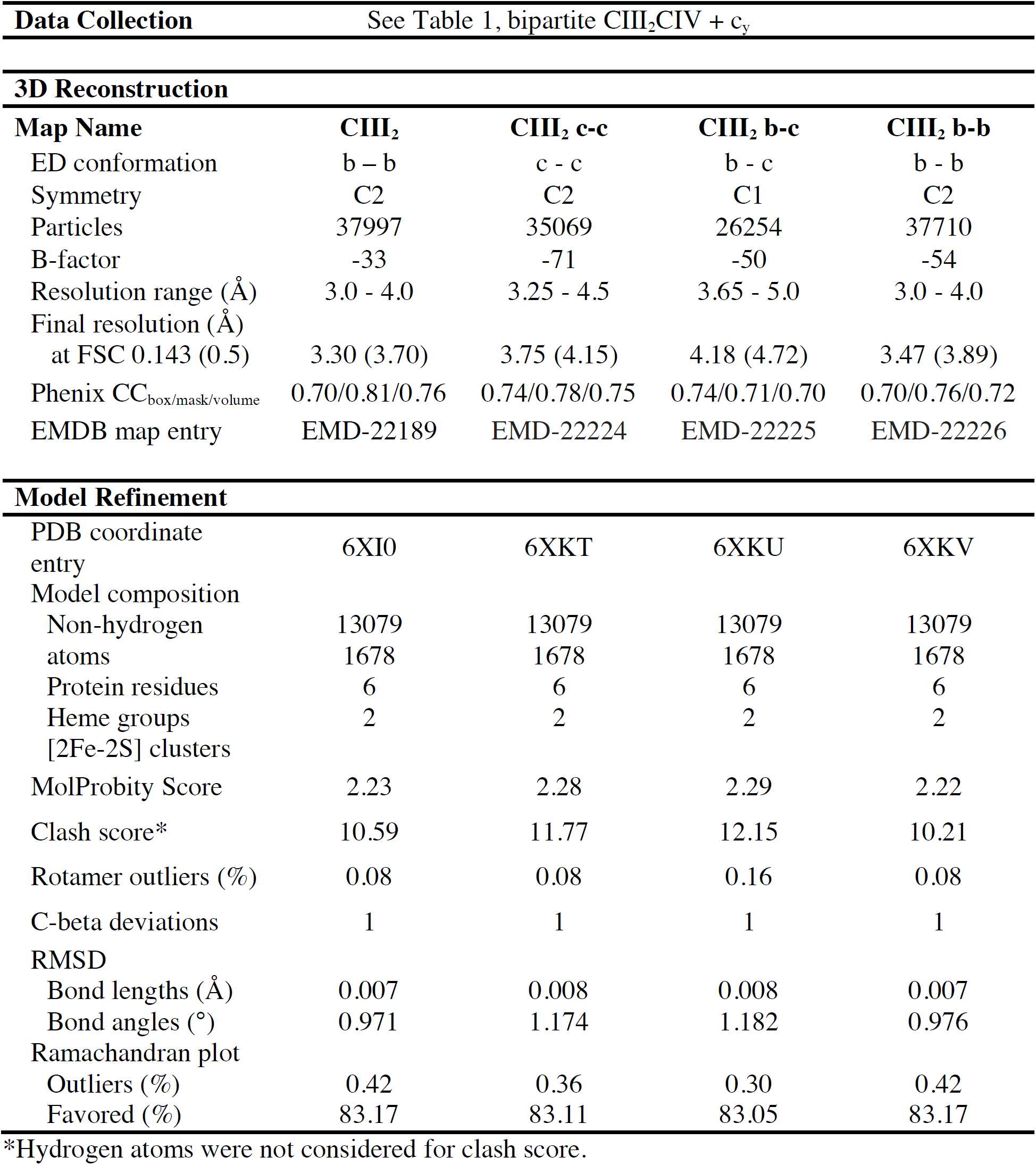
Statistics of 3D reconstruction and model refinement of CIII_2_. The datasets of the bipartite CIII_2_CIV + c_y_ SC (**Table 1**) were used for the 3D reconstruction of CIII_2_. The model (PDB: 6XI0) was refined in map CIII_2_ (EMD-22189) and then used for rigid body fitting in maps CIII_2_ c-c (EMD-22224), CIII_2_ b-c (EMD-22225) and CIII_2_ b-b (EMD-22226).

Superimposition of the CIII_2_ portions of SC-1A and SC-1B maps showed that CIV was in different orientations in different maps (**Fig. S4C**). The two extreme locations of CIV with respect to CIII_2_ were displaced from each other by a translation of ∼3Å and a rotation of ∼37 degrees (**Fig. S4D, E**; SC-1A in red, and SC-1B in blue). Other subclasses identified in 3D classifications showed CIV in slightly different orientations between those seen in SC-1A and SC-2B maps. This variable rotation of CIV around CIII_2_ is attributed to the limited interaction interface between the CcoP (N-ter TMH) of CIV and the cyt *b* (TMH7) of CIII_2_ (see **Fig. 3C**), indicating that the CIII_2_CIV interface is flexible. In the interface regions of SC-1A and SC-1B maps, additional weaker features that are not readily attributable to CIII_2_ and CIV structures were also observed. Intriguingly though, no membrane-external features corresponding to cyt *c* domain of *c*_y_, which is an integral part of the cyt *c*_1_-CcoP-*c*_y_ subunit of tripartite CIII_2_CIV, could be discerned in these maps.

**Figure 3.**
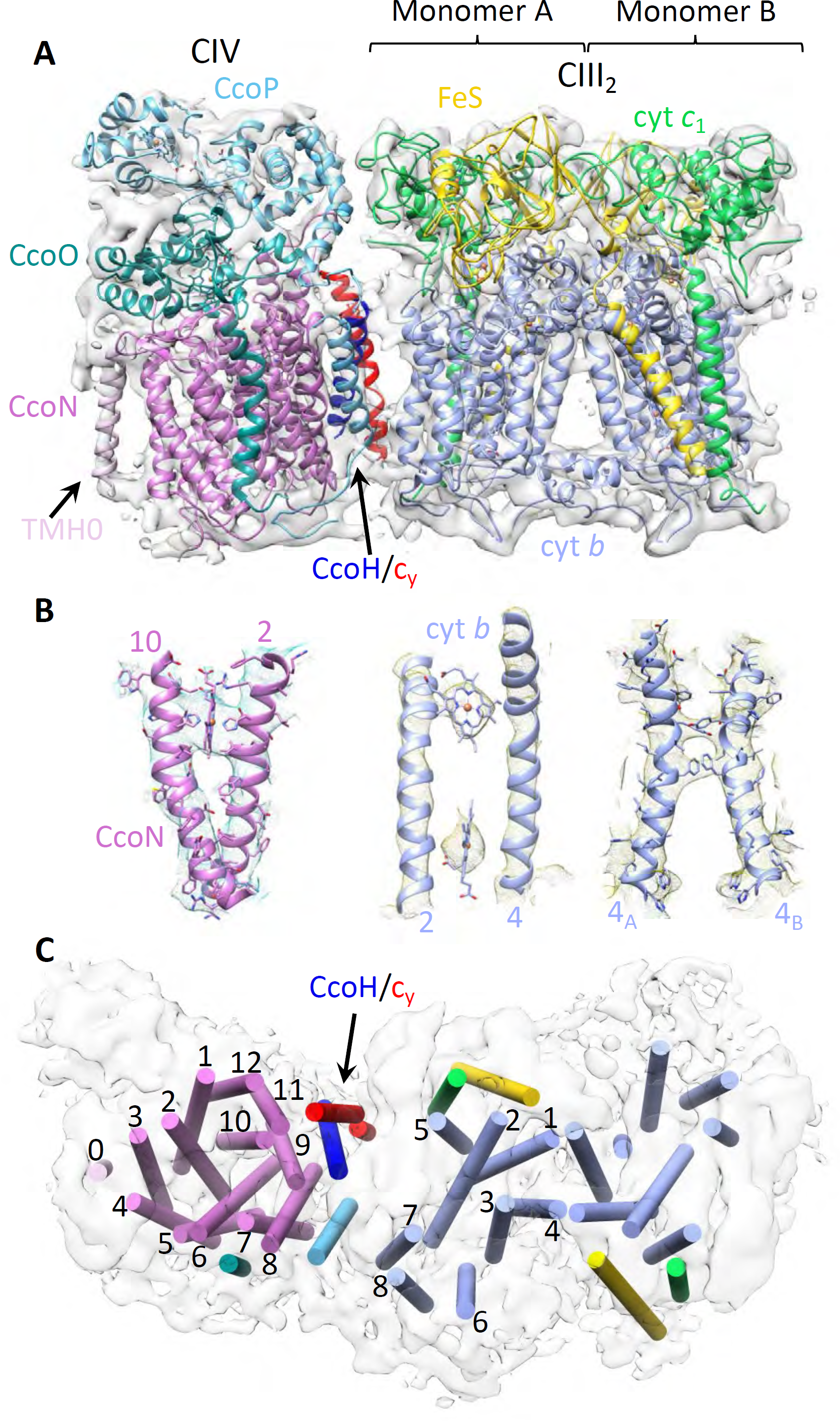
Cryo-EM structure of CIII_2_CIV. **A**. Side view of CIII_2_CIV. The structures of CIII_2_ (PDB: 6XI0, refined in map CIII_2_ (EMD-22189) starting with PDB: 1ZRT), and the homology model of *cbb*_3_-type CIV obtained using *P. stutzeri cbb*_3_ structure (PDB: 3MK7) as a template, were fitted into the cryo-EM map SC-2A depicted in transparent grey. All subunits are colored and labelled as indicated, and the additional feature at the edge of CcoN subunit of CIV, indicated by an arrow, corresponds to the extra N-ter TMH (TMH0, light purple) specific to *R. capsulatus*. The large arrow points out the CcoH/*c*_y_ helices in red/blue. **B.** Representative regions of the cryo-EM map showing the map quality and model fitting. The TMH2 and TMH10 of CcoN (left) shows clearly heme *b* and some bulky side chains. Only the protein backbone and hemes *b*_L_ and *b*_H_ are resolved between the TMH2 and TMH4 of cyt *b* (center) (see **Fig. 5B** for comparison with CIII_2_ map at 3.3 Å). Large side chains are clearly visible between the TMH4 of monomer A and TMH4 of monomer B of CIII_2_ (4_A_ and 4_B_, respectively; right). **C**. Top view of CIII_2_CIV TMHs depicted as cylinders and colored as in **A**. The TMHs of cyt *b* (only CIII_2_ monomer A) and CcoN of CIV are numbered, and the TMHs of the FeS protein (yellow), cyt *c*_1_ (green), CcoO (dark green), CcoP (light blue), CcoH/*c*_y_ (blue/red with an arrow) and CcoN TMH0 (light purple) are shown.

### Structure of bipartite SC supplemented with cyt *c*_y_

In an attempt to locate the cyt *c* domain of *c*_y_, the bipartite SC preparations devoid of it (**Fig. 2A**, fraction A-1) were supplemented with either purified full-length cyt *c*_y_, or with its soluble variant lacking the TMH (*i.e*., cyt S-*c*_y_) (Ozturk et al., 2008), to yield the bipartite SC+*c*_y_ and SC+S-*c*_y_ samples. Following SEC, the elution fractions analyzed by SDS-PAGE showed that only the intact cyt *c*_y_, but not the cyt S-*c*_y_, associated with the SC (**Fig. S5A**). Thus, the cyt *c* domain of *c*_y_ does not bind tightly to, and its TMH is required for association with, this SC.

The cryo-EM analyses of the bipartite SC+*c*_y_ samples were carried out as above, and yielded a map (SC-2A, EMD-22227) at 5.2Å resolution (**Fig. S6A,B**), with local resolutions ranging from 4.3-8.0Å (**Fig. S7A,C**). The homology model of CIV and the refined model of CIII_2_ (PDB: 6XI0) were fitted as rigid bodies into SC-2A with a correlation coefficient CC_box_ of 0.74 (**Fig. 3A**) (**Table 1**). Comparison of SC-2A (bipartite CIII_2_CIV) with SC-1A (tripartite CIII_2_CIV) maps showed that they were highly similar with RMSD of 1.6 Å. They are collectively referred to as CIII_2_CIV, irrespective of their bipartite or tripartite origins.

The dimensions of the slightly curved CIII_2_CIV structure (∼155×60×90Å, LxWxH) were consistent with a CIII_2_ dimer associated with one CIV. On SC-2A map at 5.2Å resolution, some large aromatic side chains could be discerned (**Fig. 3B**), and of the TMHs seen, 34 accounted for by two FeS proteins, two cyts *b* and two cyts *c*_1_ (2, 16 and 2 TMHs per dimer, respectively) of CIII_2_, and single CcoN, CcoO and CcoP (12, 1 and 1 TMHs, respectively) of CIV (**Fig. 3C**). The features corresponding to the heme cofactors of CIII_2_CIV were readily attributed to hemes *b*_H_ and *b*_L_ of cyt *b*, heme *c*_1_ of cyt *c*_1_, and to hemes *b* and *b*_3_ of CcoN, heme *c* of CcoO and hemes *c*_p1_ and *c*_p2_ of CcoP proteins. As seen with the tripartite maps, the [2Fe-2S] clusters of CIII_2_ could be recognized closer to heme *b*_L_ (in b position), but had lower resolution because of conformational heterogeneity.

An additional TMH was observed at the distal end of CIV (**Fig. 3A**, rotated 180 degrees in **Fig. 4A**) close to CcoN TMH3 and TMH4 (**Fig. 3C**). Due to its location, this TMH (depicted in **Fig. 3** and **Fig. 4** as an *ab initio* model of the CcoN Arg25-Leu48 residues generated by I-TASSER (Yang et al., 2015)) was tentatively attributed to the extra N-ter TMH (*i.e*., TMH0) of CcoN (**Fig. 4B**).

**Figure 4.**
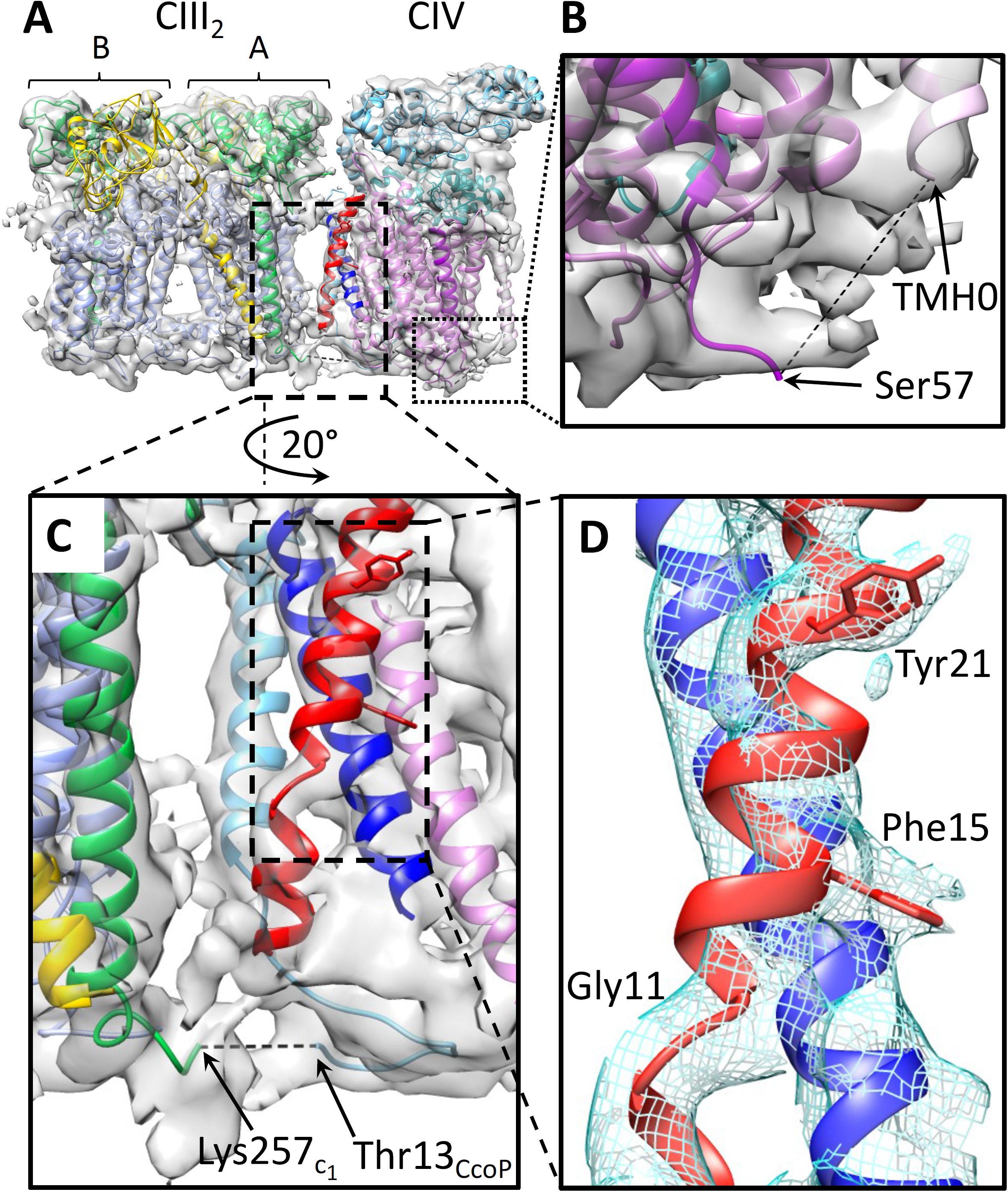
Extra features seen in the cryo-EM map of CIII_2_CIV. **A.** The CIII_2_CIV structure fitted into the map SC-2A (transparent grey) is shown with the same subunit colorings as in **Fig. 3A**, but rotated by 180° for the back view of CIII_2_CIV interface. The two extra TMHs at the interface are attributed to those of CcoH (blue) and cyt *c*_y_ (red). An additional TMH at the edge of CIV is attributed to the predicted N-ter TMH of CcoN (named TMH0, light purple), and depicted as an *ab initio* model generated by I-TASSER server. **B.** Enlarged view of the region linking CcoN TMH1 (dark purple with N-ter Ser57) to the predicted N-ter TMH0 (light purple). The connection between the two TMHs (dashed line) is not resolved. **C.** Enlarged view of CIII_2_CIV interface. The view is slightly rotated relative to **A** for better visibility of CcoP TMH in the background (light blue). For clarity, only CcoN TMH9 is shown next to CcoH (blue) and cyt *c*_y_ (red) TMHs. The fusion region between cyt *c*_1_ and CcoP is shown at the bottom, with the C-ter of cyt *c*_1_ (green) and the N-ter (resolved portion in the map) of CcoP (light blue), and their respective terminal residues (Lys257_c1_ and Thr13_CcoP_) are indicated. The 12 N-terminal CcoP residues connecting these two chains (dashed line) are not clearly resolved. **D.** Enlarged view showing close interaction between the CcoH and cyt *c*_y_ TMHs. Characteristic features of cyt *c*_y_ TMH (NH_2_-**Gly11**xxx**Phe15**xxxxx**Tyr21**-COOH) are used to determine the registration. The helix break induced by Gly11, and the bulky sidechain densities for Phe15 and Tyr21 are clearly visible.

The interface of CIII_2_CIV is roughly delimited by CcoN TMH8 and TMH9, CcoP TMH, cyt *b*-TMH5 and TMH7, and cyt *c*_1_ C-ter TMH of monomer A, with the closest interaction being between CcoP TMH and cyt *b* TMH7 (**Fig. 3A,C**). Two highly confined inter-complex connections and two interacting TMHs of unknown identities were present at the interface (**Fig. 4A**, red and blue TMHs). One such connection was at the *n* face of the membrane, near the cyt *c*_1_ and CcoP TMHs (**Fig. 4C**, Lys257_*c*1_ and Thr13_CcoP_). These subunits being covalently linked, the connecting feature in the map was tentatively attributed to their junction linking CIII_2_ and CIV.

### The assembly factor CcoH and cyt *c*_y_ TMHs are located at CIII_2_CIV interface

The identities of the unknown TMHs at the interface of CIII_2_CIV (**Fig. 3** and **Fig. 4)** were sought using a co-evolution based approach, RaptorX-ComplexContact (Zeng et al., 2018), predicting the residue-residue contacts in protein-protein interactions. All known single TMH containing CIV-related proteins (*i.e*., CcoQ subunit, CcoS and CcoH assembly factors (Koch et al., 2000) and cyt *c*_y_ (Myllykallio et al., 1997)) were analyzed against all subunits of CIII_2_ and CIV. Significant predictions of interacting residue pairs (confidence value >0.5) were observed only between CcoN (primarily TMH9) and the putative CcoH N-term TMH (**Table S4**). An *ab initio* model of CcoH TMH (**Fig. S8A**, residues 11 to 35) was generated by I-TASSER (Yang and Zhang, 2015), and docked onto CIV using PatchDock (Schneidman-Duhovny et al., 2005) with the predicted residue-residue contacts as distance restraints (15 Å threshold) and without using the corresponding cryo-EM maps (SI, Methods). The top scoring models converged to a single cluster around the location of the unknown TMH, close to CcoN TMH9 at CIII_2_CIV interface (**Fig. S8B**). Close examination of the interactions between CcoH TMH and CcoN TMH9 showed that multiple co-evolutionarily conserved residues are in close contacts (**Fig. S8C**). Earlier studies had suggested that CcoH is near the CcoP and CcoN, to which it can be cross-linked by disuccinimidyl suberate (DSS, spacer length ∼11Å) (Pawlik et al., 2010). Thus, the unknown TMH located close to CcoN TMH9 (**Fig. 3C** and **Fig. 4C**, blue TMH) was tentatively assigned to the assembly factor CcoH.

An important difference between the maps of the bipartite CIII_2_CIV+*c*_y_ (SC-2A) and tripartite CIII_2_CIV (SC-1A) was in the features corresponding to the unidentified TMHs at the interface. These densities were barely visible in SC-1A, but highly enhanced in SC-2A (**Fig. 4C**), indicating higher occupancy. The observation that only the native cyt *c*_y_ binds to bipartite SC via its TMH (not its cyt *c* domain, *i.e*., cyt S-*c*_y_), suggested that the TMH (red in **Fig. 4C)**, next to CcoH TMH (blue in **Fig. 4C**), may correspond to the membrane-anchor of cyt *c*_y_. This explanation is most plausible since the bipartite CIII_2_CIV+*c*_y_ samples were supplemented with full-length cyt *c*_y_ while the tripartite samples contained only the fused cyt *c* domain but not the TMH. Indeed, landmark densities corresponding to the helix-breaking Gly11 and two correctly spaced bulky sides chains of Phe15 and Tyr21 of cyt *c*_y_ TMH (NH_2_-xxx**Gly11**xxx**Phe15**xxxxx**Tyr21**-COOH) were discerned (**Fig. 4D**).

Additionally, some CIII_2_CIV+*c*_y_ subclasses exhibited a weak feature on the *p* side of the membrane that may reflect the cyt *c* domain of *c*_y_ (**Fig. S6G**, SC-2B). However, this feature could not be refined to high resolution, consistent with the weak binding of cyt *c* domain of *c*_y_ to CIII_2_CIV (**Fig. S5A**). Moreover, the predominant conformation of CIV in the bipartite CIII_2_CIV+*c*_y_ (**Fig. S6A,B**, SC-2A) shifted towards that found in SC-1A map of tripartite SC (**Fig. S3A**), with no major class corresponding to SC-1B. This suggested that the local interactions between the CcoH and cyt *c*_y_ TMHs and CIV decreased the interface flexibility of CIII_2_CIV (**Fig. 4C**).

### Cryo-EM structures of *R. capsulatus* native CIII_2_

During this study we noted that the bipartite SC+*c*_y_ samples contained large amounts of smaller particles (∼110Å length, **Fig. S3**, Box 2) that were the size of CIII_2_ (**Fig. S6C,D).** Analyses of these particles using C2 symmetry led to the map CIII_2_ at 3.3Å resolution for native CIII_2_ (**Fig. S6E**), with local resolutions ranging from 3.0 to 4.0Å (**Fig. S7B,D**) (**Table 2**). The FeS-ED parts showed a lower occupancy and resolution compared to the rest of the map, indicating conformational heterogeneity. Interestingly, when similar analyses were carried out without imposing C2 symmetry, three distinct maps (CIII_2_ c-c, CIII_2_ b-c and CIII_2_ b-b) for CIII_2_ were obtained at 3.8, 4.2 and 3.5Å resolutions, respectively (**Fig. S6F**). These maps were superimposable with respect to cyt *b* and cyt *c*_1_ subunits, except for the FeS-ED portions. The CIII_2_ structures depicted by the CIII_2_ b-b (**Fig. 5A-C**) and CIII_2_ c-c (**Fig. 5D**) maps exhibited overall C2 symmetry, but in the former the FeS-EDs were located in b, whereas in the latter they were in c position (Esser et al., 2006). Notably, the third structure (**Fig. S6F**, CIII_2_ b-c) was asymmetric, with the FeS-ED of one monomer being in c, and the other in b positions (**Fig. 5E**). Such asymmetric structures of native CIII_2_ have been rarely seen using crystallographic approaches, although proposed to occur during QH_2_ oxidation by CIII_2_ (Castellani et al., 2010; Cooley et al., 2009; Covian and Trumpower, 2005). Similar low occupancy and resolution of the FeS-EDs, suggesting conformational heterogeneity, were also seen with the CIII_2_CIV maps.

**Figure 5:**
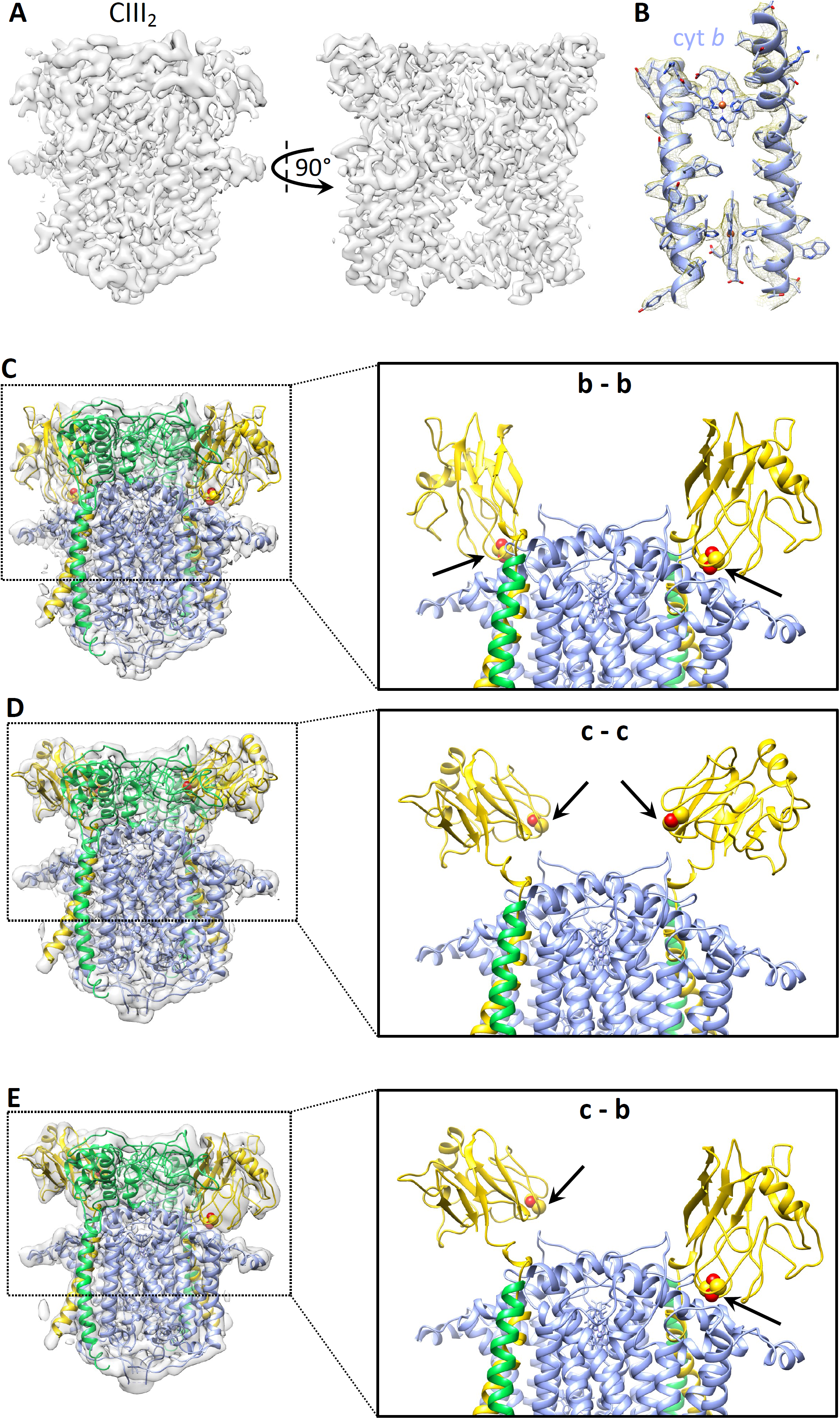
Structures of native CIII_2_ conformers with their FeS proteins in different positions. **A.** Cryo-EM map CIII_2_ b-b with both FeS proteins in b position. **B.** Representative region of map CIII_2_ b-b demonstrating map quality and model fitting. TMH2 and TMH4 of cyt *b* with hemes *b*_L_ and *b*_H_ are shown. **C-E.** Maps and models showing different conformations of the FeS proteins. In each case, the left panels show the CIII_2_ structure fitted into the corresponding maps (**Fig. S6F**) with the subunit colorings (cyt *b* in periwinkle, cyt *c*_1_ in green, and the FeS protein in yellow) as in **Fig. 3**. The right panels depict the top half of the models with the membrane-external domain of cyt *c*_1_ omitted for better visibility of the positions (b - b, c – c and c – b) of FeS-EDs, and the [2Fe-2S] clusters are shown as adjacent yellow-red spheres and indicated by arrows. **C.** Structure of native CIII_2_ with both FeS-EDs in b position (map CIII_2_ b-b, EMD-22226; PDB: 6XKV). **D.** Structure of native CIII_2_ with both FeS-EDs in c position (map CIII_2_ c-c, EMD-22224; PDB: 6XKT). and **E.** Structure of native CIII_2_ with one FeS-ED in c and one in b position (map CIII_2_ b-c, EMD-22225; PDB: 6XKU).

### Interactions of cyt *c*_2_ and cyt *c*_y_ with CIII_2_CIV

The interaction interfaces between CIII_2_CIV and its physiological electron carriers were pursued using cross-linking mass spectrometry (XL-MS) (Gotze et al., 2015; Slavin and Kalisman, 2018). First, the co-crystal structure (PDB: 3CX5) of yeast CIII_2_ with its soluble electron carrier iso-1 cyt *c* (Solmaz and Hunte, 2008) was used as a template (homology between yeast cyt *c*_1_ and *R. capsulatus* cyt *c*_1_: 31% identity and 58% similarity; iso-cyt *c* and cyt *c*_2_: 25% identity and 56% similarity) to model the binding of cyt *c*_2_ on bacterial CIII_2_. As the co-crystal structure contains only one iso-1 cyt *c* bound to one of the two cyt *c*_1_ of yeast CIII_2_, *R. capsulatus* cyt *c*_1_ (PDB: 6XI0) and cyt *c*_2_ (PDB: 1C2N) structures were superimposed with their counterparts on the co-crystal structure, and a model with a single cyt *c*_2_ docked to one monomer of CIII_2_ was generated. To experimentally verify this model, the protein cross-linker 4-(4,6-dimethoxy-1,3,5-triazin-2-yl)-4-methyl-morpholinium chloride (DMTMM) was used with *R. capsulatus* cyt *c*_2_ bound to CIII_2_ (SI, Methods). Multiple intra-subunit cross-links (XLs) within CIII_2_CIV detected in several experiments served as controls (**Table S5** and **Fig. S9A)**. High-confidence XLs were obtained using both FindXL (Kalisman et al., 2012) and MeroX (Iacobucci et al., 2018) search engines, and only those identified by both were retained. The three XLs between cyt *c*_1_ and cyt *c*_2_ provided distance restraints (∼30Å for DMTMM) for docking cyt *c*_2_ to CIII_2_ using PatchDock (**Table S5** and **Fig. S9B**). The docking models clustered at a single region per monomer of CIII_2_ (**Fig. 6A**, right), which overlapped with the binding site of cyt *c*_2_ defined by the model generated by alignment to the yeast co-crystal structure (**Fig. S9C**). The distance from cyt *c*_2_ heme-Fe to cyt *c*_1_ heme-Fe is ∼16.8Å for the co-crystal derived model, while comparable distances between ∼13.8 −20.4Å were obtained with the docking models. Thus, docking with Patchdock integrating XL-MS based distance restraints defined reliably, but with limited accuracy, the interaction region of cyt *c*_2_ on CIII_2_.

**Figure 6:**
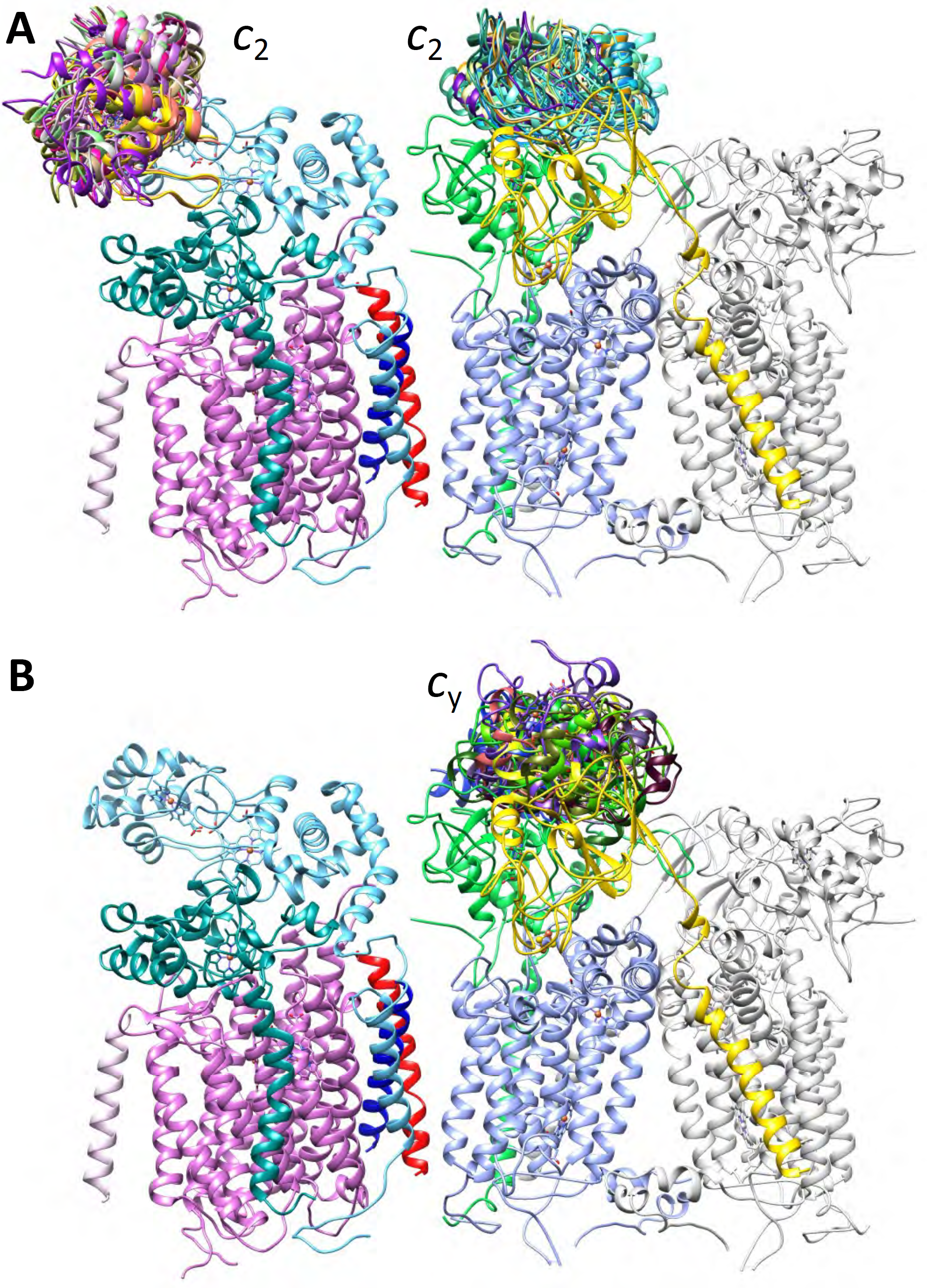
Binding regions of cyt *c*_2_ and cyt *c*_y_ on CIII_2_CIV. The binding regions were defined by XL-MS guided docking, and the subunits of CIII_2_CIV are colored as in **Fig. 3**, except that the monomer B of CIII_2_ is shown in light grey for clarity. Only binding regions on monomer A are shown. **A.** Cyt *c*_2_ (PDB: 1C2N) was docked onto CIII_2_ and CIV using Patchdock with the DMTMM generated XLs as distance restraints, and yielded one cluster of models on CIV and one per monomer of CIII_2_. **B.** A model of cyt *c* domain of *c*_y_, generated using *P. denitrificans* cyt *c*_552_ structure (PDB: 3M97) as a template (RMSD between template and model: 0.2 Å) was docked on CIII_2_ as in **A**, except that both DMTMM and DSBU generated XLs provided distance restraints. Two binding clusters for cyt *c* domain of *c*_y_ per monomer of CIII_2_ were found. These two clusters are located behind each other on a side view, but they are clearly visible on top views (**Fig. S11C**, labelled 1 and 2). Here, only cluster 1 which is closer to cyt *c*_1_ and overlapping with the binding region of cyt *c*_2_ is shown. In all cases, the top 10 representative models are shown to depict the clusters of binding models. No binding region for cyt *c*_y_ on CIV could be defined since no XL was found between these proteins.

No information about the binding sites between cyt *c*_2_ and *cbb*_3_-type CIV was available, so the XL-MS with DMTMM was extended to this case. Similarly, the XLs found between the proteins (1 between cyt *c*_2_ and CcoP, and 8 between cyt *c*_2_ and CcoO) provided distance restraints for docking cyt *c*_2_ to CIV via Patchdock (**Table S5**). The cyt *c*_2_ docking models also clustered in a single region of CIV (**Fig. 6A**, left), closer to heme *c*_p2_ (*c*_2_ heme-Fe to *c*_p2_ heme-Fe: ∼15.2 to 35.6Å) than heme *c*_p1_ (*c*_2_ heme-Fe to *c*_p1_ heme-Fe: ∼23.0 to 42.0Å) of CcoP subunit (**Fig. 7**). Surface charge complementarities between the positively charged face of cyt *c*_2_ and the negatively charged likely binding regions on both CIV and on CIII_2_ are seen (**Fig. S10A**). These two cyt *c*_2_ binding regions on CIII_2_CIV are distant from each other (closest *c*_2_ heme-Fe on CIII_2_ to that on CIV is ∼69Å) (**Fig. 7A**).

**Figure 7.**
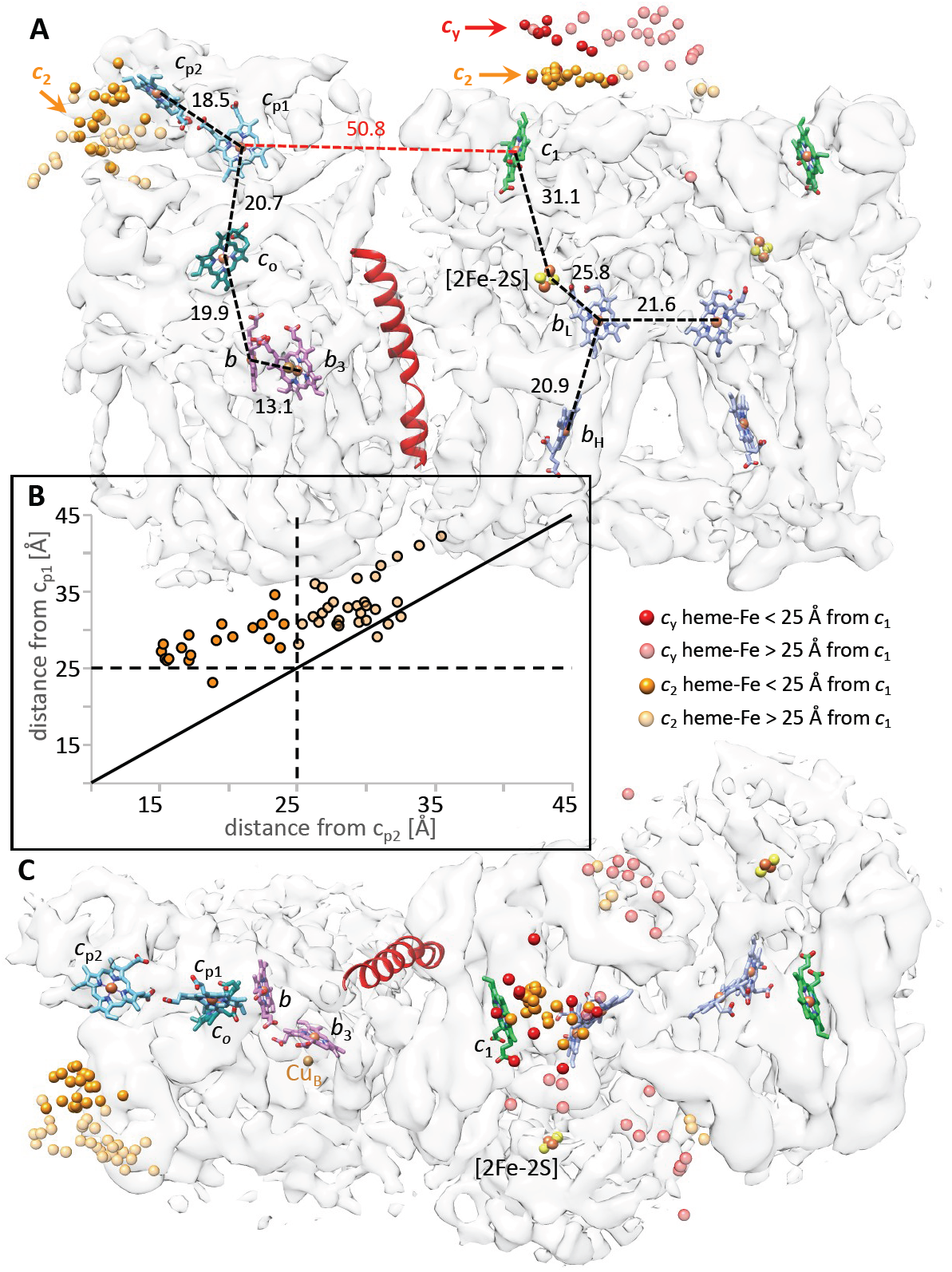
Organization of CIII_2_CIV cofactors and redox partners. **A.** The hemes and [2Fe-2S] clusters are shown inside the transparent cryo-EM map SC-2A of CIII_2_CIV (EMD-22227) with the same subunit colors as in **Fig. 3**: hemes *b*_L_ and *b*_H_ (periwinkle), heme *c*_1_ (green), hemes *c*_p1_ and *c*_p2_ (light blue), heme *c*_o_ (dark green), hemes *b* and *b*_3_ (purple). The [2Fe-2S] clusters are shown as yellow-red spheres. In all cases the distances (heme-Fe to heme-Fe) between the heme cofactors are indicated. The positions of docked cyt *c*_2_ and cyt *c* domain of *c*_y_ are indicated as orange (heme *c*_2_) and red (heme *c*_y_) spheres, respectively, representing their heme-Fe atoms. All heme-Fe atoms corresponding to the top 50 docking positions for cyt *c*_2_ on CIV are shown as solid (< 25Å) or transparent (> 25Å) spheres, depending on their distances to heme *c*_p2_. In the case of CIII_2_, only the docking positions of cyt *c*_2_ and cyt *c*_y_ on monomer A and between the monomers A and B are shown, omitting those located entirely on monomer B. The TMH of cyt *c*_y_ is shown in red at CIII_2_CIV interface. **B.** The heme-Fe atoms of all 50 cyt *c*_2_ models docked onto CIV are plotted in function of their distances from the heme *c*_p1_ and heme *c*_p2_, with the Fe atoms within 25Å shown as solid spheres, and those beyond 25Å as transparent spheres, as indicated. The vast majority of heme-Fe atoms of docked cyt *c*_2_ models are closer to heme *c*_p2_ than heme *c*_p1_ of CIV (above the diagonal line). **C.** Top view of the map shown in **A** is presented to better visualize the distribution of the docked cyt *c* domain of *c*_y_ on monomer A and between the monomers A and B. In all cases, the heme-Fe atoms are depicted by spheres and colored as indicated above and on the figure.

Next, the binding interactions between cyt *c* domain of *c*_y_ and CIII_2_CIV were addressed using DMTMM and disuccinimidyl dibutyric urea (DSBU) as cross-linkers. Similar to DMTMM, DSBU yielded multiple intra-subunit XLs within the subunits of CIII_2_CIV, providing experimental controls (**Table S6** and **Fig. S9D**). Six XLs (five cyt *c*_y_ to cyt *c*_1_ and one cyt *c*_y_ to FeS protein) with DMTMM (**Table S5**) and four XLs (only cyt *c*_y_ to FeS protein) with DSBU (**Table S6**) were identified. Although chemically different cross-linkers were used, XLs were observed only between cyt *c*_y_ and CIII_2_, and not with CIV, suggesting that this cyt *c* domain is closer to CIII_2_ in CIII_2_CIV. Using the XLs as distance restraints (∼35Å for DSBU and ∼30Å for DMTMM) PatchDock generated two binding clusters for cyt *c* domain of *c*_y_ on each CIII_2_ monomer of SC. One of the clusters was on cyt *c*_1_, overlapping with the binding region of cyt *c*_2_ (**Fig. 6B**), whereas the other one was located between cyt *c*_1_ and the FeS-ED near the inter-monomer region of CIII_2_ (**Fig. S11**). To further support these binding locations obtained by XL-MS-based docking, we sought classes that have extra densities corresponding to cyt *c* domain of *c*_y_ in our cryo-EM datasets, and found a minor 3D class containing ∼18,000 particles (**Fig. S6G**), which has an extra feature between CIV and CIII that may be attributable to this domain (**Fig. S11**). The two docking clusters, clearly visible in top view (**Fig. S11C**), were more spread out compared with those of cyt *c*_2_ (**Fig. 6A, Fig. 7A,C**), with the distances between cyt *c*_y_ heme-Fe and cyt *c*_1_ heme-Fe of CIII_2_ monomer A being between 13.8 to 47.1Å, consistent with the weak binding of cyt *c* domain of *c*_y_.

Patchdock mediated docking of cyt *c* domain of *c*_y_ was also performed with the same XLs as above but using the conformers of native CIII_2_ with differently located FeS-EDs (**Fig. 5C-E**, CIII_2_ b-b, c-c and b-c). The data showed that when the FeS-EDs are in c position (CIII_2_ c-c), the docking models gathered as a single cluster on cyt *c*_1_, slightly displaced towards the FeS-ED of the same monomer (**Fig. S12A-C**). However, when the FeS-EDs are in b position (CIII_2_ b-b), such models were more spread out (**Fig. S12D-F**). The third model with one FeS-ED in c and the other in b positions showed the expected clustering pattern depending on the local FeS-ED conformation. As in the SC both FeS-EDs appear to be in the b position, we assume that the docking pattern of cyt *c* domain of *c*_y_ is like that seen with CIII_2_ b-b. Thus, the relatively spread docking position observed with SC (**Fig. 7, Fig. S11**) was attributed to variable conformations of the FeS-EDs on CIII_2_. Furthermore, since heme *c*_1_, and not the FeS protein, is the electron exit site of CIII_2_ (Crofts et al., 2008; Osyczka et al., 2005), the cluster on cyt *c*_1_ was taken as the productive binding region of cyt *c* domain of *c*_y_.

Examination of all pertinent distances between the cofactors of CIII_2_CIV (**Fig. 7A**) indicates that the binding region of cyt *c* domain of *c*_y_ near heme *c*_1_ of CIII_2_ is far away from the expected electron entry point(s) of CIV. The large distance (∼50.8Å) separating cyt *c*_1_ heme-Fe of CIII_2_ monomer A from CcoP *c*_p1_ heme-Fe (the closest compared with heme *c*_p2_ of CIV) renders it impossible to define a location for cyt *c*_*y*_ close enough to heme *c*_1_ reducing it, and heme *c*_p1_ oxidizing it, to sustain productive electron transfer from CIII_2_ to CIV. This distance constraint, the inability to resolve the cyt *c* domain of cyt *c*_y_ by cryo-EM, and the higher frequency of XLs to CIII_2_ strongly infer that the cyt *c* domain of *c*_y_ must oscillate to carry out soluble carrier-independent electron transfer within CIII_2_CIV to couple QH_2_ oxidation to O_2_ reduction (**Fig. 8**).

**Figure 8.**
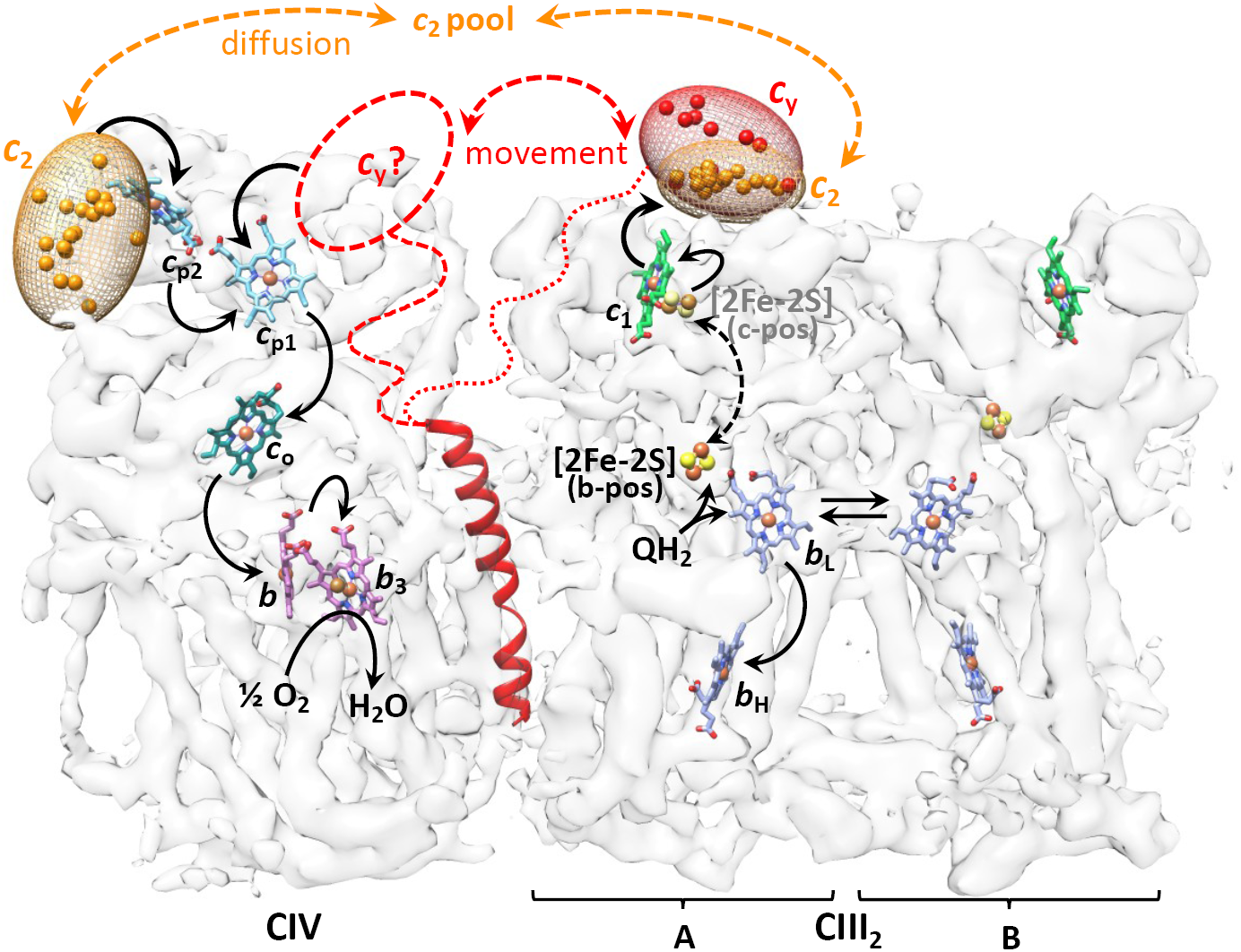
Proposed cyt *c*_2_ and cyt *c*_y_ binding regions of CIII_2_CIV and electron transfer pathways. The likely binding regions of cyt *c*_2_ and cyt *c* domains of *c*_y_ (orange and red ellipsoids, respectively), defined by XL-MS guided docking, are depicted by the distributions of their heme-Fe atoms on the transparent map SC-2A of CIII_2_CIV. Only the positions that are within 25Å of heme *c*_1_ of CIII_2_ or heme *c*_p2_ of CIV are indicated. The CIII_2_ and CIV cofactors together with the TMH of cyt *c*_y_ are shown as in **Fig. 7**. The linker region (indicated by dotted or dashed lines) between the TMH and the cyt *c* domain of *c*_y_ is not resolved in the cryo-EM map. The proposed electron transport pathways are shown by thicker black arrows: upon QH_2_ oxidation by CIII_2_, cyt *c*_y_ which is integral to CIII_2_CIV receives an electron from heme *c*_1_. It then moves (double-headed dashed red arrow) to an undefined binding region (dashed oval with *c*_*y*_?) on CIV, where it delivers the electron to the nearest heme *c*_p1_ of CIV. Similarly, cyt *c*_2_ which is peripheral to CIII_2_CIV also receives an electron from heme *c*_1_, diffuses away to reach CIV and conveys it to heme *c*_p2_. The canonical electron transfers occurring from QH_2_ to heme *c*_1_ in CIII_2_, and from heme *c*_p1_ to O_2_ in CIV, are indicated by thinner arrows. The double headed dashed black arrow depicts the movement of the [2Fe-2S] of FeS protein from the b position (b-pos, in black) to the c position (c-pos, in grey) in CIII_2_ during QH_2_ oxidation. Electron equilibration between the two heme *b*_L_ of CIII_2_ is indicated by double arrows, and the electron transfer steps subsequent to heme *b*_H_ reduction are not shown for the sake of clarity.

## Discussion

Prior to this work, no structural information was available on any bacterial *cbb*_3_-type CIV containing SC, or on its interactions with its physiological redox partners. Here, we describe the first cryo-EM structures of CIII_2_CIV, a *bc*_1_-*cbb*_3_ type respiratory SC from the Gram-negative, facultative phototroph *R. capsulatus*. We also define the likely binding regions of the electron carriers cyt *c*_2_ and cyt *c*_y_ to CIII_2_CIV, and report the structures of both homo- and hetero-dimeric conformers of native CIII_2_. Although X-ray based structures of bacterial *bc*_1_-type CIII_2_ are available, native CIII_2_ heterodimers have not been observed frequently. Similarly, only a single structure, that of *P. stutzeri* (Buschmann et al., 2010), was available for *cbb*_3_-type CIV. Members of this subfamily of heme-Cu:O_2_ reductases are widespread among bacteria and essential for major micro-aerobic processes, including anaerobic photosynthesis, nitrogen fixation, symbiosis and bacterial infection (Khalfaoui-Hassani et al., 2016). Unlike the obligate CIII_2_CIV_2_ SC of *Actinobacteria*, which is rigid and devoid of a free electron carrier (Gong et al., 2018; Wiseman et al., 2018), the *R. capsulatus* facultative CIII_2_CIV is naturally of low abundance and flexible, limiting its structural resolution. The dual function of bacterial CIII_2_ interacting with both the photochemical reaction center in photosynthesis, and cyt *c* oxidase in respiration, may necessitate this natural plasticity to allow swift metabolic adaptations. Similar flexibilities have also been seen with the yeast and human SCs (Sousa and Vonck, 2019).

Isolation of CIII_2_CIV was only possible using a genetically modified strain carrying a translational fusion between CIII_2_ and CIV (SI, Methods). Despite the complexity of translocation, maturation and assembly processes of multi-cofactor containing membrane complexes, this fusion approach is of general use. Our fused SC preparations were compositionally heterogeneous, containing mixtures of CIII_2_CIV_2_, CIII_2_CIV and CIII_2_ particles. The basis of this heterogeneity is unclear, though it may stem from subunit sub-stoichiometry, incomplete assembly, or higher susceptibility to degradation during sample preparations. Insertion of different spacers at the cyt *c*_1_-CcoP fusion junction, overexpression of the subunits and the related assembly components could not overcome the heterogeneity (SI, Methods). Consequently, structural studies required extensive data collections and limited structural resolutions, but allowed analyses of fragmented particles.

### Structures of CIII_2_CIV

The structures of the tripartite CIII_2_CIV or bipartite CIII_2_CIV+*c*_y_ at sub-nanometer resolution (∼5.2 to 7.2Å) were highly similar. Limited protein-protein interaction between the subunits of CIII_2_ and CIV was seen at the interface where the TMHs of cyt *c*_y_ and CcoH were located (**Fig. 4**), limiting the flexibility of CIII_2_CIV. Another helix-like feature found at the exterior edge of CIV was attributed to the extra N-ter helix (TMH0) unique to *R. capsulatus* CcoN. However, due to the limited resolutions of the structures, these attributions are tentative. Limited resolution also precluded identification of non-protein constituents at the CIII_2_CIV interface. In this respect, *R. capsulatus* lacks cardiolipin, often implicated in SC stability (Arias-Cartin et al., 2012). Instead, it produces ornithine lipid that can mimic cardiolipin upon dimerization (Aygun-Sunar et al., 2006). Ornithine lipid-less mutants contain very low amounts of CIV and CIII_2_, and if any SC is unknown.

Previously, neither the exact location nor the mobility of cyt *c*_y_, which is the basis of the “soluble carrier-independent” electron transfer from CIII_2_ to CIV, were known. The SC structure shows that locking the N-terminal TMH of cyt *c*_y_ at the interface allows mobility of its cyt *c* domain (**Fig. 8**). The linker region attaching the TMH to cyt *c* domain remains unresolved, but is long enough to allow oscillations between CIII_2_ and CIV. Earlier studies with *R. capsulatus* cyt *c*_y_ had shown that a full-length linker is needed for rapid (< ∼50 μsec) electron transfer from CIII_2_ to the photosynthetic reaction center in photosynthesis (Myllykallio et al., 1998). In contrast, a shorter linker (∼45-residue instead of 69) is fully proficient for respiratory electron transfer to CIV (Daldal et al., 2001).

### Structures of bacterial native CIII_2_

In native CIII_2_ conformers, different positions of the [2Fe-2S] cluster bearing FeS-EDs were seen. Crystallographic structures have often depicted bacterial CIII_2_ as symmetrical homodimers (Berry et al., 2004; Esser et al., 2006; Xia et al., 2008). These structures were obtained in the presence of inhibitors constraining FeS-EDs near heme *b*_L_ or used mutants stabilizing it on cyt *b* surface. Alternatively, they contained crystal contacts restricting the FeS-ED movement (Esser et al., 2008). To our knowledge, no native heterodimeric CIII_2_ structure of bacterial origin with different conformations of its FeS-EDs has been reported. Only recently, the cryo-EM structures of mitochondrial SCs with different maps for CIII_2_ FeS-EDs have been reported (Letts et al., 2019; Sousa et al., 2016). Thus, native CIII_2_ is not always a symmetric homodimer, and the FeS-ED of each monomer is free to move independently from each other, which has functional implications. The Q-cycle models describe the mechanism of CIII_2_ catalysis by two turnovers of a given monomer (Crofts and Berry, 1998; Crofts et al., 2008; Osyczka et al., 2005). The mobility of the FeS-ED between the b and c positions is essential for QH_2_ oxidation, and the different positions of the FeS-ED protein are often attributed to different catalytic steps (Esser et al., 2006). Emerging asymmetric structures of bacterial and mitochondrial native CIII_2_ obtained by cryo-EM in the absence of inhibitors or mutations, combined with the well-established inter-monomer electron transfer between the heme *b*_L_ of the monomers (Lanciano et al., 2013; Lanciano et al., 2011; Swierczek et al., 2010; Yu et al., 2002), start to provide a glimpse into plausible “heterodimeric Q cycle” mechanism(s) (Castellani et al., 2010; Cooley, 2010; Cooley et al., 2009), at least when CIII_2_ is a part of SCs. Accordingly, CIII_2_ may cycle between homo- and hetero-dimeric conformations in regards to its FeS-EDs during catalysis. These mechanistic implications remain to be studied.

### Electronic communication between CIII_2_CIV partners

Earlier, binding interactions between CIII_2_CIV and its physiological electron carriers were unknown. Here we defined the likely interaction regions between the cyt *c*_2_ or the cyt *c*_y_ and CIII_2_CIV (**Fig. 8)**. The CIII_2_CIV structure indicates that the distances separating heme *c*_1_ of CIII_2_ monomer **A** and hemes *c*_p1_ and *c*_p2_ of CIV are too large (**Fig. 7**) for direct microsecond scale electronic communication (Moser et al., 1992) to sustain the turnover rate of CIII_2_CIV. Thus, even when CIII_2_ and CIV form a SC, a freely diffusing cyt *c*_2_ or a membrane-anchored mobile cyt *c*_y_, is required for QH_2_:O_2_ oxidation.

The binding region of cyt *c*_2_ on CIII_2_ was identified earlier (Solmaz and Hunte, 2008), but that on CIV was unknown. The binding location of cyt *c*_2_ on CIV determined in this study, the redox midpoint potentials (E_m_) of the cofactors and the distances separating them (**Fig. 7A**) suggest that cyt *c*_2_ would confer electrons to the closer heme *c*_p2_, rather than the more distant heme *c*_p1_, of CcoP. This will then initiate canonical electron transfer via heme *c*_p1_, heme *c*_o_ and heme *b* to heme *b*_3_-Cu_B_ site for O_2_ reduction (Brzezinski and Gennis, 2008; Wikstrom et al., 2018) (**Fig. 8**). For purified *R. capsulatus* proteins, the E_m_ value of cyt *c*_2_ is ∼350 mV (Myllykallio et al., 1999), while those of CIII_2_ heme *c*_1_ and CIV heme *c*_o_ are ∼320 mV (Valkova-Valchanova et al., 1998) and ∼210 mV (Gray et al., 1994), respectively. The E_m_ values of *R. capsulatus* CIV hemes *c*_p1_ and *c*_p2_ are unknown, but based on similar E_m_ values of heme *c*_o_ for *B. japonicum* (200 mV) and *R. capsulatus* (210 mV), they are expected to be close to those of *B. japonicum c*_p1_ (∼300 mV) and *c*_p2_ (∼390 mV) (Verissimo et al., 2007).

In the case of cyt *c*_y_, its interaction region on CIV remains less well defined. Of the two binding regions of cyt *c*_y_ on CIII_2_, that on cyt *c*_1_ was taken as the most likely functional site. This binding region on cyt *c*_1_ is close to that of cyt *c*_2_, but cyt *c*_y_ has less complementary surface charges (**Fig. S10B**), consistent with its weaker binding to CIII_2_CIV. Anchoring cyt *c*_y_ to the membrane, next to its redox partners, might have enhanced its electron transfer efficiency while minimizing its electrostatic interactions with its partners.

The distance separating the redox centers is a major factor that controls the rate of electron transfer (Moser et al., 1992). The binding region of cyt *c* domain of *c*_y_ on CIII_2_ suggests that reduced cyt *c*_y_, upon its movement to CIV, might preferentially convey electrons to the closer heme *c*_p1_ than heme *c*_p2_ of CcoP (**Fig. 8**). If so, then under physiological conditions, heme *c*_p1_ would be the primary receiver of electrons derived from QH_2_ oxidation by CIII_2_, forming a fully membrane-confined electronic wiring within CIII_2_CIV. In contrast, cyt *c*_2_ carries electrons from heme *c*_1_ to heme *c*_p2_ via free diffusion. Significantly, this membrane-external pathway might accommodate electrons not only from QH_2_ but also from other donors distinct from CIII_2_. As such, reduction of cyt *c*_2_ during methylamine oxidation (Otten et al., 2001), or degradation of sulfur containing amino acids, converting toxic sulfite (SO_3_ ^2-^) to sulfate (SO_4_ ^-2^) by sulfate oxidase (Kappler and Dahl, 2001) might provide electrons to CIV, contributing to cellular energy production.

In summary, for the first time, the architecture of CIII_2_CIV SC along with its dynamics and interactions with its physiological redox partners established salient structural features of two distinct respiratory electron transport pathways (membrane-confined and membrane-external) that operate between CIII_2_ and CIV in Gram-negative bacteria.

## Abbreviations

Q: quinone
QH_2_: Quinol or hydroquinone
Complex III: CIII_2_ or cytochrome
*bc*_1_: ubiquinolcytochrome *c* oxidoreductase
Complex IV or CIV: *cbb*_3_-type cytochrome *c* oxidase
cyt: cytochrome
cyt *c*_2_: cytochrome *c*_2_, soluble cytochrome *c*
cyt *c*_y_: cytochrome *c*_y_, membrane-anchored cytochrome *c*
cyt S-*c*_y_: soluble part of cytochrome *c*_y_ without its membrane anchor
SC: super-complex
MS: mass spectrometry
XL-MS: cross-linking mass spectrometry
XL: cross-links
TMBZ: 3,3’,5,5’-tetramethyl-benzidine
DBH_2_: 2,3-dimethoxy-5-methyl-6-*decyl*-1,4-*benzoquinone*
FeS: Rieske iron-sulfur protein
FeS-ED: membrane-extrinsic domain of FeS protein
b position: location of the [2Fe-2S] cluster near heme *b*_L_
c position: location of the [2Fe-2S] cluster near heme *c*_1_
cryo-EM: cryogenic electron microscopy
BN-PAGE: blue native polyacrylamide gel electrophoresis
SDS-PAGE: sodium dodecylsulfate polyacrylamide gel electrophoresis
C-ter: C-terminus
N-ter: N-terminus
His-tag: 8-histidine tag
FLAG-tag: DYKDDDDK-tag
SEC: size exclusion chromatography
TMH: transmembrane helix
DSS: disuccinimidyl suberate
DSBU: disuccinimidyl dibutyric urea; DMTMM, 4-(4,6-dimethoxy-1,3,5-triazin-2-yl)-4-methyl-morpholinium chloride
heme-Fe: heme-iron
E_m_: redox midpoint potential
heme *c*_p1_: N-ter located *c*-type heme 1 of CcoP
heme *c*_p2_: C-ter located *c*-type heme 2 of CcoP
heme *c*_o_: *c*-type heme of CcoO
SO_3_ ^2-^: sulfite
SO_4_ ^2-^: sulfate
RMSD: root-mean-square deviation
DDM: n-dodecyl β-D-maltoside.

## Acknowledgments

This work was supported partly by the National Institute of Health grants, GM 38237 to FD, GM123233 to KM, GM110174 and AI118891 to BAG, T32-GM008275 to TV, T32-GM071339 to HJK, and partly by the Division of Chemical Sciences, Geosciences and Biosciences, Office of Basic Energy Sciences of Department of Energy grant DE-FG02-91ER20052 to FD. This research was funded by ISF 1466/18, BSF 2016070, and Ministry of Science and technology 80802 grants to DS. YO was supported by the grant GRK2202-23577276/RTG from DFG, Germany. Data analysis was partly supported by the National Institute of Health grant S10OD023592.

We thank Drs. Saif S. Hasan, Brian G. Pierce and Christian Presley at the Institute for Bioscience and Biotechnology Research (IBBR), University of Maryland, for insightful discussions and invaluable help they provided during this study. SS and FD also thank Vivian Kitainda for her assistance with protein purification and O_2_ consumption measurements.

This research was, in part, supported by the National Cancer Institute, National Cryo-EM Facility at the Frederick National Laboratory for Cancer Research under contract HSSN261200800001E. The authors would like to thank Ulrich Baxa, Thomas Edwards and Adam Wier for their support and helpful discussions. Some cryo-EM data were also obtained at the University of Massachusetts Cryo-EM Core Facility, and we thank Drs. Chen Xu, KangKang Song and Kyounghwan Lee for their support. Cryo-EM sample screening and optimization was performed at the Electron Microscopy Resource Laboratory at the Perelman School of Medicine, University of Pennsylvania, and we thank Dr. Sudheer Molugu for his support.

## Data deposition

The following *R. capsulatus* structures and the corresponding cryo-EM maps are deposited to PDB and EMDB with the accession codes listed in the table below:

| Structure | PDB | map |
| --- | --- | --- |
| CIII<sub>2</sub> | 6XI0 | EMD-22189 |
| CIII<sub>2</sub> c-c | 6XKT | EMD-22224 |
| CIII<sub>2</sub> b-c | 6XKU | EMD-22225 |
| CIII<sub>2</sub> b-b | 6XKV | EMD-22226 |
| SC-2A | 6XKW | EMD-22227 |
| SC-1A | 6XKX | EMD-22228 |
| SC-1B | 6XKZ | EMD-22230 |

The raw XL-MS data deposited to PRIDE repository (http://www.ebi.ac.uk/pride/archive/) with the dataset identifier PXD020038

## References

Acin-Perez, R., and Enriquez, J.A. (2014). The function of the respiratory supercomplexes: the plasticity model. Biochim Biophys Acta 1837, 444–450.

Albert, I., Rutherford, A.W., Grav, H., Kellermann, J., and Michel, H. (1998). The 18 kDa cytochrome *c*_553_ from *Heliobacterium gestii*: gene sequence and characterization of the mature protein. Biochemistry 37, 9001–9008.

Arias-Cartin, R., Grimaldi, S., Arnoux, P., Guigliarelli, B., and Magalon, A. (2012). Cardiolipin binding in bacterial respiratory complexes: structural and functional implications. Biochim Biophys Acta 1817, 1937–1949.

Aspholm, M., Aas, F.E., Harrison, O.B., Quinn, D., Vik, A., Viburiene, R., Tonjum, T., Moir, J., Maiden, M.C., and Koomey, M. (2010). Structural alterations in a component of cytochrome *c* oxidase and molecular evolution of pathogenic *Neisseria* in humans. PLoS Pathog 6, e1001055.

Aygun-Sunar, S., Mandaci, S., Koch, H.G., Murray, I.V., Goldfine, H., and Daldal, F. (2006). Ornithine lipid is required for optimal steady-state amounts of *c*-type cytochromes in *Rhodobacter capsulatus*. Mol Microbiol 61, 418–435.

Bergdoll, L., Brink, F., Nitschke, W., Picot, D., and Baymann, F. (2016). From low-to high-potential bioenergetic chains: Thermodynamic constraints of Q-cycle function. Biochim Biophys Acta 1857, 1569–1579.

Berry, E.A., Huang, L.S., Saechao, L.K., Pon, N.G., Valkova-Valchanova, M., and Daldal, F. (2004). X-Ray Structure of *Rhodobacter capsulatus* Cytochrome *bc*_1_: Comparison with its Mitochondrial and Chloroplast Counterparts. Photosynth Res 81, 251–275.

Berry, E.A., and Trumpower, B.L. (1985). Isolation of ubiquinol oxidase from *Paracoccus denitrificans* and resolution into cytochrome *bc*_1_ and cytochrome *c*-*aa*_3_ complexes. J Biol Chem 260, 2458–2467.

Bott, M., Ritz, D., and Hennecke, H. (1991). The *Bradyrhizobium japonicum cycM* gene encodes a membrane-anchored homolog of mitochondrial cytochrome *c*. J Bacteriol 173, 6766–6772.

Brzezinski, P. (2019). New Structures Reveal Interaction Dynamics in Respiratory Supercomplexes. Trends Biochem Sci.

Brzezinski, P., and Gennis, R.B. (2008). Cytochrome *c* oxidase: exciting progress and remaining mysteries. J Bioenerg Biomembr 40, 521–531.

Buschmann, S., Warkentin, E., Xie, H., Langer, J.D., Ermler, U., and Michel, H. (2010). The structure of *cbb*_3_ cytochrome oxidase provides insights into proton pumping. Science 329, 327–330.

Castellani, M., Covian, R., Kleinschroth, T., Anderka, O., Ludwig, B., and Trumpower, B.L. (2010). Direct demonstration of half-of-the-sites reactivity in the dimeric cytochrome bc1 complex: enzyme with one inactive monomer is fully active but unable to activate the second ubiquinol oxidation site in response to ligand binding at the ubiquinone reduction site. J Biol Chem 285, 502–510.

Cooley, J.W. (2010). A structural model for across membrane coupling between the Q_o_ and Q_i_ active sites of cytochrome *bc*_1_. Biochim Biophys Acta 1797, 1842–1848.

Cooley, J.W., Lee, D.W., and Daldal, F. (2009). Across membrane communication between the Q_o_ and Q_i_ active sites of cytochrome *bc*_1_. Biochemistry 48, 1888–1899.

Covian, R., and Trumpower, B.L. (2005). Rapid electron transfer between monomers when the cytochrome *bc*_1_ complex dimer is reduced through center N. J Biol Chem 280, 22732–22740.

Crofts, A.R., and Berry, E.A. (1998). Structure and function of the cytochrome *bc*_1_ complex of mitochondria and photosynthetic bacteria. Curr Opin Struct Biol 8, 501–509.

Crofts, A.R., Holland, J.T., Victoria, D., Kolling, D.R., Dikanov, S.A., Gilbreth, R., Lhee, S., Kuras, R., and Kuras, M.G. (2008). The Q-cycle reviewed: How well does a monomeric mechanism of the bc(1) complex account for the function of a dimeric complex? Biochim Biophys Acta 1777, 1001–1019.

Daldal, F., Cheng, S., Applebaum, J., Davidson, E., and Prince, R.C. (1986). Cytochrome *c*_2_ is not essential for photosynthetic growth of *Rhodopseudomonas capsulata*. Proc Natl Acad Sci U S A 83, 2012–2016.

Daldal, F., Mandaci, S., Winterstein, C., Myllykallio, H., Duyck, K., and Zannoni, D. (2001). Mobile cytochrome *c*_2_ and membrane-anchored cytochrome *c*_y_ are both efficient electron donors to the *cbb*_3_- and *aa*_3_-type cytochrome *c* oxidases during respiratory growth of *Rhodobacter sphaeroides*. J Bacteriol 183, 2013–2024.

Darrouzet, E., Moser, C.C., Dutton, P.L., and Daldal, F. (2001). Large scale domain movement in cytochrome *bc*_1_: a new device for electron transfer in proteins. Trends Biochem Sci 26, 445–451.

Davidson, E., Ohnishi, T., Tokito, M., and Daldal, F. (1992). *Rhodobacter capsulatus* mutants lacking the Rieske FeS protein form a stable cytochrome *bc*_1_ subcomplex with an intact quinone reduction site. Biochemistry 31, 3351–3358.

Ducluzeau, A.L., Ouchane, S., and Nitschke, W. (2008). The *cbb*_3_ oxidases are an ancient innovation of the domain bacteria. Mol Biol Evol 25, 1158–1166.

Enriquez, J.A. (2016). Supramolecular Organization of Respiratory Complexes. Annu Rev Physiol 78, 533–561.

Esser, L., Elberry, M., Zhou, F., Yu, C.A., Yu, L., and Xia, D. (2008). Inhibitor-complexed structures of the cytochrome *bc*_1_ from the photosynthetic bacterium *Rhodobacter sphaeroides*. J Biol Chem 283, 2846–2857.

Esser, L., Gong, X., Yang, S., Yu, L., Yu, C.A., and Xia, D. (2006). Surface-modulated motion switch: capture and release of iron-sulfur protein in the cytochrome *bc*_1_ complex. Proc Natl Acad Sci U S A 103, 13045–13050.

Gong, H., Li, J., Xu, A., Tang, Y., Ji, W., Gao, R., Wang, S., Yu, L., Tian, C., Li, J., et al. (2018). An electron transfer path connects subunits of a mycobacterial respiratory supercomplex. Science 362.

Gotze, M., Pettelkau, J., Fritzsche, R., Ihling, C.H., Schafer, M., and Sinz, A. (2015). Automated assignment of MS/MS cleavable cross-links in protein 3D-structure analysis. J Am Soc Mass Spectrom 26, 83–97.

Gray, K.A., Grooms, M., Myllykallio, H., Moomaw, C., Slaughter, C., and Daldal, F. (1994). *Rhodobacter capsulatus* contains a novel *cb*-type cytochrome *c* oxidase without a Cu_A_ center. Biochemistry 33, 3120–3127.

Gu, J., Wu, M., Guo, R., Yan, K., Lei, J., Gao, N., and Yang, M. (2016). The architecture of the mammalian respirasome. Nature 537, 639–643.

Hartley, A.M., Lukoyanova, N., Zhang, Y., Cabrera-Orefice, A., Arnold, S., Meunier, B., Pinotsis, N., and Marechal, A. (2019). Structure of yeast cytochrome *c* oxidase in a supercomplex with cytochrome *bc*_1_. Nat Struct Mol Biol 26, 78–83.

Hochkoeppler, A., Jenney, F.E., Jr., Lang, S.E., Zannoni, D., and Daldal, F. (1995). Membrane-associated cytochrome *c*_*y*_ of *Rhodobacter capsulatus* is an electron carrier from the cytochrome *bc*_1_ complex to the cytochrome *c* oxidase during respiration. J Bacteriol 177, 608–613.

Iacobucci, C., Gotze, M., Ihling, C.H., Piotrowski, C., Arlt, C., Schafer, M., Hage, C., Schmidt, R., and Sinz, A. (2018). A cross-linking/mass spectrometry workflow based on MS-cleavable cross-linkers and the MeroX software for studying protein structures and protein-protein interactions. Nat Protoc 13, 2864–2889.

Jenney, F.E., Jr., and Daldal, F. (1993). A novel membrane-associated *c*-type cytochrome, cyt *c*_y_, can mediate the photosynthetic growth of *Rhodobacter capsulatus* and *Rhodobacter sphaeroides*. EMBO J 12, 1283–1292.

Kalisman, N., Adams, C.M., and Levitt, M. (2012). Subunit order of eukaryotic TRiC/CCT chaperonin by cross-linking, mass spectrometry, and combinatorial homology modeling. Proc Natl Acad Sci U S A 109, 2884–2889.

Kao, W.C., Kleinschroth, T., Nitschke, W., Baymann, F., Neehaul, Y., Hellwig, P., Richers, S., Vonck, J., Bott, M., and Hunte, C. (2016). The obligate respiratory supercomplex from *Actinobacteria*. Biochim Biophys Acta 1857, 1705–1714.

Kappler, U., and Dahl, C. (2001). Enzymology and molecular biology of prokaryotic sulfite oxidation. FEMS Microbiol Lett 203, 1–9.

Khalfaoui-Hassani, B., Verissimo, A.F., Shroff, N., Ekici, S., Trasnea, P.-I., Utz, M., Koch, H.-G., and Daldal, F. (2016). Biogenesis of cytochrome *c* complexes: from insertion of redox cofactors to assembly of different subunits. In Cytochrome Complexes: Evolution, Structures, Energy Transduction, and Signaling, W. Cramer, and T. Kallas, eds. (Dordrecht: Springer), pp. 527–555.

Kim, M.S., Jang, J., Ab Rahman, N.B., Pethe, K., Berry, E.A., and Huang, L.S. (2015). Isolation and Characterization of a Hybrid Respiratory Supercomplex Consisting of *Mycobacterium tuberculosis* Cytochrome *bcc* and *Mycobacterium smegmatis* Cytochrome *aa*_3_. J Biol Chem 290, 14350–14360.

Koch, H.G., Winterstein, C., Saribas, A.S., Alben, J.O., and Daldal, F. (2000). Roles of the *ccoGHIS* gene products in the biogenesis of the *cbb*_3_-type cytochrome *c* oxidase. J Mol Biol 297, 49–65.

Kulajta, C., Thumfart, J.O., Haid, S., Daldal, F., and Koch, H.G. (2006). Multi-step assembly pathway of the *cbb*_3_-type cytochrome *c* oxidase complex. J Mol Biol 355, 989–1004.

Lanciano, P., Khalfaoui-Hassani, B., Selamoglu, N., and Daldal, F. (2013). Intermonomer electron transfer between the *b* hemes of heterodimeric cytochrome *bc*_1_. Biochemistry 52, 7196–7206.

Lanciano, P., Lee, D.W., Yang, H., Darrouzet, E., and Daldal, F. (2011). Intermonomer electron transfer between the low-potential *b* hemes of cytochrome *bc*_1_. Biochemistry 50, 1651–1663.

Lee, D.W., Ozturk, Y., Osyczka, A., Cooley, J.W., and Daldal, F. (2008). Cytochrome *bc*_1_-*c*_y_ fusion complexes reveal the distance constraints for functional electron transfer between photosynthesis components. J Biol Chem 283, 13973–13982.

Letts, J.A., Fiedorczuk, K., Degliesposti, G., Skehel, M., and Sazanov, L.A. (2019). Structures of Respiratory Supercomplex I+III2 Reveal Functional and Conformational Crosstalk. Mol Cell 75, 1131–1146 e1136.

Letts, J.A., Fiedorczuk, K., and Sazanov, L.A. (2016). The architecture of respiratory supercomplexes. Nature 537, 644–648.

Letts, J.A., and Sazanov, L.A. (2017). Clarifying the supercomplex: the higher-order organization of the mitochondrial electron transport chain. Nat Struct Mol Biol 24, 800–808.

Melo, A.M., and Teixeira, M. (2016). Supramolecular organization of bacterial aerobic respiratory chains: From cells and back. Biochim Biophys Acta 1857, 190–197.

Milenkovic, D., Blaza, J.N., Larsson, N.G., and Hirst, J. (2017). The Enigma of the Respiratory Chain Supercomplex. Cell Metab 25, 765–776.

Moser, C.C., Keske, J.M., Warncke, K., Farid, R.S., and Dutton, P.L. (1992). Nature of biological electron transfer. Nature 355, 796–802.

Myllykallio, H., Drepper, F., Mathis, P., and Daldal, F. (1998). Membrane-anchored cytochrome *c*_y_ mediated microsecond time range electron transfer from the cytochrome *bc*_1_ complex to the reaction center in *Rhodobacter capsulatus*. Biochemistry 37, 5501–5510.

Myllykallio, H., Drepper, F., Mathis, P., and Daldal, F. (2000). Electron-transfer supercomplexes in photosynthesis and respiration. Trends Microbiol 8, 493–494.

Myllykallio, H., Jenney, F.E., Jr., Moomaw, C.R., Slaughter, C.A., and Daldal, F. (1997). Cytochrome *c*_y_ of *Rhodobacter capsulatus* is attached to the cytoplasmic membrane by an uncleaved signal sequence-like anchor. J Bacteriol 179, 2623–2631.

Myllykallio, H., Zannoni, D., and Daldal, F. (1999). The membrane-attached electron carrier cytochrome *c*_y_ from *Rhodobacter sphaeroides* is functional in respiratory but not in photosynthetic electron transfer. Proc Natl Acad Sci U S A 96, 4348–4353.

Nicholls, D.G., and Ferguson, S.J. (2013). Bioenergetics 4 (Elsevier).

Niebisch, A., and Bott, M. (2003). Purification of a cytochrome *bc*-*aa*_3_ supercomplex with quinol oxidase activity from *Corynebacterium glutamicum*. Identification of a fourth subunity of cytochrome *aa*_3_ oxidase and mutational analysis of diheme cytochrome *c*_1_. J Biol Chem 278, 4339–4346.

Osyczka, A., Moser, C.C., and Dutton, P.L. (2005). Fixing the Q cycle. Trends Biochem Sci 30, 176–182.

Otten, M.F., van der Oost, J., Reijnders, W.N., Westerhoff, H.V., Ludwig, B., and Van Spanning, R.J. (2001). Cytochromes *c*_550_, *c*_552_, and *c*_1_ in the electron transport network of *Paracoccus denitrificans*: redundant or subtly different in function? J Bacteriol 183, 7017–7026.

Ozturk, Y., Lee, D.W., Mandaci, S., Osyczka, A., Prince, R.C., and Daldal, F. (2008). Soluble variants of *Rhodobacter capsulatus* membrane-anchored cytochrome *c*_y_ are efficient photosynthetic electron carriers. J Biol Chem 283, 13964–13972.

Pawlik, G., Kulajta, C., Sachelaru, I., Schroder, S., Waidner, B., Hellwig, P., Daldal, F., and Koch, H.G. (2010). The putative assembly factor CcoH is stably associated with the *cbb*_3_-type cytochrome oxidase. J Bacteriol 192, 6378–6389.

Quintana-Cabrera, R., and Soriano, M.E. (2019). ER Stress Priming of Mitochondrial Respiratory suPERKomplex Assembly. Trends Endocrinol Metab 30, 685–687.

Sakamoto, J., Matsumoto, A., Oobuchi, K., and Sone, N. (1996). Cytochrome *bd*-type quinol oxidase in a mutant of *Bacillus stearothermophilus* deficient in *caa*_3_-type cytochrome *c* oxidase. FEMS Microbiol Lett 143, 151–158.

Schneidman-Duhovny, D., Inbar, Y., Nussinov, R., and Wolfson, H.J. (2005). PatchDock and SymmDock: servers for rigid and symmetric docking. Nucleic Acids Res 33, W363–367.

Slavin, M., and Kalisman, N. (2018). Structural Analysis of Protein Complexes by Cross-Linking and Mass Spectrometry. Methods Mol Biol 1764, 173–183.

Smith, M.A., Finel, M., Korolik, V., and Mendz, G.L. (2000). Characteristics of the aerobic respiratory chains of the microaerophiles *Campylobacter jejuni* and *Helicobacter pylori*. Arch Microbiol 174, 1–10.

Solmaz, S.R., and Hunte, C. (2008). Structure of complex III with bound cytochrome *c* in reduced state and definition of a minimal core interface for electron transfer. J Biol Chem 283, 17542–17549.

Sone, N., Sekimachi, M., and Kutoh, E. (1987). Identification and properties of a quinol oxidase super-complex composed of a *bc*_1_ complex and cytochrome oxidase in the thermophilic bacterium PS3. J Biol Chem 262, 15386–15391.

Sousa, J.S., Mills, D.J., Vonck, J., and Kuhlbrandt, W. (2016). Functional asymmetry and electron flow in the bovine respirasome. Elife 5.

Sousa, J.S., and Vonck, J. (2019). Respiratory supercomplexes III_2_IV_2_ come into focus. Nat Struct Mol Biol 26, 87–89.

Stroh, A., Anderka, O., Pfeiffer, K., Yagi, T., Finel, M., Ludwig, B., and Schagger, H. (2004). Assembly of respiratory complexes I, III, and IV into NADH oxidase supercomplex stabilizes complex I in *Paracoccus denitrificans*. J Biol Chem 279, 5000–5007.

Swierczek, M., Cieluch, E., Sarewicz, M., Borek, A., Moser, C.C., Dutton, P.L., and Osyczka, A. (2010). An electronic bus bar lies in the core of cytochrome *bc*_1_. Science 329, 451–454.

Turba, A., Jetzek, M., and Ludwig, B. (1995). Purification of *Paracoccus denitrificans* cytochrome *c*_552_ and sequence analysis of the gene. Eur J Biochem 231, 259–265.

Valkova-Valchanova, M.B., Saribas, A.S., Gibney, B.R., Dutton, P.L., and Daldal, F. (1998). Isolation and characterization of a two-subunit cytochrome *b*-*c*_1_ subcomplex from *Rhodobacter capsulatus* and reconstitution of its ubihydroquinone oxidation (Q_o_) site with purified Fe-S protein subunit. Biochemistry 37, 16242–16251.

Verissimo, A.F., Sousa, F.L., Baptista, A.M., Teixeira, M., and Pereira, M.M. (2007). Thermodynamic redox behavior of the heme centers of *cbb*_3_ heme-copper oxygen reductase from *Bradyrhizobium japonicum*. Biochemistry 46, 13245–13253.

Weyer, K.A., Lottspeich, F., Gruenberg, H., Lang, F., Oesterhelt, D., and Michel, H. (1987). Amino acid sequence of the cytochrome subunit of the photosynthetic reaction centre from the purple bacterium *Rhodopseudomonas viridis*. EMBO J 6, 2197–2202.

Wikstrom, M., Krab, K., and Sharma, V. (2018). Oxygen Activation and Energy Conservation by Cytochrome *c* Oxidase. Chem Rev 118, 2469–2490.

Winstedt, L., and von Wachenfeldt, C. (2000). Terminal oxidases of *Bacillus subtilis* strain 168: one quinol oxidase, cytochrome *aa*_3_ or cytochrome *bd*, is required for aerobic growth. J Bacteriol 182, 6557–6564.

Wiseman, B., Nitharwal, R.G., Fedotovskaya, O., Schafer, J., Guo, H., Kuang, Q., Benlekbir, S., Sjostrand, D., Adelroth, P., Rubinstein, J.L., et al. (2018). Structure of a functional obligate complex III_2_IV_2_ respiratory supercomplex from *Mycobacterium smegmatis*. Nat Struct Mol Biol 25, 1128–1136.

Wu, M., Gu, J., Guo, R., Huang, Y., and Yang, M. (2016). Structure of Mammalian Respiratory Supercomplex I_1_III_2_IV_1_. Cell 167, 1598–1609 e1510.

Xia, D., Esser, L., Elberry, M., Zhou, F., Yu, L., and Yu, C.A. (2008). The road to the crystal structure of the cytochrome bc_1_ complex from the anoxigenic, photosynthetic bacterium *Rhodobacter sphaeroides*. J Bioenerg Biomembr 40, 485–492.

Yang, J., Yan, R., Roy, A., Xu, D., Poisson, J., and Zhang, Y. (2015). The I-TASSER Suite: protein structure and function prediction. Nat Methods 12, 7–8.

Yang, J., and Zhang, Y. (2015). Protein Structure and Function Prediction Using I-TASSER. Curr Protoc Bioinformatics 52, 5 8 1–5 8 15.

Yu, C.A., Wen, X., Xiao, K., Xia, D., and Yu, L. (2002). Inter- and intra-molecular electron transfer in the cytochrome *bc*_1_ complex. Biochim Biophys Acta 1555, 65–70.

Zeng, H., Wang, S., Zhou, T., Zhao, F., Li, X., Wu, Q., and Xu, J. (2018). ComplexContact: a web server for inter-protein contact prediction using deep learning. Nucleic Acids Res 46, W432–W437.

