## Supplemental Information for "Cryo-EM Structures of Respiratory *bc*_1_-*cbb*_3_ type CIII_2_CIV Supercomplex and Electronic Communication Between the Complexes"

[1]Department of Biology, University of Pennsylvania, Philadelphia, PA, 19104; [2]Biochemistry and Molecular Biophysics Graduate Group, Perelman School of Medicine, University of Pennsylvania, Philadelphia, PA 19104; [#]Institute of Biochemistry and Molecular Biology, Faculty of Medicine, Albert Ludwigs University of Freiburg, 79104 Freiburg, Germany; [3]School of Computer Science and Engineering, Institute of Life Sciences, The Hebrew University of Jerusalem, Jerusalem, 9190401, Israel and [4]Department of Biochemistry and Biophysics, Perelman School of Medicine, University of Pennsylvania, Philadelphia, PA 19104

**Running title**: Bacterial respiratory cytochrome *$bc_1$-$cbb_3$* supercomplex

**Key words**: cytochrome *$bc_1$* or Complex III; Cytochrome *$cbb_3$* oxidase or Complex IV; respiratory super-complex; electron carrier cytochrome *c*; membrane-anchored cytochrome $c_y$; soluble cytochrome $c_2$; *Rhodobacter capsulatus*; respiratory electron transport chain

Kenji Murakami:

### Methods

#### *Bacterial strains and growth conditions*

Bacterial strains and plasmids used are listed in **Table S1**. LB medium supplemented as appropriate with ampicillin, gentamicin, kanamycin or tetracycline at 100, 12, 50 or 12.5 μg/mL, respectively, was used for growing *E. coli* strains at 37 °C (Darrouzet and Daldal, 2002). *R. capsulatus* strains were grown chemoheterotrophically under semi-aerobic/dark conditions at 35 °C on enriched (MPYE) medium, supplemented as needed with gentamicin, kanamycin, spectinomycin, or tetracycline at 3, 10, 10 or 2.5 μg/mL, respectively (Davidson et al., 1992). As needed, colonies were stained for $cbb_3$-type CIV activity by incubating plates with a 1:1 (v/v) mixture of 35 mM 1-naphtol and 30 mM N,N-dimethyl-1,4-phenylenediamine (NADI-staining) as done recently (Khalfaoui-Hassani et al., 2018)

#### *Molecular genetic techniques*

**Construction of *petABC::ccoP*-His$_8$ (cyt *bc*$_1$-CcoP) fusion.** Using the primers Fw-ccoP (StuI) and Rv-ccoP (HindIII) (**Table S2**), *petABC::ccoP*-His$_8$ fusion was constructed by ligating in-frame the PCR amplified 1.13 kb StuI-HindIII fragment containing *ccoP* (with a C-terminal His$_8$ tag) to the 3' end of *petABC*, after elimination of the stop codon of *petC* and start codon of *ccoP* on StuI-HindIII digested plasmid pMTSI, to yield pYO60 (**Table S1**). The 3.85 kb BamHI fragment of pYO60 carrying *petABC::ccoP* fusion was transferred to the plasmids pBSII and pRK415 using the same sites, yielding pYO63 and pYO76, respectively. The linker-spaced versions of *petABC::ccoP* fusion (*petABC::L2-ccoP*, *petABC::L3-ccoP* and *petABC::L4-ccoP*) were also constructed by exchanging the StuI-HindIII fragment of pYO63 carrying *petABC-ccoP* with the PCR amplified linker added versions, yielding pYO77, pYO78 and pYO80, respectively. L2, L3 and L4 linkers were introduced via the primers F-L2, F-L3 and F-L4 (**Table S2**) and contained the amino acid sequences of $NH_2$-GGSGGGSG-COOH, $NH_2$-GGSGGGSGGGSG-COOH and $NH_2$-ASIAGGRTASGP-COOH, respectively. The 3.9 kb KpnI-XbaI fragments carrying these linkers added versions were cloned to pRK415, yielding pYO81, pYO82 and pYO83, respectively. All constructs were subsequently verified by DNA sequencing. As the protein yields and enzymatic activities of all fusion super-complexes (SCs) were similar, only pYO76, which has no linker (*i.e.*, native-like fusion), was used for subsequent work.

**Construction of *petABC::ccoP::cycY-Flag* (cyt *bc*$_1$-CcoP-cyt *c*$_y$-Flag) fusion.** Using the primers Fw-cp-cy (BalI) and Rv-cp-cy (BglII+HinIII), the C-terminally Flag tagged *cycY* lacking its N-terminal TMH anchor (*i.e.*, amino acid residues 1-30) was PCR amplified (**Table S2**). The 619 bp long PCR fragment containing *cycY-Flag* without its anchor was digested with BalI-HinIII and exchanged with its counterpart that encompasses the 3' end of *ccoP* on pYO63 to yield the *petABC::ccoP::cycY-Flag* fusion carried by pYO91. The 4.5 kb KpnI-XbaI fragment of pYO91 containing this fusion was cloned into pRK415 using the same sites to yield pYO92 (**Table S1**).

**Chromosomal inactivation of CIII$_2$ (cyt *bc*$_1$) and CIV (*cbb*$_3$-type Cox) structural genes, and construction of strains producing bipartite and tripartite fusion SCs.** The *Δ*(*petABC::gm*) deletion-insertion allele carried by pYO34 (Ozturk et al., 2008) was transferred using the gene transfer agent (GTA) (Yen et al., 1979) into the chromosome of *R. capsulatus* strain MG1 (*ccoP::kan*) to yield YO12, providing a mutant background lacking both CIII$_2$ and the CcoP subunit of CIV. In later experiments, a similar strain, M7G-CBC1 that also lacks both CIII$_2$ and the CcoP subunit but overproduces the CcoN and CcoO subunits of CIV (Gray et al., 1994), was opted as a better background. The plasmids pYO76 and pYO92 encoding the bipartite (*bc*$_1$-*cbb*$_3$) type and tripartite (*bc*$_1$-*ccbb*$_3$) type SCs, respectively, were conjugated into YO12 and M7G-CBC1 strains using triparental mating (Daldal et al., 1986), to yield pYO76/YO12 and pYO92/M7G-CBC1 used for protein purification.

#### *Purification and characterization of fusion SCs*

**Protein purification**. *R. capsulatus* cells (~25-30 g from 8 L media) were resuspended in a final volume of 35-40 mL of buffer A (50 mM Tris-HCl, pH 8.0, 100 mM NaCl, 20 mM EDTA, 1 mM aminocaproic acid and 1 mM 4-(2-Aminoethyl) benzenesulfonyl fluoride hydrochloride (AEBSF) supplemented with Pierce Protease Inhibitor (Thermo Scientific, 1 mini tablet per 30 mL). Cells were disrupted by three passages through a French Pressure Cell (SLM Aminco) at 13,000 psi in the presence of 5-10 mg of DNase I (GoldBio), and cell debris removed by centrifugation at 27,000 x g for 30 min (Darrouzet and Daldal, 2002). Chromatophore membranes were sedimented by centrifugation at 190,000 x g for 2 h and resuspended in buffer A. The protein concentration was determined using the Pierce Protein BCA assay kit (Thermo Scientific), the suspension was supplemented with 10% glycerol and adjusted to a protein concentration of 10 mg/mL with buffer A. Membrane proteins were solubilized with 2% n-dodecyl-ß-D-maltoside (DDM, Anatrace) for 15 min on ice, and non-solubilized materials sedimented by centrifugation at 100,000 g for 15 min. The supernatant was subjected to chromatography, as appropriate.

The His-tagged bipartite SC was purified by anion exchange chromatography on Bio-Gel A-50 (Bio-Rad), followed by affinity chromatography on Ni-Sepharose High Performance (GE Healthcare). Solubilized proteins were loaded onto a 40 mL Bio-Gel column equilibrated with Tris buffer (50 mM Tris-HCl, pH 8.0, 100 mM NaCl, 1% glycerol and 0.01% DDM) and washed with three column volumes (CV) of the buffer. Another washing step with 3 CV of Tris buffer, but containing 150 mM NaCl, was done to eliminate weakly bound proteins, and elution was carried out with a linear, five CV gradient from 150-400 mM NaCl in Tris buffer. Fractions containing both cyt *c* reductase and cyt *c* oxidase activities were combined and loaded onto a 7 mL Ni-Sepharose column equilibrated with the binding buffer (50 mM Tris-HCl, pH 8.0, 500 mM NaCl, 20 mM imidazole, 1% glycerol and 0.01% DDM). After washing with 30 mL of binding buffer, the column was eluted with a linear 100 mL gradient from 20-260 mM imidazole.

The Flag-tagged tripartite SC was purified by affinity chromatography using the Anti-Flag affinity gel (Bimake). The supernatant containing solubilized proteins was loaded onto a 1 mL column equilibrated with TBS buffer (50 mM Tris-HCl, pH 7.4, 150 mM NaCl, 0.01% DDM and 1 mM AEBSF), washed with 10 ml TBS and eluted with 5 mL of Flag (DYKDDDDK) peptide (100 μg/mL) (Sigma) in TBS buffer.

Eluted proteins were concentrated (Amicon Ultra-15, 30 kDa MWCO) (Millipore) to a final volume of 150-200 μL and loaded onto a Superose 6 Increase 10/300 GL (GE Healthcare) size exclusion column equilibrated with TBS buffer, and eluted with the same buffer at a flow rate of 0.4 mL/min. Peak fractions were combined and concentrated, and PD MiniTrap G-25 columns (GE Healthcare) were used for buffer exchanges, as needed.

**Protein analyses**. Native PAGE was performed according to (Wittig et al., 2006) using 4-13% gels, and SDS-PAGE according to (Laemmli, 1970) using 12.5% or 18% gels. Gels were stained with Coomassie Brilliant Blue (Bio-Rad) or colloidal silver according to (Bartsch et al., 2012). Immunoblot analysis was performed as earlier (Valkova-Valchanova et al., 1998) and membranes were stained with polyclonal antibodies specific of *R. capsulatus* cyt *b* (Davidson et al., 1992) and monoclonal anti-rabbit-IgG-alkaline phosphatase (Sigma) used at 1:10.000 dilution. CIV in-gel activity was revealed by incubating native gels with 0.5 mg/ml 3,3'-diaminobenzidine in 50 mM Na-phosphate, pH 7.2 (Yan and Forster, 2009). The *c*-type cyts were revealed by their heme peroxidase activity, using 3,3',5,5'-tetramethylbenzidine (TMBZ) and hydrogen peroxide ($H_2O_2$) according to (Thomas et al., 1976). Briefly, gels were washed with 0.25 M Na-acetate buffer (pH 5) and incubated with 6.3 mM 3,3',5,5'-tetramethylbenzidine (TMBZ) in 30% methanol, 0.25 M Na-acetate, pH 5.0. After incubation for 1 h, 0.4% hydrogen peroxide were added to reveal the peroxidase activity of *c*-type cyts.

Reduced *minus* oxidized spectra were recorded in 50 mM MOPS, pH 7.0 using a Cary 60 UV-Vis spectrophotometer (Agilent). Usually, 50 µg of purified protein was first fully oxidized by adding a grain of $K3[Fe(CN)_6]$ and then gradually reduced with a few grains of sodium ascorbate or sodium dithionite, as appropriate. Protein concentrations were determined with a NanoDrop 2000c (Thermo Scientific) using the $A_{280}$ method (Gill and von Hippel, 1989).

**Enzyme activities.** Cyt *c* reductase (cyt $bc_1$) activity was determined as described (Atta-Asafo-Adjei and Daldal, 1991). Briefly, 10 mM stock solution of 2,3-dimethoxy-5-methyl-6-decyl-1,4-benzoquinone (DB) (Sigma) in DMSO was reduced to $DBH_2$ with a few grains of sodium borohydride and excess borohydride was quenched by adding HCl to a final pH of 6.0. For assays, 40 µM $DBH_2$ were added to 25 µM horse heart cyt *c* (Sigma) in 500 µL assay buffer (40 mM sodium phosphate, pH 7.4, 20 mM sodium malonate, 0.5 mM EDTA, 0.5 mM KCN and 0.01% DDM) in a stirred cuvette, and the non-enzymatic rate of cyt *c* reduction was recorded at 550 nm for 1 min (Cary 60 UV-Vis spectrophotometer, Agilent). The reaction was initiated by adding 1-2 µg of purified protein and cyt *c* reduction was monitored. The enzymatic activity was inhibited by adding 1 µM stigmatellin (Fluka), a specific inhibitor of cyt $bc_1$-type $CIII_2$.

The CIV (cyt $cbb_3$-Cox) activity was determined as described earlier (Peters et al., 2008). Reduced horse heart cyt *c* (Sigma) was prepared by incubating a 1-2 mM stock solution with 10 mM sodium dithionite for 15 min at room temperature, excess dithionite was removed by gel filtration using a PD-10 desalting column (GE Healthcare), and the final concentration of reduced cyt *c* was calculated based on its absorption at 550 nm and an ε=20 $mM^{-1}cm^{-1}$. 20 μM reduced cyt *c* in 500 μL assay buffer (10 mM Tris-HCl, pH 7.0, 100 mM KCl) were prepared in a stirred cuvette (Cary 60 UV-Vis spectrophotometer), and the baseline was recorded for 1 min. The reaction was started by adding 1-2 μg of purified protein, and cyt *c* oxidation monitored at 550 nm. As needed, the enzymatic activity was inhibited by addition of 250 μM KCN, as a specific inhibitor of $cbb_3$-Cox.

$DBH_2$ dependent oxygen consumption was monitored using a mini Clark-type oxygen electrode (Instech Laboratories, PA) at a constant temperature of 25 °C. The assay was performed in the same buffer as that used for cyt *c* reductase activity (40 mM sodium phosphate, pH 7.4, 20 mM sodium malonate, 0.5 mM EDTA and 0.01% DDM), but without the $cbb_3$-Cox inhibitor KCN. A baseline was recorded after adding 100 μM $DBH_2$ to 1 mL of the assay buffer, and the reaction started by adding 30 μg of purified protein. As needed, purified protein was pre-incubated with a 1:1 (w/w) mixture of *E. coli* polar lipids (Avanti), which improved the activity. 20 μM horse heart cyt *c* (Sigma), 250 μM KCN or 1 μM of stigmatellin were used as specificity controls.

**Identification of protein subunits by mass spectrometry.** The SDS/PAGE bands were excised, and after reduction (dithiothreitol; Sigma) and alkylation (iodoacetamide; Bio-Rad) subjected to in-gel trypsin digestion (Promega, Sequencing Grade Modified Trypsin) overnight at 37 °C. Peptides eluted from the gel samples were dried, desalted using ZipTips (Millipore U-C18 P10, Millipore), lyophilized and stored at −80 °C. They were resuspended in 10 μL 5% acetonitrile/0.1% formic acid prior to mass spectrometry, using either a LCQ Deca XP+ ion trap, or a Q-Exactive Quadrupole-Orbitrap, mass spectrometer (both from Thermo Fisher Scientific) (Selamoglu et al., 2020). The LCQ Deca XP+ mass spectrometer was coupled to a Thermo-Dionex

LC Packings Ultimate Nano HPLC system controlled by Thermo Xcalibur version 2.0 software. Peptides were separated on a 15 cm C18 nanocolumn (Thermo-Dionex, NAN-75-15-03-PM) using a 45-min linear gradient from 4% to 40% buffer B (100% acetonitrile, 0.1% formic acid), followed by a 7-min gradient from 40% to 80% buffer B and 8-min wash with 80% B (constant flow rate 150 nL·min$^{-1}$). MS/MS data were acquired in data-dependent analysis mode with dynamic exclusion enabled (repeat count: 3, exclusion duration: 3 min). Full MS survey scans (mass range 300–2000 *m*/*z*) were followed by MS/MS fragmentation (normalized collision energy 35) of the top 3 most intense ions. The Q-Exactive Quadrupole-Orbitrap mass spectrometer was coupled to an Easy-nLC™ 1000 nano liquid chromatography system (Thermo Fisher Scientific), and samples were loaded in buffer A (0.1% formic acid) onto a 20-cm-long fused silica capillary column (75 μm ID), packed with reversed-phase Repro-Sil Pur C18-AQ 3 μm resin (Dr. Maisch GmbH, Ammerbuch, Germany). Peptides were eluted using a 45-min linear gradient from 4% to 40% buffer B (100% acetonitrile, 0.1% formic acid), followed by a 7-min gradient from 40% to 80% buffer B and 8-min wash with 80% B (constant flow rate 300 nL·min$^{-1}$). The Q-Exactive was operated in data-dependent acquisition mode with dynamic exclusion enabled (repeat count: 1, exclusion duration: 20 s). Full MS survey scans (mass range 300–1600 *m*/*z*) at high resolution (70 000 at 200 *m*/*z*) were followed by MS/MS fragmentation of the top 15 most intense ions with higher energy collisional dissociation at a normalized collision energy of 22 (resolution 17 500 at 200 *m*/*z*). Dual lock mass calibration was enabled with 371.101233 and 445.120024 *m*/*z* background ions.

MS spectra were searched against the *R. capsulatus* protein database (https://www.uniprot.org, last modified 01/15/2020; *Rhodobacter capsulatus* (strain ATCC BAA-309 / NBRC 16581 / SB1003)) using Proteome Discoverer 1.4 (Thermo Fisher Scientific) with Sequest-HT search engine. Search parameters were set to full trypsin digestion, with maxima of three missed cleavages and three modifications per peptide. Oxidation of methionine (+16 Da) and carbamidomethylation of cysteine (+57 Da) were selected as dynamic modifications. Precursor and fragment ion tolerances were set to 2 Da and 1 Da, respectively, for the LCQ DecaXP+, and 10 ppm and 0.6 Da, respectively, for the Q-Exactive data. False discovery rates by target-decoy search (FDR) were set to 0.01 (high confidence) and $X_{corr}$ filter based on charge (z) were: > 2 for $z = 2$; > 2.5 for $z = 3$; and > 2.6 for $z = 4$ (Selamoglu et al., 2020). All identifications are listed in **Table S3**.

***Binding of cyt $c_y$, soluble variant of cyt $c_y$ (cyt S-$c_y$) and cyt $c_2$ to purified proteins***

**Cyt $c_y$ and cyt S-$c_y$.** Binding of purified cyt $c_y$ (Myllykallio et al., 1997), or a variant of it containing only the soluble cyt *c* domain (cyt S-$c_y$) (residues 99-199) to the bipartite SC was assayed by mixing a 5-10 fold molar excess of purified cyt *c* with 125 µg of purified SC in a final volume of 150 µL TBS buffer (50 mM Tris-HCl, pH 7.4, 150 mM NaCl and 0.01% DDM). The mixture was separated by chromatography using Superose 6 Increase size exclusion column, and elution fractions were analyzed by SDS-PAGE followed by silver staining. Cyt S-$c_y$ was purified from the *R. capsulatus* strain pYO135/FJ2-R4 (**Table S1**) grown under photosynthetic conditions to maximize its yield (Ozturk et al., 2008). Cells were washed with 20 mM Tris-HCl, pH 8.0 and resuspended 1:5 (w/v) in the same buffer supplemented with 50 mM NaCl. Polymyxin B sulfate (1 mg/mL) was added, and the cell suspension incubated for 75 min on ice with gentle stirring. After centrifugation for 20 min at 10,000 g followed by 3 h at 150,000 g, the supernatant was collected and concentrated to a final volume of 2 ml (Amicon Ultra-15 3 kDa MWCO) (Millipore). Aliquots of 300 µL were loaded onto Superose 6 Increase (GE Healthcare) equilibrated in 30 mM

Tris-HCl, pH 8.0 and eluted in the same buffer. Fractions eluting after 20 mL were combined and concentrated using Amicon Ultra-15 3 kDa MWCO filters (Millipore).

**Cyt $c_2$.** Binding of cyt $c_2$ to $cbb_3$-type CIV was determined by mixing 2-fold molar excess of purified *R. capsulatus* cyt $c_2$ (Holden et al., 1987) with 300 µg of purified CIV (Gray et al., 1994) under low-salt conditions (20 mM Tris-HCl, pH 7.4, 1 mM NaCl and 0.01% DDM). The mixture (250 µL total volume) was loaded onto a Superose 6 Increase (GE Healthcare) sizing column equilibrated with the same buffer to separate the proteins. Elution fractions were concentrated (Amicon Ultra-15, 3 kDa MWCO) (Millipore) and a volume containing 1 to 5 µg of protein was analyzed by SDS-PAGE followed by silver staining.

***Protein cross-linking and mass spectrometry***

**Protein cross-linking using chemical crosslinkers.** 180 µg of purified bipartite SC at a concentration of 1 mg/mL in PBS buffer (50 mM Na-Phosphate, pH 7.4, 150 mM NaCl and 0.01% DDM) mixed with 5-fold molar excess (90 µg) of purified cyt $c_y$ (Myllykallio et al., 1997) were supplemented with 6 mM disuccinimidyl dibutyric urea (DSBU) (Thermo Fisher Scientific), and incubated on ice for 2 h. The reaction was quenched by adding 50 mM of ammonium bicarbonate, and the mixture analyzed by mass spectrometry. 300 µg of purified $cbb_3$-type CIV (Gray et al., 1994) were mixed with a 2-fold molar excess of purified cyt $c_2$ (Holden et al., 1987) at a final protein concentration of 200 µg/mL under low-salt conditions (20 mM Na-phosphate, pH 7.4, 1 mM NaCl and 0.01% DDM), and incubated with 20 mM 4-(4,6-Dimethoxy-1,3,5-triazin-2-yl)-4-methylmorpholinium chloride (DMTMM) (Sigma) for 1 h at room temperature. Similarly, 300 µg of purified $bc_1$-type $CIII_2$ were mixed with 2-fold molar excess of purified cyt $c_2$, and treated with DMTMM as above. In both cases, the reaction was stopped by removing excess DMTMM with a

PD MiniTrap G-25 desalting column (GE Healthcare), and crosslinked proteins were precipitated with 20% (w/v) trichloroacetic acid (TCA, Sigma) at 4°C for 1 h. Proteins were pelleted by centrifugation at 21,000 x g for 15 min and washed with 10% TCA in 0.1 M Tris-HCl and then with acetone (Fisher). The solvent was discarded, the pellet air-dried and then stored at -80°C for analysis by mass spectrometry.

**Mass spectrometry of crosslinked proteins**. Crosslinked proteins were resuspended in an appropriate volume of solution A (2.5% TCA, 50 mM SDS and 50 mM triethylammonium bicarbonate (TEAB) final concentrations) and reduced with 10 mM DTT (US Biological) for 30 min at 30 °C, followed by alkylation with 50 mM iodoacetamide (Sigma Aldrich) for 30 min at 30 °C. The proteins were processed using an S-Trap™ according to the protocol recommended by the supplier (Protifi, C02-mini), and digested with trypsin (Thermo Fisher Scientific) in 1:10 (w/w) enzyme/protein ratio for 1 h at 30 °C. Peptides eluted from this column were vacuum-dried and resuspended with the peptide fractionation-elution buffer for LC-MS [(70% (v/v) LC-MS grade water (Thermo Fisher Scientific), 30% (v/v) acetonitrile (ACN, Thermo Fisher Scientific) and 0.1 % (v/v) trifluoroacetic acid (TFA, Thermo Fisher Scientific)]. Peptides were first fractionated using AKTA Pure 25 with Superdex 30 Increase 3.2/300 (GE Life Science) at a flow rate of 30 μL $min^{-1}$ of the elution buffer, and 100 μL fractions were collected. Based on the elution profile, fractions containing enriched crosslinked peptides of higher molecular masses, were vacuum-dried and resuspended with LC-MS grade water containing 0.1% (v/v) TFA for mass spectrometry analysis. One half of each fraction was analyzed by a Q-Exactive HF mass spectrometer (Thermo Fisher Scientific) coupled to a Dionex Ultimate 3000 UHPLC system (Thermo Fischer Scientific) equipped with an in-house made 15 cm long fused silica capillary column (75 μm ID), packed with reversed-phase Repro-Sil Pur C18-AQ 2.4 μm resin (Dr. Maisch GmbH, Ammerbuch, Germany).

Elution was performed using a gradient from 5% to 45% B (90 min), followed by 90% B (5 min), and re-equilibration from 90% to 5% B (5 min) with a flow rate of 400 nL/min (mobile phase A: water with 0.1% formic acid; mobile phase B: 80% acetonitrile with 0.1% formic acid). Data were acquired in data-dependent MS/MS mode. Full scan MS settings were: mass range 300−1800 m/z, resolution 120,000; MS1 AGC target 1E6; MS1 Maximum IT 200. MS/MS settings were: resolution 30,000; AGC target 2E5; MS2 Maximum IT 300 ms; fragmentation was enforced by higher-energy collisional dissociation with stepped collision energy of 25, 27, 30; loop count top 12; isolation window 1.5; fixed first mass 130; MS2 Minimum AGC target 800; charge exclusion: unassigned, 1, 2, 3, 8 and >8; peptide match off; exclude isotope on; dynamic exclusion 45 sec (Slavin and Kalisman, 2018). Raw files were converted to mgf format with TurboRawToMGF 2.0.8 (Sheng et al., 2015).

**Crosslinked peptide searches.** Search engines MeroX 2.0.0.5 (Gotze et al., 2015), FindXL (Kalisman et al., 2012) and MassAI 19.07 (http://www.massai.dk) were used to identify and validate crosslinked peptides. MeroX was run in RISEUP mode, with default crosslinker mass and fragmentation parameters for DSBU, and in Quadratic mode with default crosslinker mass parameters for DMTMM; precursor mass range, 300–10,000 Da; minimum precursor charge 4; precursor and fragment ion precisions 5.0 and 10.0 ppm, respectively; maximum number of missed cleavages 3; carbamidomethylation of cysteine and oxidation of methionine, as fixed and variable modifications, respectively; results were filtered for score (>10) and false discovery rate, FDR (<1%). FindXL was used to analyze and validate MeroX results for DMTMM crosslinks. The default FindXL parameters were used as described before (Kalisman et al., 2012) with the possible crosslink amino acids adjustments for K, Y, S, or T on one peptide, and E or D on the other peptide. MassAI was used to validate crosslinks identified by MeroX for DSBU with standard settings, except: 5 ppm MS accuracy, 0.05 Da MS/MS accuracy, 3 allowed missed cleavages, and carbamidomethylation of cysteine and oxidation of methionine, as fixed and variable modifications, respectively. Only the crosslinks that were consistently identified by two different search engines (MeroX and FindXL for DMTMM, **Table S5**, and MeroX and MassAI for DSBU, **Table S6**) were used for method validation (**Fig. S9**) and subsequent docking experiments. Visualization of the crosslinks in $CIII_2CIV$ structure used Chimera (Pettersen et al., 2004) with the Xlink Analyzer plug-in (Kosinski et al., 2015).

***Negative staining and cryo-EM sample preparation***

For negative staining, purified SCs were diluted to 0.01-0.05 mg/mL concentrations in TBS (50 mM Tris-HCl, pH 7.4, 150 mM NaCl, 0.01% DDM and 1 mM AEBSF) buffer, and 5 µL were applied to glow-discharged (20 sec, 25 mA, Pelco easiGlow) carbon-coated Cu grids (CF300-CU, EMS), incubated for 1 min, then stained with 2% uranyl acetate. Grids were imaged on FEI Tecnai 12 transmission electron microscope (TEM) operating at 120 kV, using a CCD camera (Gatan BM-Ultrascan).

For cryo-EM, the tripartite SC was used as purified, whereas the purified bipartite SC was mixed with a 5-fold molar excess of purified cyt $c_y$ (Myllykallio et al., 1997). 2.5 µL of the mixture, containing 3-5 mg/mL proteins, were applied to CFlat holey carbon grids (1.2/1.3-400 mesh or 2/2-300 mesh) (EMS), which were glow discharged (2 min, 25 mA, Pelco easiGlow) before sample application. Grids were blotted for 9 sec at force 0 (CFlat-1.2/1.3), or for 3 sec at force -5 (CFlat-2/2), and flash-frozen in liquid ethane cooled with liquid nitrogen using FEI Vitrobot Mark IV (25°C, 100% humidity). Plunge freezing conditions were optimized using FEI Tecnai TF20 TEM operating at 200 kV, equipped with a FEI Falcon II camera.

***Cryo-EM data acquisition and processing***

All cryo-EM grids were imaged using FEI Titan Krios electron microscope operating at 300 kV. Images of the tripartite SC (SC~$c_y$) were recorded using a Gatan K2 Summit direct electron detector, equipped with an energy quantum filter (20 eV), and operated in super-resolution mode at a nominal magnification of 105,000x, resulting in a binned pixel size of 1.32 Å. Images were dose-fractionated to 40 frames with a total exposure time of 10 sec, and a total dose of 40 $e^-/A^2$. Automated data acquisition was carried out using Latitude software (Gatan), and nominal defocus values varied from 1 to 2.5 μm. Movies were motion corrected using MotionCor2 (Zheng et al., 2017) and CTF parameters were determined with CTFFIND 4.1 (Rohou and Grigorieff, 2015). About 10,000 particles were manually picked and subjected to an initial reference-free 2D classification using Relion 3.0 (Zivanov et al., 2018). Representative classes were selected and used as a template for auto-picking. After sorting and two rounds of 2D classification, ~30,000 particles were retained from the dataset. For 3D classification, an initial model was created using the structure of *R. capsulatus* $CIII_2$ (PDB: 1ZRT) and low-pass filtered to 60 Å using EMAN2 (Tang et al., 2007). A 3D map with a nominal resolution of ~11 Å containing a total of ~12,000 particles (40%) was obtained, showing a $CIII_2$ associated on one side with a single copy of CIV. For further analyses, additional datasets were collected using samples from the same batch and identical imaging conditions. From a total of ~17,000 images, ~1,000,000 particles were automatically picked using the template obtained from the first dataset, and after sorting and two rounds of 2D classification, ~500,000 particles were retained. Various subsets of all images were processed separately, and yielded the same overall results with slightly different versions of individual maps. For clarity, only the paths to the best representative of each of the final maps are shown (**Fig. S3A**, **B** and **C**). For 3D classification, the map obtained from the first dataset (**Fig. S3**, Box 3) was low-pass filtered to 60 Å and used as the initial model. Of the five classes obtained, one showed clear features of a $CIII_2$ dimer associated with a single monomer of CIV (**Fig. S3**, class005 in **A**, class002 in **B** and **C**). The remaining classes showed the same overall shape, but lacked density or resolution in different parts of the structure. Only two of the five maps contained a second monomer of CIV associated with the other monomer of $CIII_2$ (**Fig. S3C**). These two classes showing this feature were combined, yielding a subset of ~ 220,000 particles. These particles were further processed by another round of 2D classification, and subjected to 3D classification, yielding three classes (**Fig. S3C**). Only one of the maps obtained in this second round of 3D classification clearly showed density for a second monomer of CIV. Aligned particles from this group were subclassified into six classes in another round of 3D classification using a soft mask and no image alignment. Due to the low number of particles (~5,000) per class, the resolution of the maps remained limited (< 10Å), and could not be further improved through 3D refinement.

Refinement of the individual maps (**Fig. S3A** and **B**) led to relatively low resolution reconstructions, particularly in the CIV portion of the map, probably due to greater structural heterogeneity in the sample with respect to this portion of the tripartite SC. The highest resolution was obtained with a map containing a single copy of CIV, by combining all classes showing well defined features corresponding to it and subjecting them to a second round of 3D classification, focused on CIV portion of the map. For this purpose, a soft mask around CIV was created by fitting the available structure of the closely related *P. stutzeri* homolog (PDB: 3MK7) into the map and low-pass filtering it to 10 Å. All information outside of this mask was subtracted from the aligned particles, and the remaining particles were subjected to a masked classification without image alignment, to yield six classes (**Fig. S3A**). Alternatively, a cylindrical mask was wrapped around the CIV portion of the map and used in a similar procedure (**Fig. S3B**). In each case, classes showing the highest level of details were retained, and the entire unsubtracted particles dataset subjected to 3D auto-refinement followed by per-particle CTF refinement, Bayesian polishing and post-processing using Relion 3.0. A major difference between the maps was the orientation of CIV relative to $CIII_2$ (**Fig. S4**). The two extreme conformations were represented by the maps SC-1A (**Fig. S3A**) and SC-1B (**Fig. S3B**), which were refined to 6.1 Å and 7.2 Å resolutions, respectively (**Table 1**).

Images of the bipartite SC supplemented with cyt $c_y$ (SC+$c_y$) were recorded by a Gatan K3 direct electron detector equipped with an energy quantum filter (20 eV) and operated in counting mode at a nominal magnification of 64,000x, corresponding to a pixel size of 1.36 Å. Images were dose-fractionated to 80 frames with a total exposure time of 3.1 sec and a total dose of 40 $e^-/A^2$. Nominal defocus values varied from 1 to 2.5 μm. Automated data acquisition was carried out using Latitude software (Gatan) and image shift (~ 2 μm, 4 images per stage position) was used for accelerated data collection. For this sample 5,480 images were collected in the first session, motion corrected using MotionCor2 (Zheng et al., 2017), and CTF parameters were determined with CTFFIND 4.1 (Rohou and Grigorieff, 2015). For auto-picking, the template previously obtained with the tripartite SC was used, with this and all subsequent steps done in Relion 3.0 (Zivanov et al., 2018). After sorting and two rounds of 2D classification, ~ 228,000 particles were retained (**Fig. S6A**). The same initial model as for the tripartite SC was used for 3D classification into five classes. Two classes (001 and 002) resembled more to the overall shape of a $CIII_2$ without CIV, while the other three classes showed the same overall shape of a $CIII_2CIV$ SC, as seen with the tripartite sample. 37,460 particles corresponding to the SC class with the highest level of detail were combined with SC particles from the second dataset (**Fig. S6B**) to yield the best final map of the bipartite SC. The second dataset consisted of 12,200 images which were collected and processed under the same conditions as the first dataset. 1.6 million particles were auto-picked and after two rounds of 2D classification, classes were split into ~340,000 large particles likely representing $CIII_2CIV$ and ~465,000 particles resembling $CIII_2$ (**Fig. S6B** and **D**, respectively). The former particles were subjected to 3D classification into three classes, and the class most similar to the overall shape of $CIII_2CIV$ was identified. It contained ~118.000 particles, but due to its low level of detail, it was subjected to another round of 2D classification. The 34,819 particles thus retained were combined with 37,460 SC particles from the first dataset (**Fig. S6A**). After 3D classification into five classes, the class with the highest level of detail contained 56% of particles, and clearly showed the overall shape of $CIII_2CIV$. 41,017 aligned particles corresponding to this class were extracted and subjected to a second round of 3D classification without image alignment using a soft mask. Of the six 3D classes, the one with the highest nominal resolution contained 14,978 particles (36%) and was subjected to 3D auto-refinement and post processing, followed by per-particle CTF refinement and Bayesian polishing. After a second round of 3D auto-refinement and post processing, the final map (SC-2A) of the bipartite $CIII_2CIV$ was obtained at a nominal resolution of 5.18 Å (**Table 1**). This map was very similar to that (SC-1A) of the tripartite $CIII_2CIV$, except for a slightly higher resolution, and strongly improved density and resolution of the extra TMHs at the interface. Unlike the tripartite samples, no major class corresponding to the conformation seen in SC-1B was identified.

As the bipartite SC samples contained a significant amount of $CIII_2$ particles without CIV, subsets of smaller 2D classes consistent with a $CIII_2$ dimer were selected and processed separately. Of the ~376,000 total particles retained from the dataset 1, ~267,000 were identified as resembling

$CIII_2$. After a second round of 2D classification (**Fig. S6C**), ~213,000 particles were retained and used to generate an *ab initio* 3D model. This model was lowpass filtered to 60 Å and used as initial model for the 3D classification of ~465,000 $CIII_2$ particles from dataset 2 (**Fig. S6D**). Particles corresponding to the class with the highest resolution and level of detail (~185,000) were combined with the $CIII_2$ particles from dataset 1 to yield ~400,000 particles, and subjected to 3D classification. The ~170,000 particles corresponding to the best class were further processed following two different strategies. $CIII_2$ being a homodimer based on X-ray structures, C2 symmetry was applied in the next round of 3D classification and in all subsequent steps (**Fig. S6E**). The best class contained ~120,000 particles (70%) which were extracted and, similar to $CIII_2CIV$, subjected to another round of 3D classification without image alignment using a soft mask. Of the six classes obtained, some showed the external domain (ED) of both monomers of the FeS protein close to heme $b_L$ ("**b** position", b-b) while other classes showed both EDs close to heme $c_1$ ("**c** position", c-c). Individual classes were subjected to 3D auto-refinement and post processing, followed by two rounds of per-particle CTF refinement, Bayesian polishing, 3D auto-refinement and post processing. Only the final map with the highest nominal resolution (3.30Å) is shown (**Fig. S6E**, $CIII_2$) (**Table 2**). This map contained 37,997 particles and showed both FeS-EDs close to b position, but at a lower local resolution and occupancy than the rest of the map, indicating structural flexibility and conformational heterogeneity. A similar approach was used for **Fig. S6F**, except that no C2 symmetry was applied in the two rounds of 3D classifications. Of the three best classes obtained in the second round, one showed both EDs in the b position, one showed both in the c position, and one showed a heterodimeric conformation with one monomer in the b and the other in the **c** position. Each of the maps was further processed as in **Fig. S6E**, but C2 symmetry was only applied in case of the homodimeric structures. The final maps were **$CIII_2$ c-c** with both

ED's in the **c** position at 3.8Å, **$CIII_2$ b-b** with both ED's in the b position at 3.5Å, and **$CIII_2$ b-c** with one ED in the b and the second in the **c** position at 4.2Å (**Table 2**). The nominal resolutions thus obtained were slightly lower than in map **$CIII_2$** (**Fig. S6E)**. However, by omitting C2 symmetry application to the 3D classifications, the conformational heterogeneity of the FeS protein EDs was resolved, and a subset of particles showing a heterodimeric conformation was identified. The different locations (*i.e.*, b or c positions) of the ED's were clear in the three maps obtained (**Fig. S6F)** but their occupancy and local resolutions remained low compared to the rest of the structure.

A third dataset was also collected and yielded ~303,000 particles after the first 2D classification. Adding these particles to the first two datasets did not improve the maps shown in **Fig. S6A** and **B**, but it turned out to be informative. As shown in **Fig. S6G**, particles from dataset 3 were subjected to two rounds of 3D classification. Interestingly, an extra density near the periplasmic domain of CcoP, could be seen in some classes. This was not observed in any subclass obtained from the datasets 1 and 2, and tentatively thought to correspond to the cyt *c* domain of cyt $c_y$. Focused classification followed by 3D auto-refinement and post processing, using wider soft masks around the periplasmic domain of CcoP to avoid cutting-off any of the extra density, led to a final map (SC-2B) with limited resolution (10.5Å), and could not be further improved by sub-classification due to the low number of particles in each subclass.

#### *Refinement of R. capsulatus $CIII_2$ in the cryo-EM maps*

The X-ray based structure of *R. capsulatus* $CIII_2$ (PDB: 1ZRT) was fitted into map **$CIII_2$** (EMD-22189) which had the highest nominal resolution of all maps obtained (**Fig. S6E**), and refined using Phenix1.16 (Liebschner et al., 2019). The real space refinement approach included four rounds of global minimization, local grid search, morphing and simulated annealing, with the final round also including ADP (B-factor) refinement. Each round included 5 cycles using default settings, and morphing and annealing were performed in each cycle. Secondary structures were determined by Phenix1.16, using default search settings and restrictions were applied during real space refinement. Due to the low occupancy and limited local resolution, the FeS-ED proteins (residues 50-191) were only subjected to rigid body fitting followed by two rounds of global minimization and local grid search (5 cycles each), but not to morphing and simulated annealing. To ensure the correct cofactor geometry, hemes and [2Fe-2S] clusters including their coordinating residues were copied from the high-resolution X-ray structure of the homologous $CIII_2$ (cyt $bc_1$) from *R. sphaeroides* (PDB: 6NHH). Validation was performed using MolProbity (Williams et al., 2018) (http://molprobity.biochem.duke.edu/), and outliers (Ramachandran, rotamer, bonds, angles) were manually corrected in Coot (Emsley et al., 2010), using real space refinement and regularization. The model that was refined in map **$CIII_2$** (EMD-22189) was subsequently used for rigid body fitting into the maps **$CIII_2$ b-b**, **$CIII_2$ c-c** and **$CIII_2$ b-c** (**Fig. S6F**) (**Table 2**). Each chain was treated as one separate body, except the FeS protein, which was split into its TMH (11 to 49) and ED (50 to 191) residues. After the procedure, the linker between the ED and TMH (residues 40-50) were remodeled in Coot using real space refinement and regularization.

#### ***Structural modeling of R. capsulatus cbb$_3$-type CIV subunits, CcoH and cyt c$_y$***

**Modeling of *R. capsulatus cbb$_3$*-type CIV.** The *Pseudomonas stutzeri cbb$_3$*-type CIV structure (PDB: 5DJQ) (sequence identities for *R. capsulatus* CcoN: 68%, CcoO: 55%, CcoP: 34%) was used as a template, and comparative models were computed using MODELLER v9.18 (Webb and Sali, 2014) with defined secondary structure and crosslinking restraints. The secondary structures for the regions without any template coverage (CcoP residues 104 to 112, 176 to 178, 184 to 197 and 204 to 215; CcoO residues 172 to 185 and 192 to 199), due to the insertion sequences that are present only in *R. capsulatus* CIV, were estimated using PsiPred Protein Sequence Analysis Workbench (Buchan and Jones, 2019; Jones et al., 1999), and added as restraints to MODELLER. Cross-link distances were added as Gaussian restraints to MODELLER with a mean of 18.0Å and a standard deviation of 1.0Å . Problematic loop regions (a total of ten loops longer than four amino acids) were detected by MolProbity (Chen et al., 2010; Williams et al., 2018) and remodeled using MODELLER "slow" refinement method (**Table S7**). As *R. capsulatus* CcoN is longer than that of *P. stutzeri* with a predicted extra N-ter TMH, whereas CcoP is shorter with only one predicted N-ter TMH instead of two, these regions were not included into the model. The regions CcoO 179-214 as well as CcoP 1-12, 53-59, 161-173 and 272-280 without template coverage that were not supported by the cryo-EM maps were omitted.

**Modeling of CcoH and cyt *c* domain of $c_y$.** For CcoH (residues Met 1 to Thr35) five *ab initio* models were obtained using I-TASSER server (Yang et al., 2015). All models were almost identical, and one with the lowest energy score was retained. For *R. capsulatus* cyt $c_y$, its homologs with known structures were detected using HHpred (Soding, 2005), and the structure of cyt *c* domain of cyt $c_{552}$ from *Paracoccus denitirificans* (PDB: 3M97; sequence identity: 61%) was used as a template for the soluble cyt *c* domain of $c_y$ and comparative models were computed using MODELLER (Webb and Sali, 2014).

**Modeling of the extra N-ter TMH of CcoN (TMH0).** A 29-residue model consisting of CcoN residues Arg25 to Asp53, which includes the predicted transmembrane region (Met30-Leu48, UniProtKB, D5ARP4), was obtained from the I-TASSER server (Zhang et al., 2016). The alpha helical region included residues Leu27 to Thr50, and the model including the residues Arg25 to

Leu48 was manually fitted into the density map for visualization purpose only (**Figs. 3** and **4**). Corresponding coordinates were not deposited in the PDB because the registration could not be determined due to the lack of side chain density.

**Modeling of cyt $c_y$ and CcoH TMHs interactions.** The 30 residues long cyt $c_y$ TMH was manually docked into the density map by positioning Phe15 and Tyr21 into the corresponding densities, and refined by real-space refinement and regularization using Coot. The I-TASSER model of CcoH was docked into the map by moving it along the corresponding density, retaining its close association with cyt $c_y$ TMH. The best fit was found when Ala23 and Val24 of CcoH were located at the interface with cyt $c_y$ TMH. The interaction of the two helices was optimized by GalaxyWEB (Ko et al., 2012), and the model with the lowest energy profile was refined in map SC-2A using Phenix 1.17. The refinement strategy included 5 cycles of global minimization, rigid body fitting (where cyt $c_y$ and CcoH chains were treated as separate bodies), local grid search and ADP (B-factor) refinement with default settings for all steps. Secondary structure restrictions were applied to the $\alpha$-helical parts of CcoH as predicted by I-TASSER (residues 12 to 34) and to cyt $c_y$ as predicted by GalaxyWEB (residues 2 to 9 and 12 to 29). $CIII_2$ and CIV models were present during the refinement to keep the rest of the map occupied, but changes made during this procedure were discarded as these models were refined separately.

**Integrative modeling and docking CcoH, cyt $c_y$ and cyt $c_2$ to bipartite SC.** The entire SC was assembled by an integrative modeling approach using the cryo-EM map, XL-MS, co-evolutionary analysis, and subunit models described above. $CIII_2$ and CIV models were fitted into the cryo-EM maps in UCSF Chimera (Pettersen et al., 2004). Additional data about the interaction interfaces between the different subunits were obtained using RaptorX-ComplexContact (Zeng et al., 2018) for each pair of the SC subunits. Based on co-evolution and machine learning, this method predicted the pairs of residues that are in contact. The PatchDock, which is an efficient rigid docking method that maximizes geometric shape complementarity (Schneidman-Duhovny et al., 2005), was used to generate docked configurations of CcoH, cyt $c_2$ and cyt $c_y$ to $CIII_2$ and CIV, as appropriate. The different subunits were docked in parallel and independently from each other.

**Docking of CcoH to CIV**. A total of 16 contacts predicted by RaptorX-ComplexContact, with probabilities higher than 0.5 (**Table S4**), were used as distance restraints for protein-protein docking (Schneidman-Duhovny et al., 2005). The models satisfied 13 of the contacts, and a single cluster of docked models evidenced by the convergence of the top 100 results, was obtained (**Fig. S5**). This cluster coincided with the additional density seen at the SC interface, and CcoH as well as the cyt $c_y$ TMH of were modelled into that density as described above (**Fig. 4C** and **D**).

**Docking of cyt $c_2$ to $CIII_2$ and CIV.** *R. capsulatus* cyt $c_2$ of known structure (PDB: 1C2N) was docked to $CIII_2$ using the three distance restraints between cyt $c_2$ and cyt $c_1$ derived from the protein cross-linking data (**Table S5**). Similarly, cyt $c_2$ was docked to CIV using nine distance restraints provided byXLs (1 crosslink to CcoP and 8 to CcoO, **Table S5**). The models yielded one main cluster in each case, and covered 100% and 89% of the data for $CIII_2$ and CIV, respectively.

**Docking of cyt $c_y$ to $CIII_2$CIV SC**. The model of cyt *c* domain of $c_y$ was docked by PatchDock to bipartite $CIII_2$CIV, which has both FeS-EDs of $CIII_2$ in the b position. Ten distance restraints derived from the XL-MS data (6 with DMTMM, **Table S5**, and 4 with DSBU, **Table S6**) were used, yielding two main clusters (1 and 2) of the docked models on the *p* side of $CIII_2$ (**Fig. S11, Fig. 6B**, **Fig. 7**), and satisfied 100% of the restraints. As the cryo-EM data revealed that $CIII_2$ particles could have their FeS-EDs in the c position, the corresponding $CIII_2$ c-c, $CIII_2$ b-b and $CIII_2$ b-c models were used for docking via Patchdock the cyt *c* domain of $c_y$ onto $CIII_2$.

All the models were ranked using statistically optimized atomic potentials (SOAP) (Dong et al., 2013), and those that have low SOAP scores were retained.

### References

Atta-Asafo-Adjei, E., and Daldal, F. (1991). Size of the amino acid side chain at position 158 of cytochrome *b* is critical for an active cytochrome $bc_1$ complex and for photosynthetic growth of *Rhodobacter capsulatus*. Proc Natl Acad Sci U S A *88*, 492-496.

Bartsch, H., Arndt, C., Koristka, S., Cartellieri, M., and Bachmann, M. (2012). Silver staining techniques of polyacrylamide gels. Methods Mol Biol *869*, 481-486.

Buchan, D.W.A., and Jones, D.T. (2019). The PSIPRED Protein Analysis Workbench: 20 years on. Nucleic Acids Res *47*, W402-W407.

Chen, V.B., Arendall, W.B., 3rd, Headd, J.J., Keedy, D.A., Immormino, R.M., Kapral, G.J., Murray, L.W., Richardson, J.S., and Richardson, D.C. (2010). MolProbity: all-atom structure validation for macromolecular crystallography. Acta Crystallogr D Biol Crystallogr *66*, 12-21.

Daldal, F. (1988). Cytochrome $c_2$-independent respiratory growth of *Rhodobacter capsulatus*. J Bacteriol *170*, 2388-2391.

Daldal, F., Cheng, S., Applebaum, J., Davidson, E., and Prince, R.C. (1986). Cytochrome $c_2$ is not essential for photosynthetic growth of *Rhodopseudomonas capsulata*. Proc Natl Acad Sci U S A *83*, 2012-2016.

Darrouzet, E., and Daldal, F. (2002). Movement of the iron-sulfur subunit beyond the ef loop of cytochrome *b* is required for multiple turnovers of the $bc_1$ complex but not for single turnover $Q_o$ site catalysis. J Biol Chem *277*, 3471-3476.

Davidson, E., Ohnishi, T., Tokito, M., and Daldal, F. (1992). *Rhodobacter capsulatus* mutants lacking the Rieske FeS protein form a stable cytochrome $bc_1$ subcomplex with an intact quinone reduction site. Biochemistry *31*, 3351-3358.

Ditta, G., Schmidhauser, T., Yakobson, E., Lu, P., Liang, X.W., Finlay, D.R., Guiney, D., and Helinski, D.R. (1985). Plasmids related to the broad host range vector, pRK290, useful for gene cloning and for monitoring gene expression. Plasmid *13*, 149-153.

Dong, G.Q., Fan, H., Schneidman-Duhovny, D., Webb, B., and Sali, A. (2013). Optimized atomic statistical potentials: assessment of protein interfaces and loops. Bioinformatics *29*, 3158-3166.

Emsley, P., Lohkamp, B., Scott, W.G., and Cowtan, K. (2010). Features and development of Coot. Acta Crystallogr D Biol Crystallogr *66*, 486-501.

Gill, S.C., and von Hippel, P.H. (1989). Calculation of protein extinction coefficients from amino acid sequence data. Anal Biochem *182*, 319-326.

Gotze, M., Pettelkau, J., Fritzsche, R., Ihling, C.H., Schafer, M., and Sinz, A. (2015). Automated assignment of MS/MS cleavable cross-links in protein 3D-structure analysis. J Am Soc Mass Spectrom *26*, 83-97.

Gray, K.A., Grooms, M., Myllykallio, H., Moomaw, C., Slaughter, C., and Daldal, F. (1994). *Rhodobacter capsulatus* contains a novel *cb*-type cytochrome *c* oxidase without a $Cu_A$ center. Biochemistry *33*, 3120-3127.

Holden, H.M., Meyer, T.E., Cusanovich, M.A., Daldal, F., and Rayment, I. (1987). Crystallization and preliminary analysis of crystals of cytochrome $c_2$ from *Rhodopseudomonas capsulata*. J Mol Biol *195*, 229-231.

Jones, D.T., Tress, M., Bryson, K., and Hadley, C. (1999). Successful recognition of protein folds using threading methods biased by sequence similarity and predicted secondary structure. Proteins *Suppl 3*, 104-111.

Kalisman, N., Adams, C.M., and Levitt, M. (2012). Subunit order of eukaryotic TRiC/CCT chaperonin by cross-linking, mass spectrometry, and combinatorial homology modeling. Proc Natl Acad Sci U S A *109*, 2884-2889.

Khalfaoui-Hassani, B., Wu, H., Blaby-Haas, C.E., Zhang, Y., Sandri, F., Verissimo, A.F., Koch, H.G., and Daldal, F. (2018). Widespread Distribution and Functional Specificity of the Copper Importer CcoA: Distinct Cu Uptake Routes for Bacterial Cytochrome *c* Oxidases. MBio *9*.

Ko, J., Park, H., Heo, L., and Seok, C. (2012). GalaxyWEB server for protein structure prediction and refinement. Nucleic Acids Res *40*, W294-297.

Koch, H.G., Hwang, O., and Daldal, F. (1998). Isolation and characterization of *Rhodobacter capsulatus* mutants affected in cytochrome $cbb_3$ oxidase activity. J Bacteriol *180*, 969-978.

Kosinski, J., von Appen, A., Ori, A., Karius, K., Muller, C.W., and Beck, M. (2015). Xlink Analyzer: software for analysis and visualization of cross-linking data in the context of three-dimensional structures. J Struct Biol *189*, 177-183.

Laemmli, U.K. (1970). Cleavage of structural proteins during the assembly of the head of bacteriophage T4. Nature *227*, 680-685.

Liebschner, D., Afonine, P.V., Baker, M.L., Bunkoczi, G., Chen, V.B., Croll, T.I., Hintze, B., Hung, L.W., Jain, S., McCoy, A.J., *et al.* (2019). Macromolecular structure determination using X-rays, neutrons and electrons: recent developments in Phenix. Acta Crystallogr D Struct Biol *75*, 861-877.

Myllykallio, H., Jenney, F.E., Jr., Moomaw, C.R., Slaughter, C.A., and Daldal, F. (1997). Cytochrome $c_y$ of *Rhodobacter capsulatus* is attached to the cytoplasmic membrane by an uncleaved signal sequence-like anchor. J Bacteriol *179*, 2623-2631.

Ozturk, Y., Lee, D.W., Mandaci, S., Osyczka, A., Prince, R.C., and Daldal, F. (2008). Soluble variants of *Rhodobacter capsulatus* membrane-anchored cytochrome $c_y$ are efficient photosynthetic electron carriers. J Biol Chem *283*, 13964-13972.

Peters, A., Kulajta, C., Pawlik, G., Daldal, F., and Koch, H.G. (2008). Stability of the $cbb_3$-type cytochrome oxidase requires specific CcoQ-CcoP interactions. J Bacteriol *190*, 5576-5586.

Pettersen, E.F., Goddard, T.D., Huang, C.C., Couch, G.S., Greenblatt, D.M., Meng, E.C., and Ferrin, T.E. (2004). UCSF Chimera--a visualization system for exploratory research and analysis. J Comput Chem *25*, 1605-1612.

Poornam, G.P., Matsumoto, A., Ishida, H., and Hayward, S. (2009). A method for the analysis of domain movements in large biomolecular complexes. Proteins *76*, 201-212.

Punjani, A., Rubinstein, J.L., Fleet, D.J., and Brubaker, M.A. (2017). cryoSPARC: algorithms for rapid unsupervised cryo-EM structure determination. Nat Methods *14*, 290-296.

Rohou, A., and Grigorieff, N. (2015). CTFFIND4: fast and accurate defocus estimation from electron micrographs. J Struct Biol *192*, 216-221.

Sambrook, J., and Russell, D.W. (2001). Molecular Cloning: a laboratory manual, 3rd ed. edn (Cold Spring Harbor: Cold Spring harbor Laboratory Press).

Schneidman-Duhovny, D., Inbar, Y., Nussinov, R., and Wolfson, H.J. (2005). PatchDock and SymmDock: servers for rigid and symmetric docking. Nucleic Acids Res *33*, W363-367.

Selamoglu, N., Onder, O., Ozturk, Y., Khalfaoui-Hassani, B., Blaby-Haas, C.E., Garcia, B.A., Koch, H.G., and Daldal, F. (2020). Comparative differential cuproproteomes of *Rhodobacter capsulatus* reveal novel copper homeostasis related proteins. Metallomics *12*, 572-591.

Sheng, Q., Li, R., Dai, J., Li, Q., Su, Z., Guo, Y., Li, C., Shyr, Y., and Zeng, R. (2015). Preprocessing significantly improves the peptide/protein identification sensitivity of high-resolution isobarically labeled tandem mass spectrometry data. Mol Cell Proteomics *14*, 405-417.

Slavin, M., and Kalisman, N. (2018). Structural Analysis of Protein Complexes by Cross-Linking and Mass Spectrometry. Methods Mol Biol *1764*, 173-183.

Soding, J. (2005). Protein homology detection by HMM-HMM comparison. Bioinformatics *21*, 951-960.

Tang, G., Peng, L., Baldwin, P.R., Mann, D.S., Jiang, W., Rees, I., and Ludtke, S.J. (2007). EMAN2: An extensible image processing suite for electron microscopy. J Struct Biol *157*, 38-46.

Thomas, P.E., Ryan, D., and Levin, W. (1976). An improved staining procedure for the detection of the peroxidase activity of cytochrome P-450 on sodium dodecyl sulfate polyacrylamide gels. Anal Biochem *75*, 168-176.

Valkova-Valchanova, M.B., Saribas, A.S., Gibney, B.R., Dutton, P.L., and Daldal, F. (1998). Isolation and characterization of a two-subunit cytochrome $b$-$c_1$ subcomplex from *Rhodobacter capsulatus* and reconstitution of its ubihydroquinone oxidation ($Q_o$) site with purified Fe-S protein subunit. Biochemistry *37*, 16242-16251.

Webb, B., and Sali, A. (2014). Comparative Protein Structure Modeling Using MODELLER. Curr Protoc Bioinformatics *47*, 5 6 1-32.

Williams, C.J., Headd, J.J., Moriarty, N.W., Prisant, M.G., Videau, L.L., Deis, L.N., Verma, V., Keedy, D.A., Hintze, B.J., Chen, V.B., *et al.* (2018). MolProbity: More and better reference data for improved all-atom structure validation. Protein Sci *27*, 293-315.

Wittig, I., Braun, H.P., and Schagger, H. (2006). Blue native PAGE. Nat Protoc *1*, 418-428.

Yan, L.J., and Forster, M.J. (2009). Resolving mitochondrial protein complexes using nongradient blue native polyacrylamide gel electrophoresis. Anal Biochem *389*, 143-149.

Yang, J., Yan, R., Roy, A., Xu, D., Poisson, J., and Zhang, Y. (2015). The I-TASSER Suite: protein structure and function prediction. Nat Methods *12*, 7-8.

Yen, H.C., Hu, N.T., and Marrs, B.L. (1979). Characterization of the gene transfer agent made by an overproducer mutant of *Rhodopseudomonas capsulata*. J Mol Biol *131*, 157-168.

Zannoni, D., Prince, R.C., Dutton, P.L., and Marrs, B.L. (1980). Isolation and Characterization of a Cytochrome $c_2$-Deficient Mutant of *Rhodopseudomonas-capsulata*. Febs Lett *113*, 289-293.

Zeng, H., Wang, S., Zhou, T., Zhao, F., Li, X., Wu, Q., and Xu, J. (2018). ComplexContact: a web server for inter-protein contact prediction using deep learning. Nucleic Acids Res *46*, W432-W437.

Zhang, W., Yang, J., He, B., Walker, S.E., Zhang, H., Govindarajoo, B., Virtanen, J., Xue, Z., Shen, H.B., and Zhang, Y. (2016). Integration of QUARK and I-TASSER for Ab Initio Protein Structure Prediction in CASP11. Proteins *84 Suppl 1*, 76-86.

Zheng, S., Palovcak, E., Armache, J.-P., Verba, K.A., Cheng, Y., and Agard, D.A. (2017). MotionCor2: anisotropic correction of beam-induced motion for improved cryo-electron microscopy. Nat Methods *14*, 331-332.

Zivanov, J., Nakane, T., Forsberg, B.O., Kimanius, D., Hagen, W.J., Lindahl, E., and Scheres, S.H. (2018). New tools for automated high-resolution cryo-EM structure determination in RELION-3. eLife *7:e42166*.

**Table S1: Strains and plasmids used in this work.**

| Strain | Description | Phenotype | Reference |
|---|---|---|---|
| ***E. coli*** | | | |
| HB101 | F$^-$Δ(*gpt-proA*)62 *leuB6 supE44 ara*-14 *galK2 lacY* 1 Δ(*mcrC-mrr*) *rpsL20* (Str$^R$) *xyl-5 mtl-1recA*13 | Str$^r$ | (Sambrook and Russell, 2001) |
| XL1-Blue | *recA1 endA1 gyrA96 thi-1 hsdr-17 supE44 relA1 lac [F' proAB lacI q ΔM15 Tn10]* | Tet$^r$ | Stratagene |
| ***R. capsulatus*** | | | |
| [a]MT1131 | *crtD121*Rif$^r$ | Wild Type | (Zannoni et al., 1980) |
| MG1 | *ccoP::kan* | Kan$^r$, Cox$^-$, NADI$^-$ | (Koch et al., 1998) |
| YO12 | *ccoP::kan Δ*(petABC::Gm) | Kan$^r$, Cox$^-$, NADI$^-$, Gm$^r$, Ps$^-$ | This work |
| M7G-CBC1 | *crtDJ21 cox-410 Δ(petBC::spe)18* | Spe$^r$, Nadi$^-$, Myx$^r$ | (Daldal, 1988) |
| F2J-R4 | *crtD121 Δ(cycA::kan) Δ(cycY::spe) X4* | cyt $c_2^-$, cyt $c_y^-$, X$^-$, Ps$^-$, Res$^+$ | (Ozturk et al., 2008) |

| Plasmid | Description | Phenotype | Reference |
|---|---|---|---|
| pBSII | pBluescriptII (KS+) | Amp$^r$ | Stratagene |
| pRK2013 | Conjugation helper | Kan$^r$ | (Ditta et al., 1985) |
| pRK415 | Broad host-range vector | Tet$^r$ | (Ditta et al., 1985) |
| pMTS1 | *petABC* on pRK415 derivative with *kan* instead of *tet* | Kan$^r$ | (Gray et al., 1994) |
| pOX15 | *ccoNOQP* on pRK415 | | |

| | | | |
|---|---|---|---|
| pYO34 | *Δ(petABC::gm)* | $Gm^r$ | (Ozturk et al., 2008) |

| Plasmid | Description | Phenotype | Reference |
|---|---|---|---|
| pYO60 | 1.13 kb StuI/HindIII fragment with *ccoP-His*$_8$ cloned to StuI/HindIII digested pMTSI to yield *petABC::ccoP-His*$_8$ | $Kan^r$ | This work |
| pYO63 | *petABC::ccoP-His*$_8$ fusion on 3.9 kb BamHI fragment of pYO60 cloned to pBSII plasmid | $Amp^r$ | This work |
| pYO76 | *petABC::ccoP-His*$_8$ fusion on 3.9 kb BamHI fragment of pYO60 cloned to pRK415 plasmid | $Tet^r$ | This work |
| pYO77 | *petABC::ccoP-His*$_8$ fusion linked by 8 AA (**GGSGGGSG)** linker on pBSII plasmid | $Amp^r$ | This work |
| pYO78 | *petABC::ccoP-His*$_8$ fusion linked by 12 AA (**GGSGGGSGGGSG)** linker on pBSII plasmid | $Amp^r$ | This work |
| pYO79 | *petABC::ccoP-His*$_8$ fusion linked by 12 AA (**ASIAGGRTASGP)** linker on pBSII plasmid | $Amp^r$ | This work |
| pYO80 | *petABC::ccoP-His*$_8$ fusion linked by 8 AA (**GGSGGGSG)** linker on pRK415 plasmid | $Tet^r$ | This work |
| pYO81 | *petABC::ccoP-His*$_8$ fusion linked by 12 AA (**GGSGGGSGGGSG)** linker on pRK415 plasmid | $Tet^r$ | This work |
| pYO82 | *petABC::ccoP-His*$_8$ fusion linked by 12 AA (**ASIAGGRTASGP)** linker on pRK415 plasmid | $Tet^r$ | This work |
| pYO91 | 619 bp BalI/HinIII fragment containing *cycY-Flag* (without membrane anchor) cloned to BalI/HindIII fragment of pYO63 to yield *petABC::ccoP::cycY-Flag* | $Amp^r$ | This work |
| pYO92 | 4.5 kb KpnI/XbaI fragment of pYO91 containing *petABC::ccoP::cycY-Flag* cloned to pRK415 | $Tet^r$ , cyt $bc_1$-CcoP-$c_y$ | This work |
| pYO135 | 1.2 kb KpnI-BamHI fragment containing the signal sequence of *cycA* and the cyt *c* domain of *cycY* with K19R and H53Y mutations cloned to pRK415 | $Tet^r$, cyt S-$c_y$ K19R H53Y | (Ozturk et al., 2008) |

[a]*R. capsulatus* strain MT1131 is derived from SB1003 in multiple steps, as described in (Zannoni et al., 1980) ; first, a Ps-deficient mutant (TL1) was obtained using tetracycline suicide, then its *crtD* derivative was constructed by GTA cross to yield MT113, which then was used to yield its Ps-proficient derivative via a second GTA cross.

**Table S2. Primers used in this work**

| Primer name | Sequence |
|---|---|
| Fw-ccoP (StuI) | 5’-AAGGAGGCGAGGCC***TTG***AGCAAAAAACCGACGAC-3’ |
| Rv-ccoP (HindIII) | 5’-TTGCAACTGAAGCTTAAAGCTATATTGCCGATGC-3’ |
| [a]Fw-ccoPseq | 5’-GGCATCCGCGATCCGCTCG-3’ |
| [a]Rv-ccoPseq | 5’-CAGATGCGCGGGCATGTTC-3’ |
| [b]F-L2 | 5’-AAAAGGCC***TCG***GGCGGCTCGGGCGGCGGCTCG GGCAGCAAAAAACCGACGACCAAA-3’ |
| [b]F-L3 | 5’-AAAAGGCC***TCG***GGCGGCTCGGGCGGCGGCTCGGGCGGCGGCTCGGGCAGCAAAAAACCGAC-GACCAAA-3’ |
| [b]F-L4 | 5’-AAAAGGCC***TCG***GCGTCGATCGCCGGCGGCCGCACCGCATCGGGCCCGAGCAAAAAACCGAC-GACCAAA-3’ |
| Fw-cp-cy (BalI) | 5’-CTTTCTGGCCAGCTCTTTGCGACGCGGCCGGC-3’ |
| Rv-cp-cy (BglII+HinIII) | 5’-GGGGCAAGCTTGCAAAGATCTGAGGGCAAAAGG-3’ |

[a]Fw-ccoPseq and Rv-ccoPseq are used for sequencing and the rest of primers for constructions, as described in Methods.
[b]F-L denotes the primers used for inserting the linker sequences (in gray) into the *petC-ccoP* fusion junction. Underlined sequences indicate the restriction sites used for cloning. Italic bold ***TCG*** sequence indicates the stop codon of cyt $c_1$ changed to a serine codon.

**Table S3: Identification of protein subunits by mass spectrometry.**

| Protein | Cofactors | AA | $M_r$ | TMH | PSM | uniqPep | Identified Peptides | $X_{corr}$ | z |
|---|---|---|---|---|---|---|---|---|---|
| **cyt *c*₁** | heme $c_1$ | 258 | 28 | 1 | 60 | 9 | AGFSGPAGSGM*NQLFK | 4.71 | 2 |
| (in $c_1$-CcoP) | | | | | | | ETDMFPTR | 1.68 | 1 |
| | | | | | | | EYAAGLDTIIDK | 3.58 | 2 |
| | | | | | | | EYAAGLDTIIDKDSGEER | 5.00 | 2 |
| | | | | | | | EYAAGLDTIIDKDSGEERDR | 3.24 | 3 |
| | | | | | | | TFQIGGVPDTCK | 3.50 | 2 |
| | | | | | | | TFQIGGVPDTCKDAAGVK | 3.86 | 3 |
| | | | | | | | TLADDGGPQLDPTFVR | 4.50 | 2 |
| | | | | | | | VGDGM*GPDLSVM*AK | 3.94 | 2 |
| **CcoP** | heme $c_{p1}$ | 297 | 32 | 1 | 31 | 5 | ATAGQQVFADNC*VSC*HGEDAK | 3.73 | 3 |
| (in $c_1$-CcoP) | heme $c_{p2}$ | | | | | | AVEDKLVATDLTAIAADPELVTYTR | 3.94 | 3 |
| | | | | | | | EVQTTGHSWDGIEELNTPLPR | 5.42 | 2 |
| | | | | | | | KEVQTTGHSWDGIEELNTPLPR | 5.31 | 3 |
| | | | | | | | LVATDLTAIAADPELVTYTR | 6.07 | 3 |
| **CcoN** | heme *b* | 532 | 59 | 12 | 16 | 3 | DEYFDGVIR | 2.69 | 2 |
| | heme $b_3$ | | | | | | EVDAQGFLVNGFADTVGAK | 5.38 | 2 |
| | | | | | | | TM*GDAKPSKDEYFDGVIR | 3.83 | 3 |
| **cyt *b*** | heme $b_H$ | 437 | 49 | 8 | 20 | 5 | ADAEKDTLPFWPYFVIK | 3.92 | 3 |
| | heme $b_L$ | | | | | | DTLPFWPYFVIK | 3.28 | 2 |
| | | | | | | | DVNGGWAMR | 3.39 | 2 |
| | | | | | | | GLYYGSYK | 2.05 | 1 |
| | | | | | | | LPIVGLVYDTIM*IPTPK | 5.40 | 2 |
| **CcoO** | heme $c_o$ | 242 | 27 | 1 | 23 | 4 | ADFVAQADPNADSATLVANYGEK | 5.45 | 2 |
| | | | | | | | VEGM*RPYTPLELTGR | 3.64 | 2/3 |
| | | | | | | | VSTDALVGVPYSAEM*IAAAK | 4.61 | 2/3 |
| | | | | | | | YGHYSLAAESM*YDHPFQWGSK | 4.61 | 3 |

| Protein | Cofactors | AA | $M_r$ | TMH | PSM | uniqPep | Identified Peptides | $X_{corr}$ | z |
|---|---|---|---|---|---|---|---|---|---|
| **FeS protein** | 2Fe2S | 191 | 20 | 1 | 9 | 3 | IRKGPAPRNLDIPVAAFVDETTIK | 2.73 | 3 |
| (Rieske) | | | | | | | NLDIPVAAFVDETTIK | 4.37 | 2 |
| | | | | | | | SGDFGGWFC*PC*HGSHYDSAGR | 3.74 | 3 |
| **cyt $c_1$** | heme $c_1$ | 258 | 28 | 1 | 27 | 12 | TLADDGGPQLDPTFVR | 4.59 | 2 |
| (in $c_1$-CcoP-$c_y$) | | | | | | | AGFSGPAGSGM*NQLFK | 4.38 | 2 |
| | | | | | | | EYAAGLDTIIDKDSGEER | 3.52 | 2 |
| | | | | | | | EYAAGLDTIIDKDSGEERDRK | 3.34 | 4 |
| | | | | | | | VGDGM*GPDLSVM*AK | 3.26 | 2 |
| | | | | | | | TFQIGGVPDTC*K | 3.00 | 2 |
| | | | | | | | EYAAGLDTIIDKDSGEERDR | 2.94 | 4 |
| | | | | | | | EYAAGLDTIIDK | 2.68 | 2 |
| | | | | | | | DRKETDMFPTR | 2.66 | 3 |
| | | | | | | | KETDMFPTR | 2.32 | 2 |
| | | | | | | | ETDM*FPTR | 2.21 | 2 |
| | | | | | | | YDQAQLR | 2.10 | 2 |
| **CcoP** | heme $c_{p1}$ | 297 | 32 | 1 | 16 | 5 | ATAGQQVFADNC*VSC*HGEDAK | 3.45 | 3 |
| (in $c_1$-CcoP-$c_y$) | heme $c_{p2}$ | | | | | | ADVEKDIAK | 3.13 | 2 |
| | | | | | | | ISGQPADEAR | 2.82 | 2 |
| | | | | | | | NAGAAVFR | 2.56 | 2 |
| | | | | | | | LSEAQIR | 2.08 | 2 |
| **cyt $c_y$** | heme $c_y$ | 199 | 21 | 1 | 14 | 7 | M*SFVGLPEAADR | 3.70 | 2 |
| (in $c_1$-CcoP-$c_y$) | | | | | | | ATATVEGFKYSTAM*K | 3.21 | 3 |
| | | | | | | | ANVIAYLNTLPR | 3.03 | 2 |
| | | | | | | | ALLPSVDEAAM*PAK | 2.81 | 2 |
| | | | | | | | LDIYLVSPK | 2.41 | 2 |
| | | | | | | | NHVGNWTPER | 2.27 | 2 |
| | | | | | | | ATATVEGFK | 2.20 | 2 |

All proteins were identified using the bipartite SC preparation, except the $c_1$-CcoP-$c_y$ fusion peptide, which was identified using the tripartite SC. The numbers of peptide spectral matches (PSM) and unique peptides (uniqPep) are given for each protein. The sequences of the identified peptides are listed along with their X-correlation ($X_{corr}$) and charges (z) (Selamoglu et al., 2020). Some peptides were identified with and without various modifications; residues that were found in the modified version are marked with an asterisk; modifications were oxidation in the case of methionine (M*) and carbamidomethylation in the case of cysteine (C*). Additional information provided for each protein subunit includes the number of amino acid residues (AA), molecular mass ($M_r$) in kDa, number of transmembrane helices (TMH) and the cofactors content.

**Table S4.** Distance constraints between CcoH and CcoN defined by RaptorX-Complex analysis.

| CcoH Residue | CcoN Residue |
|---|---|
| 13 | 369 |
| 14 | 369 |
| 15 | 362 |
| 15 | 365 |
| 15 | 366 |
| 15 | 369 |
| 17 | 369 |
| 17 | 372 |
| 24 | 492 |
| 25 | 383 |
| 28 | 383 |
| 28 | 489 |
| 29 | 383 |
| 29 | 454 |
| 32 | 388 |
| 33 | 388 |
| 115* | 386 |

Of all the subunits of $CIII_2CIV$, reliable protein-protein interactions (> 0.5 score) were found by RaptorX-Complex only between CcoH and CcoN of CIV. The minimum and maximum distances between the listed residues were set to 1 and 15Å, respectively, and these constraints were used for docking CcoH to CIV at the interface of $CIII_2CIV$ via PatchDock (Schneidman-Duhovny et al., 2005). *this residue is outside of the modelled TMH of CcoH and not used for docking.

**Table S5.** Cross links identified in samples treated with DMTMM.

**Intra-Subunit Cross Links**

| Protein 1 | Protein 2 | Peptide 1 | Peptide 2 | Absolute Position 1 | Absolute Position 2 |
|---|---|---|---|---|---|
| FeS protein | FeS protein | G**K**PVFIR | **D**IELAR | 70 | 81 |
| FeS protein | FeS protein | DTSAENANKPGA**E**ATDENR | DE**K**DIELAR | 107 | 80 |
| FeS protein | FeS protein | DTSAENANKPGA**E**ATDENR | **K**GPAPR | 107 | 168 |
| FeS protein | FeS protein | SG**D**FGGWFC*PC*HGSHYDSAGR | **K**GPAPR | 147 | 168 |
| cyt $c_1$ | cyt $c_1$ | N**S**NVPDHAFSFEGIFGK | K**E**TDMFPTR | 2 | 84 |
| cyt $c_1$ | cyt $c_1$ | N**S**NVPDHAFSFEGIFGK | VGDGMGP**D**LSVM*AK | 2 | 99 |
| cyt $c_1$ | cyt $c_1$ | **E**YAAGLDTIIDKDSGEER | **#**NSNVPDHAFSFEGIFGK | 63 | -1 |
| cyt $c_1$ | cyt $c_1$ | EYAAGLDTIID**K**DSGEER | **E**TDMFPTR | 74 | 84 |
| cyt $c_1$ | cyt $c_1$ | **D**RKETDMFPTR | DR**K**ETDMFPTR | 81 | 83 |
| cyt $c_1$ | cyt $c_1$ | VGDGMGPDLSVM*A**K**AR | KET**D**M*FPTR | 105 | 86 |
| CcoP | CcoP | EVQTTGHSWDGIE**E**LNTPLPR | **K**PTTK | 23 | 4 |
| CcoP | CcoP | ADVEK**D**IAKFAEM*NK | ADVE**K**DIAKFAEMNK | 76 | 75 |
| CcoP | CcoP | FAEMN**K**AVEDK | ADV**E**KDIAK | 85 | 74 |
| cyt $c_2$ | cyt $c_2$ | GA**K**TGPNLYGVVGR | TAGTYP**E**FK | 32 | 50 |

| | | | | | |
|---|---|---|---|---|---|
| cyt $c_2$ | cyt $c_2$ | Y**K**DSIVALGASGFAWTEEDIATYVK | **D**PGAFLK | 54 | 78 |
| cyt $c_2$ | cyt $c_2$ | YK**D**SIVALGASGFAWTEEDIATYVK | A**K**TGMAFK | 55 | 93 |
| cyt $c_2$ | cyt $c_2$ | YKDSIVALGASGFAWTE**E**DIATYVK | TGMAF**K**LAK | 70 | 99 |
| cyt $c_2$ | cyt $c_2$ | YKDSIVALGASGFAWTEE**D**IATYVK | DPGAFL**K**EKLDDKK | 71 | 84 |
| cyt $c_2$ | cyt $c_2$ | DPGAFL**K**EK | **E**KLDDK | 84 | 85 |
| cyt $c_2$ | cyt $c_2$ | **E**KLDDKK | A**K**TGMAFK | 85 | 93 |
| cyt $c_2$ | cyt $c_2$ | TGMAF**K**LAK | **D**PGAFLK | 99 | 78 |
| cyt $c_2$ | cyt $c_2$ | TGM*AF**K**LAK | **E**KLDDKK | 99 | 85 |
| cyt $c_2$ | cyt $c_2$ | LA**K**GGEDVAAYLASVVK | **E**KLDDKK | 102 | 85 |
| cyt $c_2$ | cyt $c_2$ | GGE**D**VAAYLASVVK | TGM*AF**K**LAK | 106 | 99 |

**Inter-Subunit Cross Links (Intra- and Inter-Complex)**

| Protein 1 | Protein 2 | Peptide 1 | Peptide 2 | Absolute Position 1 | Absolute Position 2 |
|---|---|---|---|---|---|
| Cyt $c_1$ | FeS protein | VG**D**GMGPDLSVMAK | **K**GPAPR | 94 | 168 |
| CcoO | CcoP | V**E**GM*RPYTPLELTGR | ADVE**K**DIAK | 47 | 75 |
| CcoO | CcoP | V**E**GM*RPYTPLELTGR | FAEMN**K**AVEDK | 47 | 85 |
| FeS protein | CcoP | **D**TSAENANKPGAEATDENR | GGVMPSWSWAADGA**K**PR | 95 | 275 |
| FeS protein | CcoP | DTSAENANKPGA**E**ATDENR | FAEMN**K**AVEDK | 107 | 85 |

**Cyt $c_2$ and cyt $c_y$ Cross Links**

| Protein 1 | Protein 2 | Peptide 1 | Peptide 2 | Absolute Position 1 | Absolute Position 2 |
|---|---|---|---|---|---|
| cyt $c_1$ | cyt $c_2$ | EYAAGLDTIIDKDSGEER | AKTGMAFK | 63 | 93 |
| cyt $c_1$ | cyt $c_2$ | VGDGMGPDLSVMAK | AKTGMAFK | 94 | 93 |
| cyt $c_2$ | cyt $c_1$ | LAKGGEDVAAYLASVVK | EYAAGLDTIIDK | 102 | 63 |
| CcoP | cyt $c_2$ | LVATDLTAIAADPELVTYTR | LAKGGEDVAAYLASVVK | 95 | 116 |
| CcoO | cyt $c_2$ | VEGM*RPYTPLELTGR | AKTGMAFK | 47 | 93 |
| CcoO | cyt $c_2$ | VEGM*RPYTPLELTGR | TGM*AFKLAK | 47 | 99 |
| CcoO | cyt $c_2$ | VEGM*RPYTPLELTGR | LAKGGEDVAAYLASVVK | 47 | 102 |
| CcoO | cyt $c_2$ | ADFVAQADPNADSATLVANYGEK | AKTGMAFK | 191 | 93 |
| CcoO | cyt $c_2$ | ADFVAQADPNADSATLVANYGEK | LAKGGEDVAAYLASVVK | 191 | 102 |
| CcoO | cyt $c_2$ | ADFVAQADPNADSATLVANYGEK | GEKEFNK | 201 | 8 |
| CcoO | cyt $c_2$ | ADFVAQADPNADSATLVANYGEK | GAKTGPNLYGVVGR | 201 | 32 |
| CcoO | cyt $c_2$ | ADFVAQADPNADSATLVANYGEK | DPGAFLKEK | 201 | 84 |
| FeS protein | cyt $c_y$ | DTSAENANKPGAEATDENR | ATATVEGFKYSTAMK | 107 | 143 |
| Cyt $c_y$ | Cyt $c_1$ | ATATVEGFKYSTAMK | VGDGMGPDLSVMAK | 143 | 94 |
| Cyt $c_1$ | Cyt $c_y$ | #NSNVPDHAFSFEGIFGK | ALLPSVDEAAM*PAK | -1 | 54 |

| | | | | | |
|---|---|---|---|---|---|
| Cyt $c_1$ | Cyt $c_y$ | E**Y**AAGLDTIIDKDSGEER | ATATV**E**GFKYSTAMK | 64 | 140 |
| Cyt $c_1$ | Cyt $c_y$ | EYAAGLDTIIDKDSG**E**ER | ATATVEGF**K**YSTAMK | 78 | 143 |
| Cyt $c_1$ | Cyt $c_y$ | EYAAGLDTIIDKDSG**E**ER | AEVPGT**K**MSFVGLPEAADR | 78 | 175 |

Cross-linked amino acids are indicated in bold red letters. The absolute position 1 and 2 indicate the positions of the cross links with respect to the amino acid sequences of *Rhodobacter capsulatus* proteins as used in the PDB files (*i.e*. mature proteins); -1 refers to linkage to the N-ter $NH_2$ of the protein. Oxidized Met and alkylated Cys residues are noted with *, and N-terminal linked amino acids are noted with **#**. Uniprot IDs for the proteins are as follows: FeS protein, D5ANZ2; cyt $c_1$, D5ANZ4; CcoP, D5ARP7; CcoO, D5ARP5; cyt $c_2$, P00094 and cyt $c_y$, Q05389. The cross linked peptides listed are those identified by both MeroX (Gotze et al., 2015) and FindXL (Kalisman et al., 2012) search engines. Search parameters are provided in Methods.

**Table S6.** Cross links identified in samples treated with DSBU.

**Intra-Subunit Cross Links**

| Protein 1 | Protein 2 | Peptide 1 | Peptide 2 | Absolute Position 1 | Absolute Position 2 |
|---|---|---|---|---|---|
| FeS protein | FeS protein | DEKDIELAR | GKPVFIR | 80 | 70 |
| FeS protein | FeS protein | DTSAENANKPGAEATDENR | RDEKDIELAR | 103 | 80 |
| FeS protein | FeS protein | SVPLGALRDTSAENANKPGAEATDENR | KGPAPR | 103 | 168 |
| cyt $c_1$ | cyt $c_1$ | EYAAGLDTIIDKDSGEER | KETDMFPTR | 74 | 83 |
| cyt $c_1$ | cyt $c_1$ | EYAAGLDTIIDKDSGEER | VGDGMGPDLSVMAKAR | 74 | 105 |
| cyt $c_1$ | cyt $c_1$ | VGDGM*GPDLSVMAKAR | KETDMFPTR | 105 | 83 |
| cyt $c_1$ | cyt $c_1$ | TFQIGGVPDTC*KDAAGVK | VGDGMGPDLSVMAKAR | 168 | 105 |
| CcoP | CcoP | FAEMNKAVEDK | ADVEKDIAK | 85 | 75 |
| CcoP | CcoP | AVEDKLVATDLTAIAADPELVTYTR | ADVEKDIAKFAEMNK | 90 | 79 |
| CcoP | CcoP | GGVMPSWSWAADGAKPR | FAEMNKAVEDK | 275 | 85 |
| cyt $c_y$ | cyt $c_y$ | AEVPGTKMSFVGLPEAADR | ATATVEGFKYSTAMK | 175 | 143 |

**Inter-Subunit Cross Links (Intra-Complex)**

| Protein 1 | Protein 2 | Peptide 1 | Peptide 2 | Absolute Position 1 | Absolute Position 2 |
|---|---|---|---|---|---|

| | | | | | |
|---|---|---|---|---|---|
| FeS protein | cyt $c_1$ | RDEKDIELAR | KETDMFPTR | 80 | 83 |
| FeS protein | cyt $c_1$ | DTSAENANKPGAEATDENR | VGDGMGPDLSVMAKAR | 103 | 105 |
| cyt $c_1$ | FeS protein | EYAAGLDTIIDKDSGEER | GKPVFIR | 74 | 70 |
| cyt $c_1$ | FeS protein | EYAAGLDTIIDKDSGEERDRK | KGPAPR | 74 | 168 |
| cyt $c_1$ | FeS protein | DRKETDMFPTR | RDEKDIELAR | 83 | 80 |
| cyt $c_1$ | FeS protein | VGDGMGPDLSVMAKAR | RRDEKDIELAR | 105 | 80 |
| cyt $c_1$ | FeS protein | VGDGMGPDLSVMAKAR | KGPAPR | 105 | 168 |
| cyt $c_1$ | FeS protein | TFQIGGVPDTC*KDAAGVK | RDEKDIELAR | 168 | 80 |
| cyt $c_1$ | FeS protein | TFQIGGVPDTC*KDAAGVK | KGPAPR | 168 | 168 |

**Inter-Complex Cross Links (Intra-Supercomplex)**

| Protein 1 | Protein 2 | Peptide 1 | Peptide 2 | Absolute Position 1 | Absolute Position 2 |
|---|---|---|---|---|---|
| cyt $c_1$ | CcoP | DRKETDMFPTR | FAEMNKAVEDK | 83 | 85 |
| cyt $c_1$ | CcoP | DAAGVKITHGSWAR | ADVEKDIAK | 174 | 75 |
| CcoP | cyt $c_1$ | ADVEKDIAKFAEM*NK | KETDMFPTR | 75 | 83 |
| CcoP | cyt $c_1$ | ADVEKDIAKFAEMNK | VGDGMGPDLSVMAKAR | 79 | 105 |
| CcoP | cyt $c_1$ | FAEMNKAVEDK | KETDMFPTR | 85 | 83 |
| CcoP | cyt $c_1$ | GGVMPSWSWAADGAKPR | KETDMFPTR | 275 | 83 |
| CcoP | FeS protein | ADVEKDIAKFAEM*NK | KGPAPR | 75 | 168 |

| | | | | | |
|---|---|---|---|---|---|
| CcoP | FeS protein | DIA**K**FAEM*NK | **K**GPAPR | 79 | 168 |
| CcoP | FeS protein | FAEMN**K**AVEDK | **K**GPAPR | 85 | 168 |
| CcoP | FeS protein | GGVMPSWSWAADGA**K**PR | **K**GPAPR | 275 | 168 |

**Cyt $c_y$ Cross Links**

| Protein 1 | Protein 2 | Peptide 1 | Peptide 2 | Absolute Position 1 | Absolute Position 2 |
|---|---|---|---|---|---|
| cyt $c_y$ | FeS protein | IDG**K**NAVGPHLNGVIGR | **K**GPAPR | 121 | 168 |
| cyt $c_y$ | FeS protein | ATATVEGF**K**YSTAMK | RRDE**K**DIELAR | 143 | 80 |
| cyt $c_y$ | FeS protein | LDIYLVSP**K**AEVPGTK | **K**GPAPR | 168 | 168 |
| cyt $c_y$ | FeS protein | AEVPGT**K**M*SFVGLPEAADR | **K**GPAPR | 175 | 168 |

Cross-linked amino acids are indicated in bold red letters. The absolute position numbering of cross links corresponds to the amino acid sequences of *Rhodobacter capsulatus* proteins in PDB. Oxidized Met and alkylated Cys residues are noted with *. Uniprot IDs for proteins are: FeS protein, D5ANZ2; cyt $c_1$, D5ANZ4; CcoP, D5ARP7; cyt $c_2$, P00094; cyt $c_y$, Q05389. The cross linked peptides listed are those identified by both MeroX (Gotze et al., 2015) and MassAI (http://www.massai.dk) search engines; parameters are given in Methods.

**Table S7.** Validation of the CIV homology model fitted in maps SC-1A (EMD-22228; PDB: 6XKX), SC-1B (EMD-22230; PDB: 6XKZ) and SC-2A (EMD-2; PDB: 6XKW).

| | |
|---|---|
| Model composition | |
| Non-hydrogen atoms | 7330 |
| Protein residues | 915 |
| Heme groups | 5 |
| Cu atoms | 1 |
| MolProbity Score | 3.41 |
| Clash score | 119 |
| Rotamer outliers (%) | 5.75 |
| C-beta deviations | 34 |
| RMSD | |
| Bond lengths (Å) | 0.02 |
| Bond angles (°) | 2.52 |
| Ramachandran plot | |
| Outliers (%) | 1.00 |
| Favored (%) | 95.78 |

### Figure legends

**Figure S1. Characterization of *R. capsulatus* wild type and fusion SCs. A.** DDM-dispersed membranes from *R. capsulatus* wild-type strain MT1131 were separated in 4-13% BN-PAGE and the gel was overstained for CIV-in-gel activity (SI, Methods) to detect the ~440 kDa entity. Under the conditions used, CIV runs as a diffuse ~232 kDa band, just below the "green band" of bacteriochlorophyll-complexes. **B.** DDM-dispersed membranes from *R. capsulatus* wild-type strain MT1131(WT), and its mutant derivative YO12, lacking both CIII$_2$ and CIV but carrying plasmid pYO76 and expressing the bipartite SC (fusion), were separated in 4-16% BN-PAGE. One part of the gel was stained for CIV-in-gel activity (CIV-IGA, left), and the other part blotted onto a PVDF membrane and treated with *R. capsulatus* cyt *b* specific antibodies (□-cyt b, right), detecting multiple forms of CIII$_2$ running between ~250-300 kDa $M_r$. The size and position of HMW markers (GE Heathcare) are indicated, and arrows points out the SCs of ~440 kDa $M_r$. The intact SCs are not readily visible in a wild type strain due to their low amounts and fragile nature. **C**. Growth properties of *R. capsulatus* strain pYO76/YO12, lacking CIII$_2$ and CIV but carrying a bipartite SC (fusion). The top and bottom rows show, respectively, the photosynthetic growth (Ps) and the *in vivo* Cox activity (NADI-staining, Methods) observed under respiratory (Res) growth conditions of colonies from wild type (WT), YO12 lacking both CIII$_2$ and CIV (□CIII$_2$ □CIV), YO12 carrying a plasmid (pOX15) containing only CIV (*i.e.*, □CIII$_2$); and YO12 carrying a plasmid (pYO76) expressing the bipartite fusion SC (**Table S1**).

**Figure S2. Biochemical characterization of purified bipartite and tripartite SCs. A.** Redox difference spectra of the bipartite (Bipartite CIII$_2$CIV, left) and tripartite (Tripartite CIII$_2$CIV, right) SCs. Ascorbate-reduced *minus* ferricyanide-oxidized visible spectra (blue) revealing the *c*-

type cyts at 550 nm, and dithionite-reduced *minus* ferricyanide-oxidized spectra (red) revealing both *c*-type and *b*-type cyts at 550 and 560 nm, respectively, are shown. **B** and **C.** $DBH_2$: cyt *c* reductase ($CIII_2$) and cyt *c* oxidase (CIV) activities, respectively, of tripartite SCs monitored at 550 nm (Methods). Arrows indicate the additions of $DBH_2$, KCN and purified tripartite SCs as appropriate. Similar activities were also observed with the bipartite SCs. **D.** $DBH_2$ dependent $O_2$ consumption activity of tripartite SCs. Additions of $DBH_2$ and KCN are indicated by arrows as in **B** and **C**. Unlike the tripartite, the bipartite SCs shows no $O_2$ consumption activity in the absence of an external electron carrier cyt *c* (not shown).

**Figure S3. Classification tree for tripartite SCs.** Six datasets (dataset-1 to 6) were collected. One representative micrograph is shown on the top left corner with a white 30 nm scale bar. The datasets were processed by Relion 3.0 as appropriate. For simplicity, only the analyses leading to the best version of relevant maps are shown. In all cases the subclasses obtained in 3D classifications are shown in different colors, and both the class distribution (%) and nominal resolution (Å) are indicated. The masks are shown in orange (not scaled to maps) and the final maps deposited to EMDB in turquoise. After one or two rounds of 2D classification, best 2D class averages were retained (~500,000 particles) and classified into three to five 3D classes, using an initial model obtained from dataset 1 and lowpass-filtered to 60 Å (Box 3). **A.** Analysis of the datasets 1-4 and 6. All classes showing strong density for *cbb*$_3$-type CIV were retained and subjected to focused classification. A soft mask was wrapped around the CIV portion of SCs, information outside the mask was subtracted from the particle images, and particles inside the mask (mask1 in orange with subtract_mask1 in yellow) were sorted into 6 classes. The class showing a complete map of CIV and the highest level of detail was retained and subjected to 3D auto-refinement and post processing. The last two steps were repeated after CTF refinement and Bayesian polishing. The final map (SC-1A) contained 61,934 particles, and had a nominal resolution of 6.09 Å. **B.** A similar strategy was used for the analysis of datasets 1-5, except that a cylindrical mask around the CIV portion was used, and the CTF refinement and Bayesian polishing are omitted, as they did not improve the resolution. The final map (SC-1B) showed a different conformation of CIV relative to $CIII_2$ as compared to map SC-1A (**A**), contained 87,026 particles and had a nominal resolution of 7.2 Å. **C**. Two classes showing density corresponding to a second copy of CIV associated with $CIII_2$ were selected, and the particles subjected to another round of 2D classification. The class averages showing the highest level of details were retained, and the particles classified into three 3D classes, of which the class showing the highest occupancy of the second copy of CIV associated with $CIII_2$ was sorted into 6 subclasses using a solvent mask. The classes with the strongest density for the second copy of CIV were subjected to refinement and post processing, and one such class (SC-1C, in gray) is shown. Due to low number of particles in each class (~5,000) these maps could not be refined to high resolution.

**Figure S4. Different orientations of CIV relative to $CIII_2$ in tripartite SCs. A.** The $CIII_2$ structure (PDB: 6XI0) and the CIV homology model were fitted into the density maps SC-1A and SC-1B, side and top views are shown. Subunits are colored and labeled as in **Fig. 3A** and the monomers A and B of $CIII_2$ are indicated. The region corresponding to panel B is highlighted with a dashed box in monomer A. **B.** Fitting of the FeS-ED in both $CIII_2$ monomers A and B of map SC-1A (EMD-22228). $CIII_2$ models with their FeS-EDs in b- (PDB: 6XI0, yellow) or in c-positions (PDB: 6XKT, grey) were fitted into SC-1A. Although the resolution of the FeS-EDs is low, clearly, they are located closer to the b-position in SC-1A. The map is shown at the same sigma level in all panels for direct comparison, and the lover occupancy in monomer B is seen. **C.** The CIV homology and $CIII_2$ (PDB: 6XI0) models were fitted into SC1-A and SC-1B maps and the $CIII_2$ portions were superimposed to visualize the different orientations of CIV in the tripartite SCs. Only the helical and beta-sheets regions of the proteins are depicted as cylinders and arrows, respectively, with the coils omitted for clarity, and the lipid bilayer is indicated by horizontal lines. Red represents $CIII_2$ and CIV fitted into SC-1A and blue represents CIV fitted into SC-1B. The rotation/translation axis (green) transforming one orientation of CIV into the other one was determined using DynDom3D (dyndom.cmp.uea.ac.uk) (Poornam et al., 2009) with default parameters. **D**. Top view of only the CIV portions of SCs shown along the rotation/translation axis (green dot) and the 37° rotation between the orientations of CIV (colored as in **C**). **E**. Top view (perpendicular to the membrane plane) of the contour maps of tripartite $CIII_2CIV$. The surface contours of $CIII_2$ (right) and CIV (left) in map SC-1A (red), and the different orientation of CIV relative to $CIII_2$ in map SC-1B (blue) are shown. For comparison, the orientations of CIV with respect to $CIII_2$ in *S. cerevisiae* $CIII_2CIV$ (light gray, dashed line, PDB: 6GIQ) and in mycobacterial $CIII_2CIV_2$ (black dotted line, PDB: 6HWH) are shown after superimposition of the cyt *b* structures of all $CIII_2$.

**Figure S5. Binding of the cyt $c_y$, cyt S-$c_y$ and cyt $c_2$ to bipartite SC. A.** Binding of cyt *c* domain of $c_y$ (S-$c_y$, left) and full-length native cyt $c_y$ ($c_y$, right) to bipartite SC ($CIII_2CIV$). Purified SCs were mixed with cyt S-$c_y$ or cyt $c_y$ as indicated, and subjected to SEC. Samples before and after SEC on Superose 6 Increase were separated on 18% SDS-PAGE and silver stained. Note that only cyt $c_y$, but not S-$c_y$, co-eluted with bipartite $CIII_2CIV$. Purified samples of cyt $c_y$, cyt S-$c_y$ (indicated by arrows) and $CIII_2CIV$ used are shown for comparison. * and ** indicate the degradation product(s) of $c_1$-CcoP subunit of bipartite SCs and of cyt $c_y$, respectively. **B.** Binding of cyt $c_2$ to $cbb_3$-type CIV. Appropriate proteins were mixed, subjected to SEC on Superose 6 Increase, followed by SDS-PAGE and silver staining as in **A**. Proteins used are also shown for comparison. CIV was partially purified, its subunits are indicated by dashes, and the nature of the other bands, including that of ~10 kDa indicated by ?, are not known. Note that cyt $c_2$ (indicated by an arrow) coeluted with CIV.

**Figure S6. Classification tree for bipartite SCs and $CIII_2$ conformers.** A representative micrograph is shown at the bottom left corner with a white 30 nm scale bar. Three datasets were collected and particles were processed by Relion 3.0. Subclasses obtained in 3D classifications are shown in various colors, and labeled with class distribution (in %) and the nominal resolution as in **Fig. S3**. All masks are shown in orange, and the final maps that are deposited to EMDB in turquoise. **A.** An initial 2D classification of particles from dataset-1 yielded 376,748 particles in suitable class averages, which were subjected to a second round of 2D classification. Class-averages likely representing different orientations of bipartite SCs were classified into five 3D classes using the same initial model used for the tripartite SC analyses (**Fig. S3**), and a single class showing the entire SC at the highest resolution was selected. These particles were processed separately (not shown) as well as together with the SC particles from dataset-2 as in **B. B.** After two rounds of 2D classification of particles from dataset-2, those likely representing SCs in different orientations were retained, and classified into three 3D classes using the same initial model as in **A**. The class most closely resembling the overall shape of bipartite $CIII_2CIV$ was chosen, and subjected to another round of 2D classification. Only those particles that clearly represent the SCs were retained and combined with the particles from dataset-1 (**A**). Combined

SC particles from both datasets were classified into five 3D classes, the best of which was selected and further processed using a solvent mask. After 3D auto-refinement and post processing, a map containing 14,978 particles was obtained with a nominal resolution of 5.72 Å, which after per-particle CTF-refinement and Bayesian polishing led to map SC-2A at 5.18 Å resolution. **C.** Only the class averages resembling a $CIII_2$ dimer (without any bound CIV) were selected after the first round of 2D classification, and subjected to a second round of 2D classification after which appropriate class averages were selected, and a reference-free initial model was created in Relion 3.0. **D.** After two rounds of 2D classification of particles from dataset-2, those resembling a $CIII_2$ dimer were selected and classified into three 3D classes using the initial model created in (**C**). The class with the highest resolution and level of detail was selected and combined with the $CIII_2$ particles obtained in **C**. The combined $CIII_2$ particles were subjected to 3D classification into 5 classes and the best class was further processed, either imposing C2 symmetry (**E**) or assuming no symmetry (**F**). **E.** The class with the highest level of detail was selected and aligned particles were subjected to a second round of 3D classification without image alignment, using a solvent mask, and imposing C2 symmetry. Six classes were obtained, some of which showed both Rieske FeS protein external domain (FeS-ED) proteins in b, while other classes showed both FeS-ED proteins in c, positions. The class with the largest number of particles and the highest resolution was selected and subjected to several rounds of refinement, post processing, per-particle CTF-refinement and Bayesian polishing. The final map consisted of ~38,000 particles has a nominal resolution of 3.3 Å, and shows both FeS-ED proteins in b position. **F.** The same strategy as in (**E**) was used, but no symmetry (*i.e.*, C1 symmetry) was applied in the two rounds of 3D classification. As a consequence, besides both FeS-EDs being in the same conformation (b-b and c-c), a third state was observed with one FeS-ED in b, and the other in c position (b-c). This class representing the b-c conformational state was refined without imposing C2 symmetry, while the best classes representing the b-b and c-c conformational states were refined with C2 symmetry as in (**E**). Three distinct maps, representing the different conformations of FeS-ED proteins in $CIII_2$ were obtained. **G.** Upon analysis of a third dataset, some maps showed an additional feature at the interface of CIII and CIV, which was tentatively attributed to cyt *c* domain of $c_y$. To obtain the best map containing the extra density, all classes showing extra density at the interface were combined after the second round of 3D classification using a soft mask and the $CIII_2$ portion of the particles was subtracted. The remaining particles were sorted into 6 classes in another round of 3D classification, using a mask that included CIV and the interface region. Resulting classes were subjected to 3D auto-refinement and post-processing, and the final map (SC-2B) showing the strongest density at the SC interface is shown.

**Figure S7. Local resolutions of the cryo-EM maps SC-2A and $CIII_2$. A** and **B**, local resolutions of SC-2A and $CIII_2$ maps, respectively. Local resolution was calculated in cryoSPARC (Punjani et al., 2017), and the resolution range is shown at the bottom of each map. **C** and **D**, Fourier Shell Correlations (FSC) for SC-2A and $CIII_2$ maps, respectively. The FSC plots for the unmasked (green), masked (blue), and corrected (red) maps are shown, and the nominal resolution estimates for the corrected maps at FSC = 0.143 and 0.5 are indicated.

**Figure S8. Docking of CcoH to *cbb*$_3$-type CIV. A**. The first 45 N-ter amino acid residues of *R. capsulatus* CcoH, with its predicted TMH (residues 12 to 34, underlined) is shown. Residues predicted to interact with CcoN TMH9 (**Table S4**) are highlighted in the same colors in which they are depicted in panel **C**. **B**. An *ab initio* model for this TMH portion of CcoH, generated by

I-TASSER server (Zhang et al., 2016), was docked to *cbb*$_3$-type CIV using Patchdock (Schneidman-Duhovny et al., 2005) guided with the distance restraints (**Table S4**) obtained from RaptorX-ComplexContact analysis (Zeng et al., 2018). A cluster of 25 representative orientations of CcoH models (depicted in a different colors) is shown on *R. capsulatus cbb*$_3$-type CIV, where the CcoN, CcoO and CcoP subunits are shown in gray. Views from two different angles are presented to better depict the location of CcoH TMH, near CcoN TMH9 (not labeled, see **Fig. 3C**, top view). The region shown in panel **C** is highlighted with a dashed box and the arrow indicates the refined position of CcoH TMH shown in panel **C**. **C.** Enlarged view showing predicted interactions between CcoH TMH and CcoN TMH9. CcoH TMH is shown in its final position as modelled in map SC-2A (EMD-22227; PDB: 6XKW), see **Fig. 4C,D**), cyt $c_y$ and some TMHs of CIV are not shown for better visibility. The entire structure is in grey, and the residues with predicted interactions are depicted in the same colors. Some overlapping interactions (**Table S4**) are omitted for clarity.

**Figure S9. XL-MS guided docking of cyt $c_2$ to CIII$_2$.** Subsets of protein-protein crosslinks obtained using XL-MS (**Tables S5** and **S6**) are shown among the SC protein subunits (colored as in **Fig. 3).** Cyt $c_2$ (orange, PDB: 1C2N) is shown on monomer **A** of *R. capsulatus* CIII$_2$, at the position defined by homology to the analogous yeast iso-1 cyt *c*-CIII$_2$ co-crystal structure (PDB: 3CX5) used as a template. **A.** Intra-subunit XLs obtained with DMTMM, and **B.** XLs between cyt $c_2$ and CIII$_2$ are shown. XLs conforming to the allowed maximum distance (25 or 30Å for DMTMM) are in blue with only one being slightly borderline (29Å) XL is in red. **C**. The binding site defined by *R. capsulatus* cyt $c_2$-CIII$_2$ co-crystal homology model (shown in **A** and **B**) is compared with the binding regions defined by docking cyt $c_2$ to CIII$_2$ using Patchdock guided by the DMTMM XLs used as distance restraints. The gray transparent volume, encompassing the binding regions of all cyt $c_2$ models docked on monomer A of $CIII_2$, was generated by choosing three representative models covering the entire range of all predicted cyt $c_2$ binding positions. **D**. Intra-subunit XLs identified using DSBU, all conforming to the maximum distance allowed (35Å), are shown in blue.

**Figure S10. Surface charge distribution of cyt $c_2$, cyt $c_y$, $CIII_2$ and CIV.** Protein surfaces were colored using the coulombic surface function in Chimera (positive charges in blue and negative charges in red). The water-exposed edge of cyt $c_2$ heme and heme $c_{p2}$ at its likely binding region on CIV (**A**), together with cyt $c_y$ heme and heme $c_1$ of $CIII_2$ (**B**) are colored as in **Fig. 7**, and indicated by arrows. These edges are barely visible at this scale, and to better visualize the surface charges on the periplasmic face of cyt $c_1$, a tilted front view of CIII is shown.

**Figure S11. Plausible location of cyt *c* domain of $c_y$ in $CIII_2CIV$.** The SC is shown fitted inside the transparent low resolution cryo-EM map SC-2B (**Fig. S6G**). Subunits are colored as in **Fig. 3**, except that $CIII_2$ monomer B is shown in grey. Twenty representative models of cyt *c* domain of $c_y$ docked onto $CIII_2$ monomer A are shown in different colors. **A** and **B** represent the back (as in **Fig. 4)** and front (as in **Fig. 3**) views of the SC. A region exhibiting extra electron density at the SC interface is indicated by an arrow. **C** shows a top view to better visualize the two partially overlapping clusters of cyt *c* domain of $c_y$ according to Patchdock, with one on cyt $c_1$ (arrow 1) and the other one at the interface of cyt $c_1$ and the FeS-ED protein (arrow 2) of $CIII_2$.

**Figure S12. Positions of the FeS-ED proteins affect docking of cyt *c* domain of $c_y$ on $CIII_2$.** Top and bottom rows show the XLs guided docking of cyt *c* domain of $c_y$ onto two $CIII_2$ conformers with their FeS-ED proteins in c ($CIII_2$ c-c) and in b ($CIII_2$ b-b) positions, respectively. **A.** One major docking cluster was observed per monomer of $CIII_2$ c-c, and sixteen models of cyt *c* domain of $c_y$ located on monomer A are shown. The subunits of monomer A are colored as in **Fig. 3** and those in monomer B shown in gray. **B** and **C** are the side and top views, respectively, of the semi-transparent map $CIII_2$-cc with its cofactors. The view shown here is similar to that of the SC in **Fig. 7A**, **C**. For clarity, the docking models of cyt *c* domain of $c_y$ are represented by their heme-Fe atoms (spheres). Unlike **A**, which shows only the clustered models, the Fe atoms of all models (28 total) docked on monomer A are shown either by fully colored red spheres for those located within 25 Å, or by semi-transparent spheres for those beyond 25 Å of cyt $c_1$ heme-Fe. **D**. Two major docking clusters were observed per monomer of $CIII_2$b-b, with one located on top of cyt $c_1$ (arrow 1 in **Fig. S11C**) and the other one at the inter-monomer interface between cyt $c_1$ and the FeS-ED protein of the other monomer (arrow 2 in **Fig. S11C**). Ten models of cyt *c* domain of $c_y$ representing the cluster located on top of cyt $c_1$ in monomer A are shown. **E** and **F** are the side and top views, respectively, of the map $CIII_2$-bb with its cofactors and the heme-Fe atoms of docked models of cyt *c* domain of $c_y$ (33 total) on monomer A, as in **B** and **C**. Comparison of the top and bottom rows shows that the docking models of cyt *c* domain of $c_y$ are clustered more tightly on top of cyt $c_1$ when the FeS-ED proteins of $CIII_2$ are in c position.

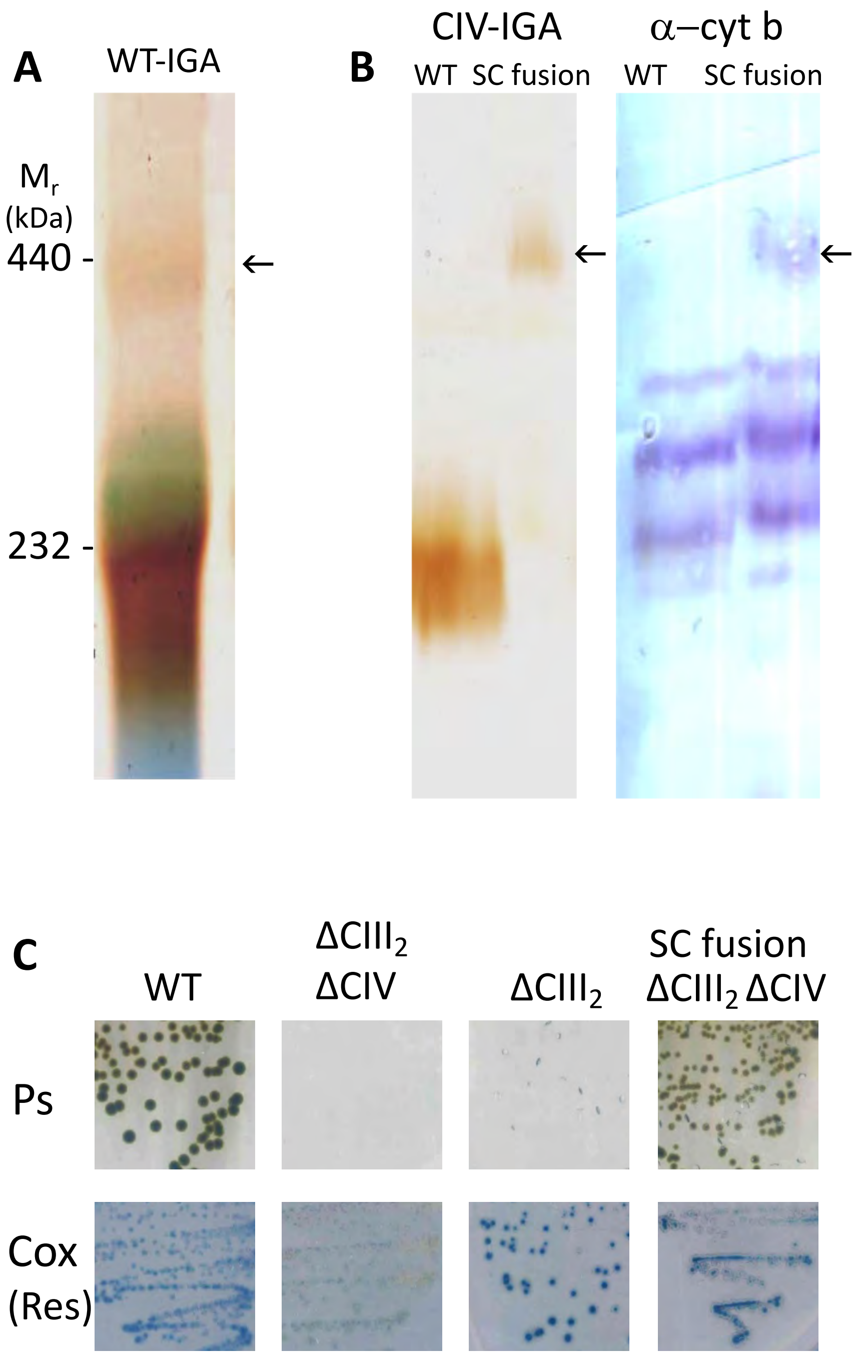

***Figure S1. Steimle et al.,***

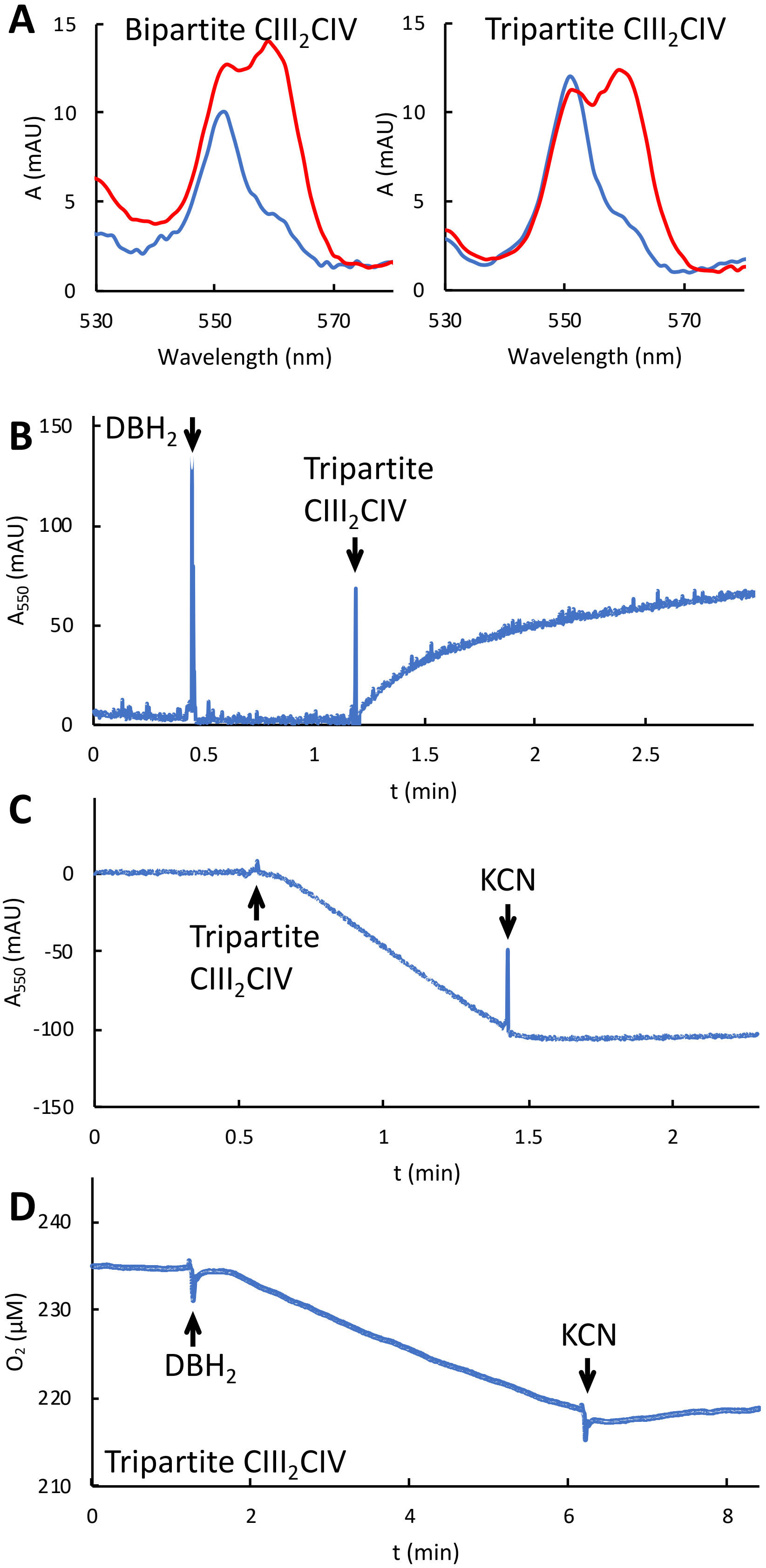

***Figure S2. Steimle et al.,***

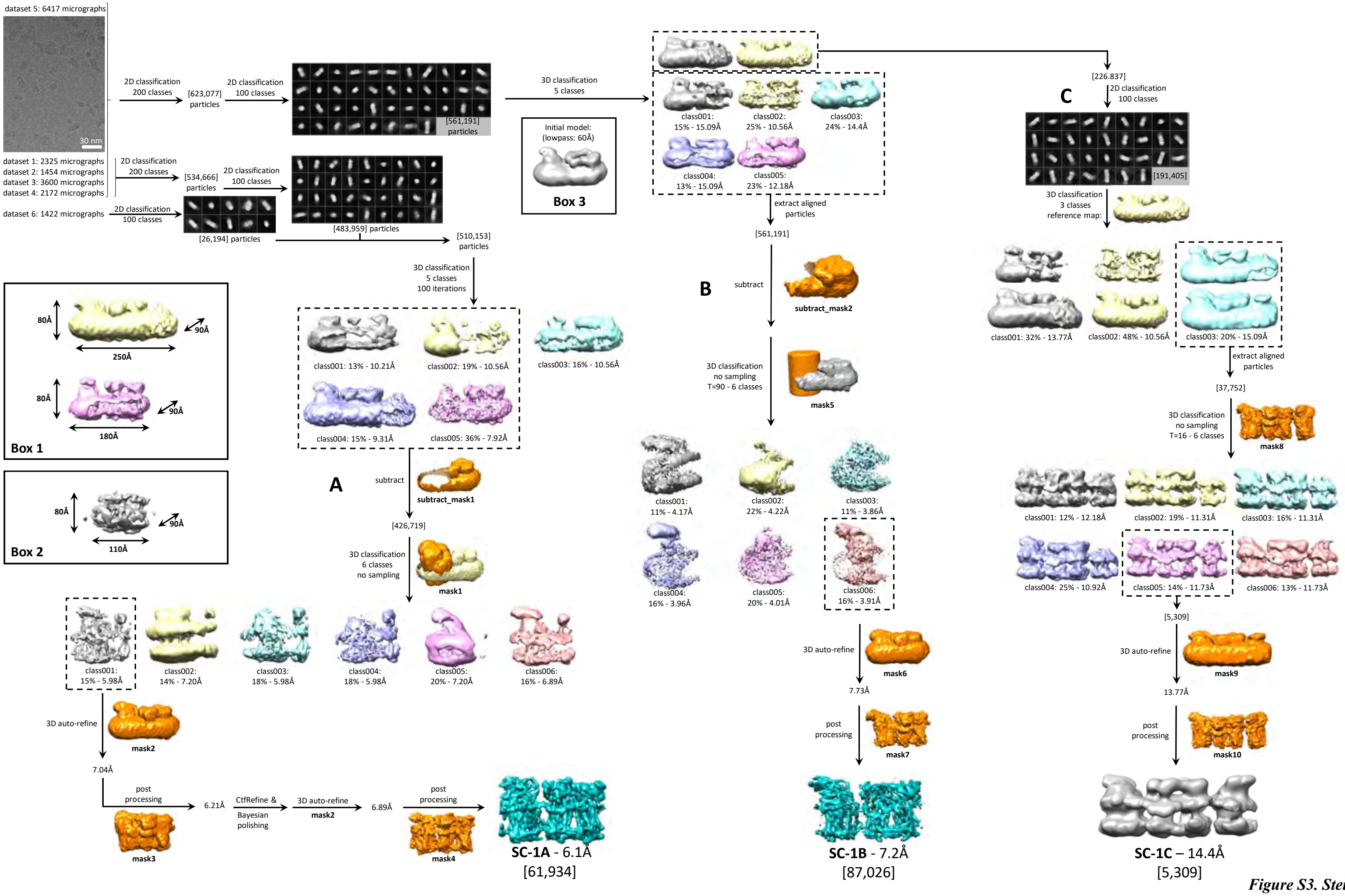

*Figure S3. Steimle et al.,*

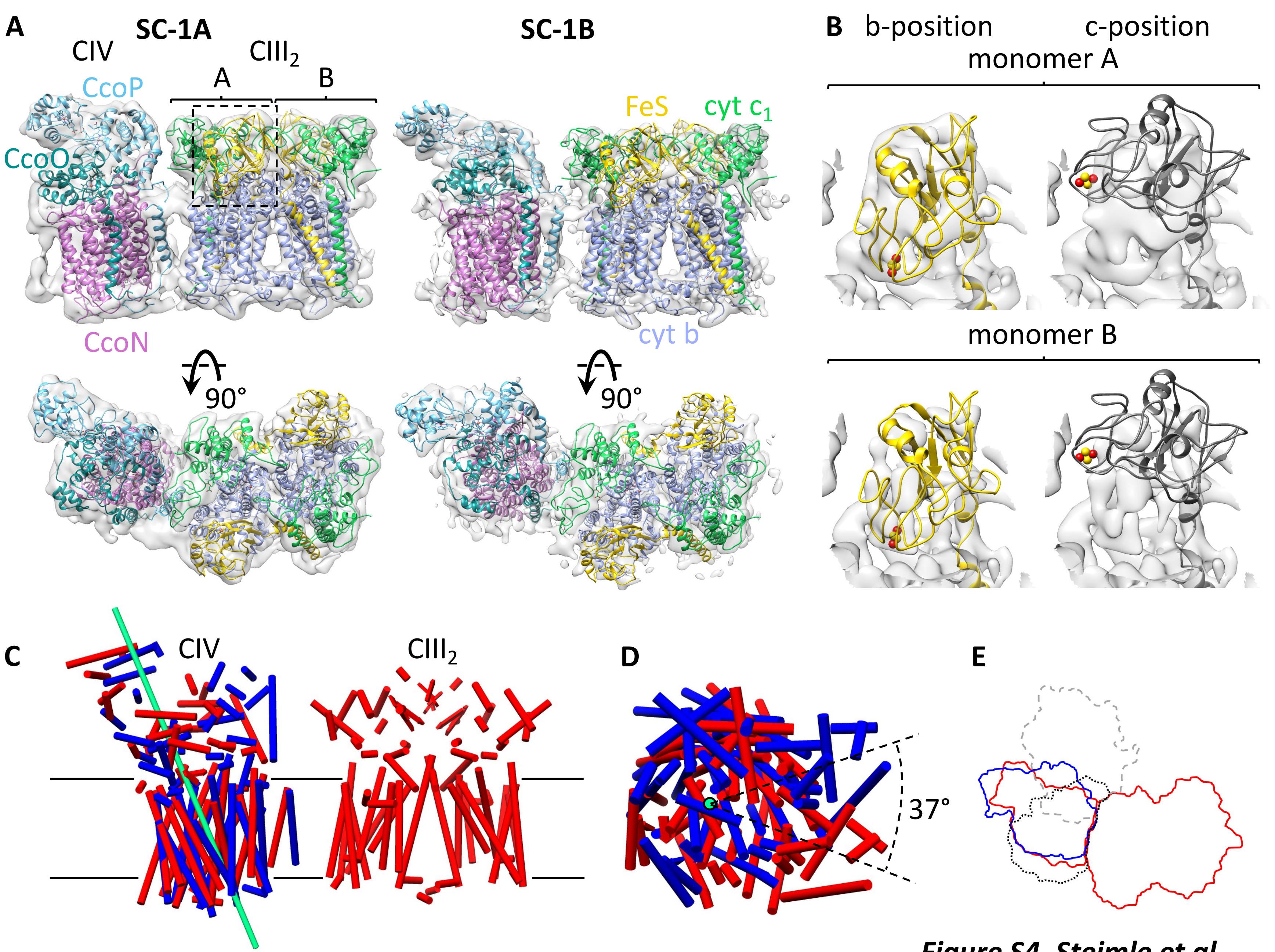

*Figure S4. Steimle et al.,*

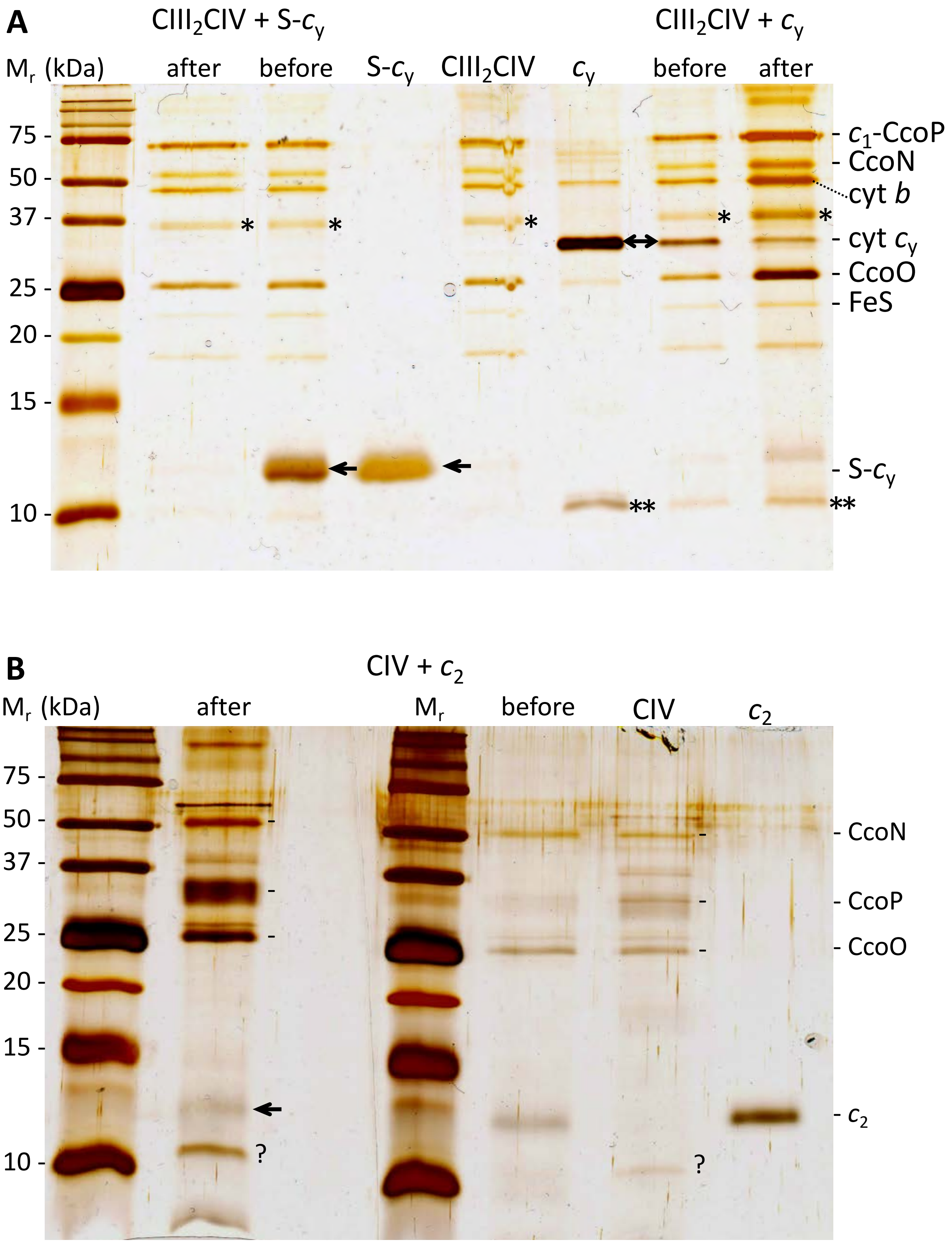

***Figure S5. Steimle et al.,***

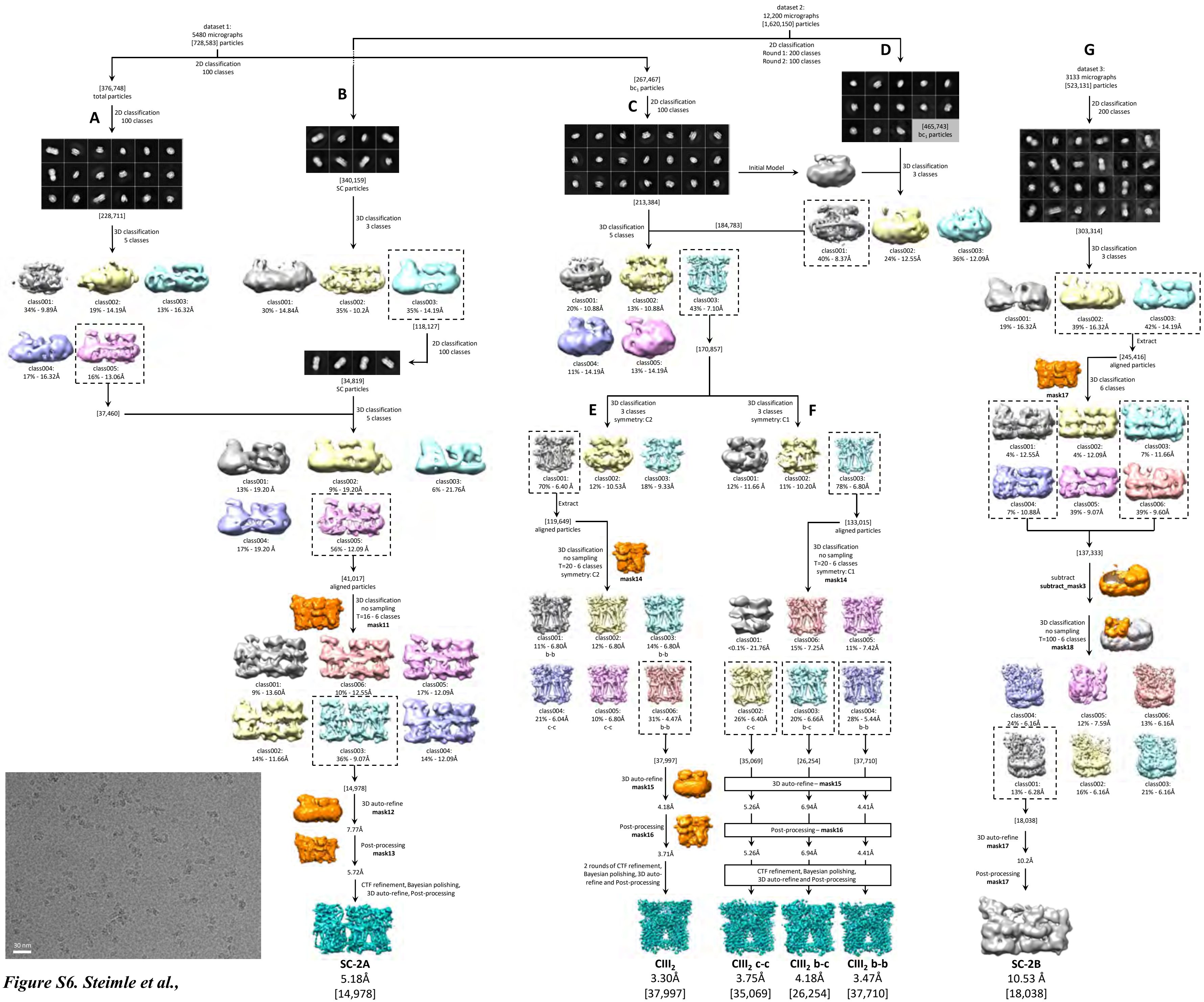

*Figure S6. Steimle et al.,*

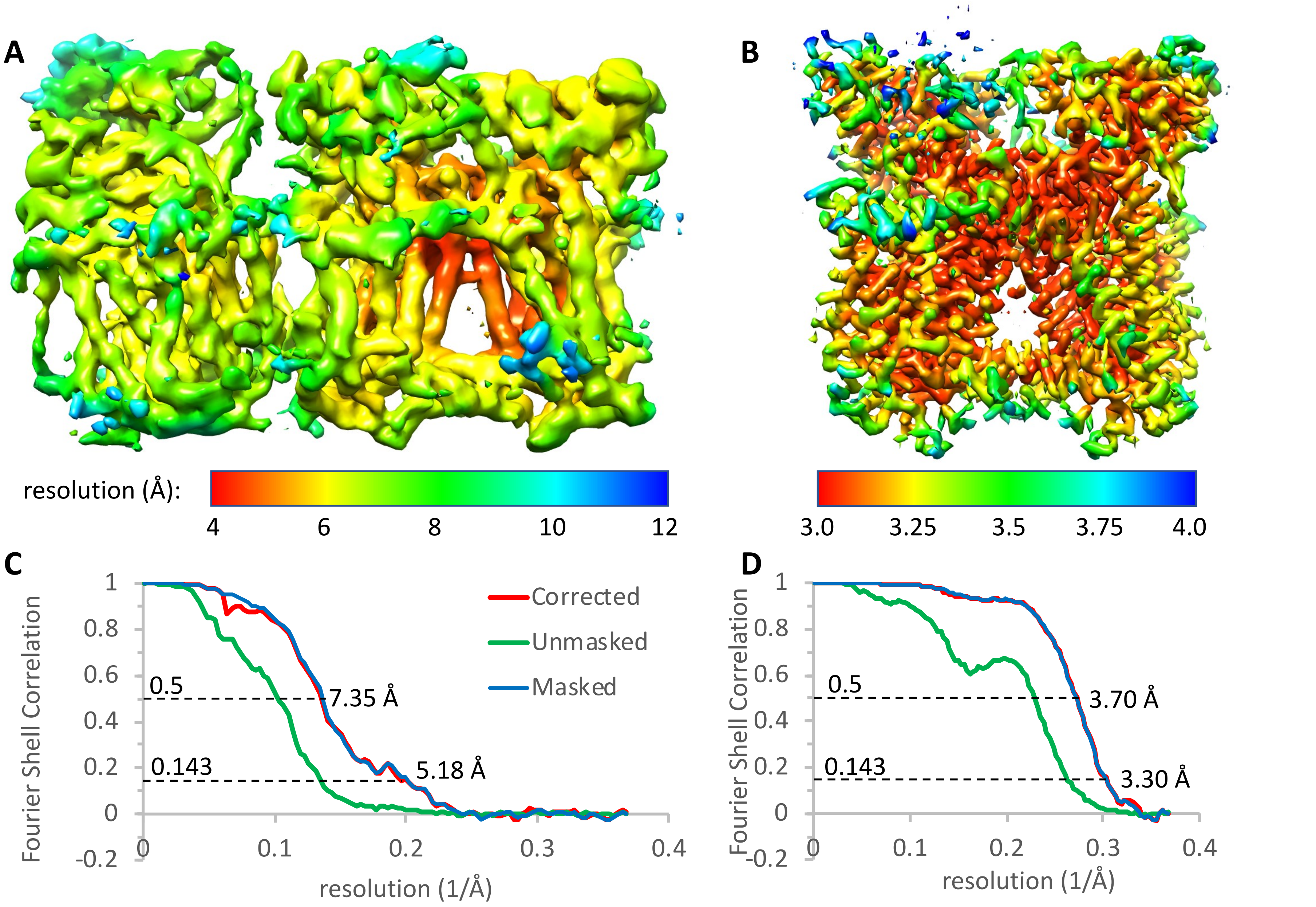

***Figure S7. Steimle et al.,***

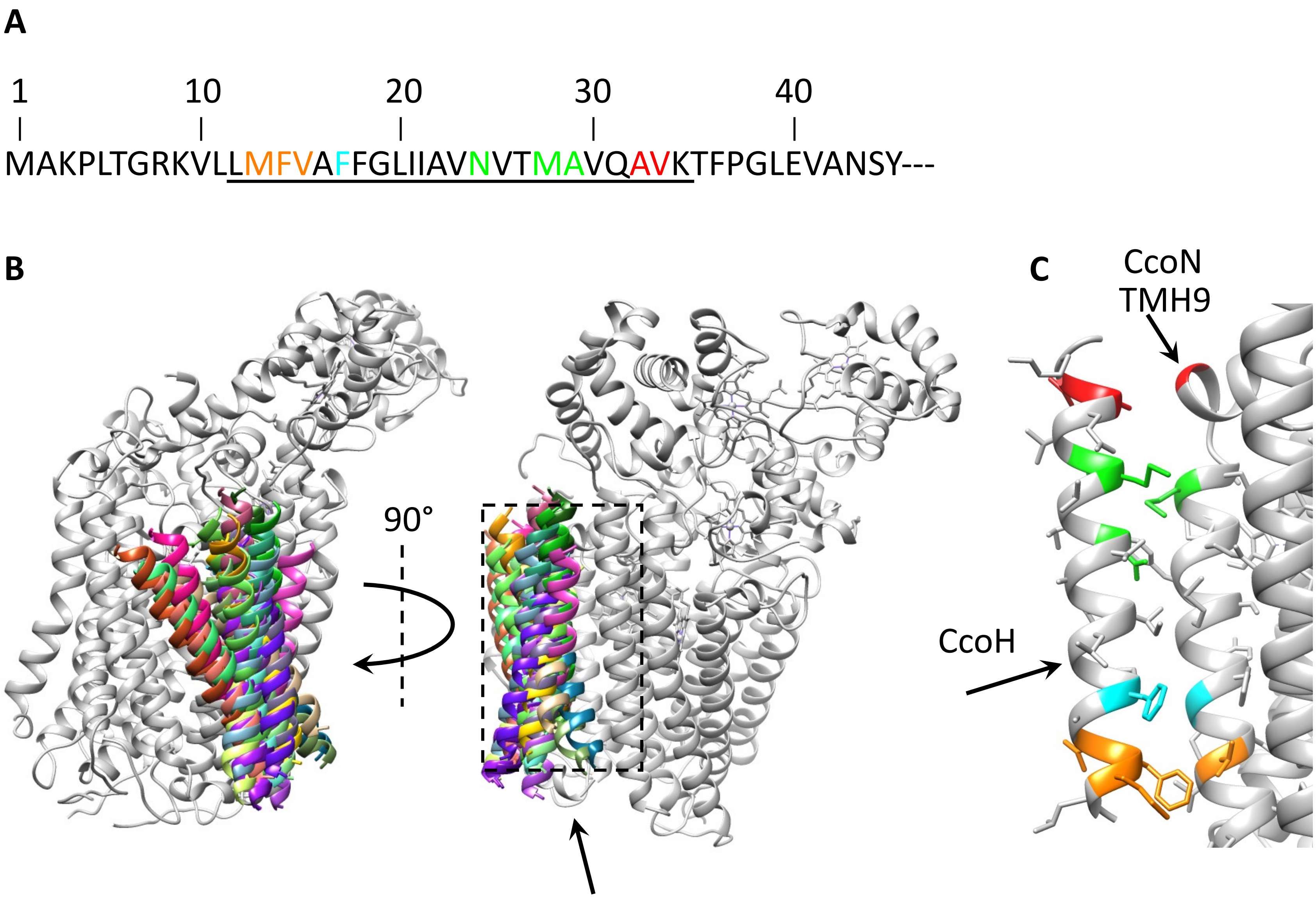

***Figure S8. Steimle et al.,***

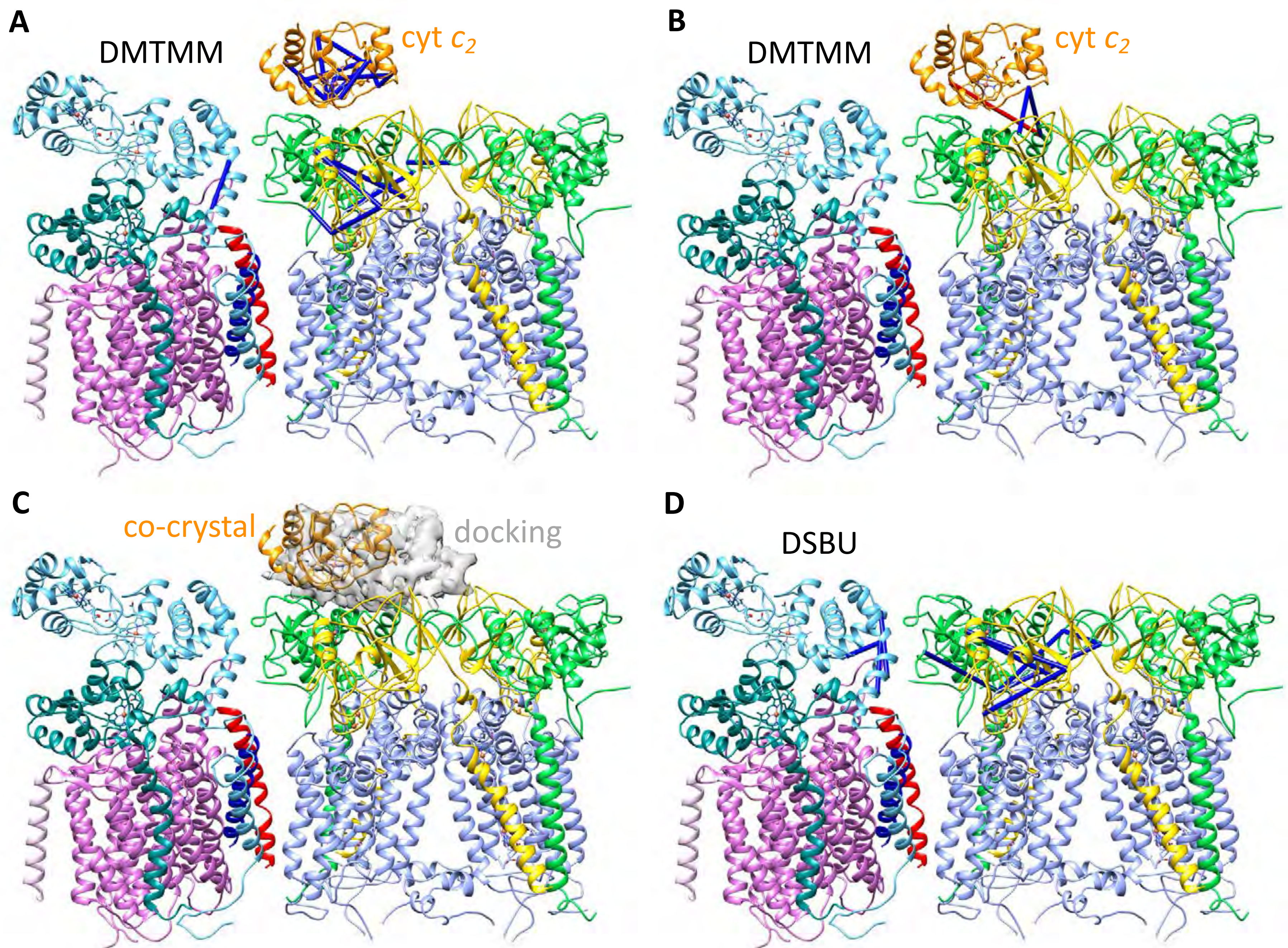

*Figure S9. Steimle et al.,*

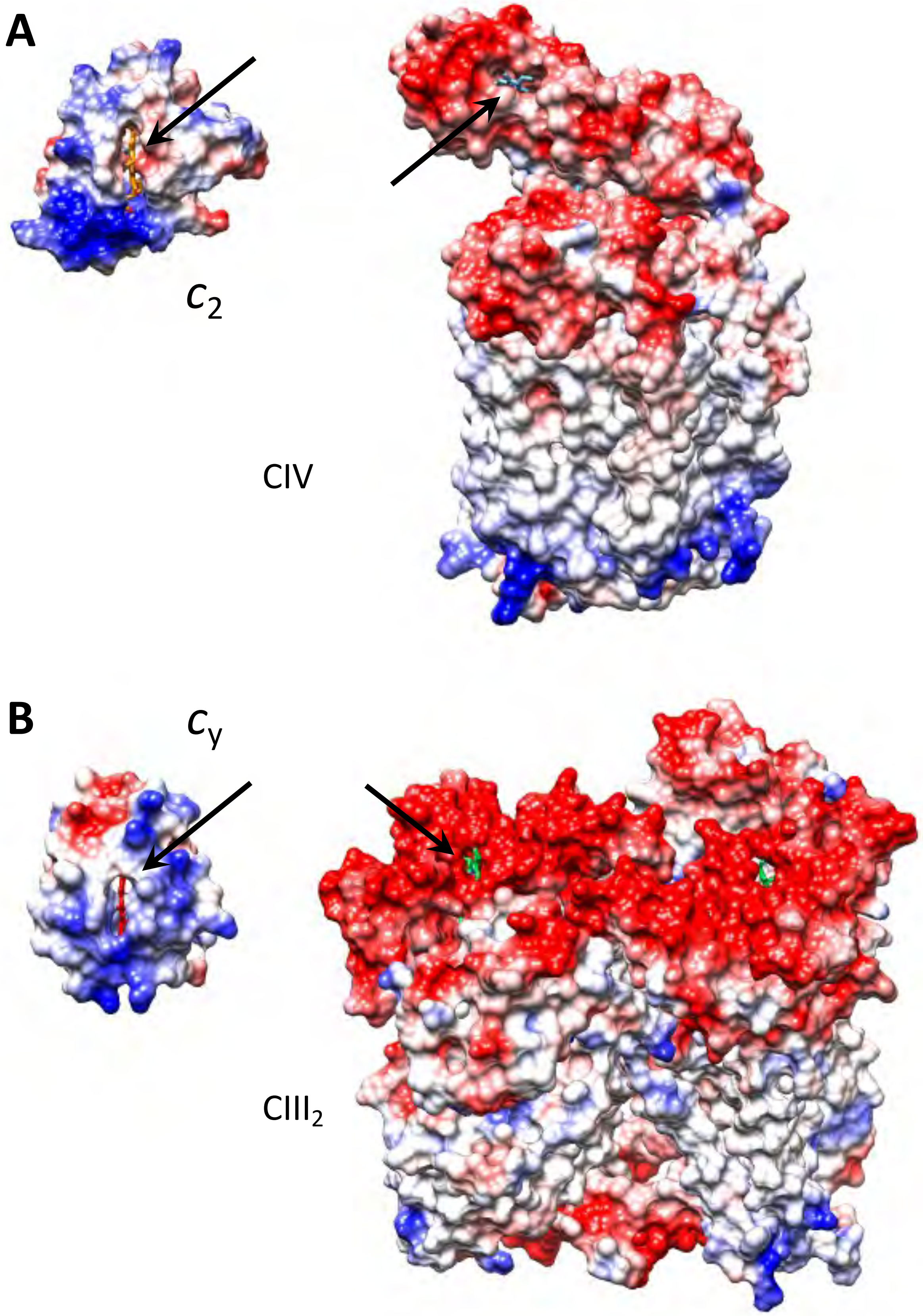

*Figure S10. Steimle et al.,*

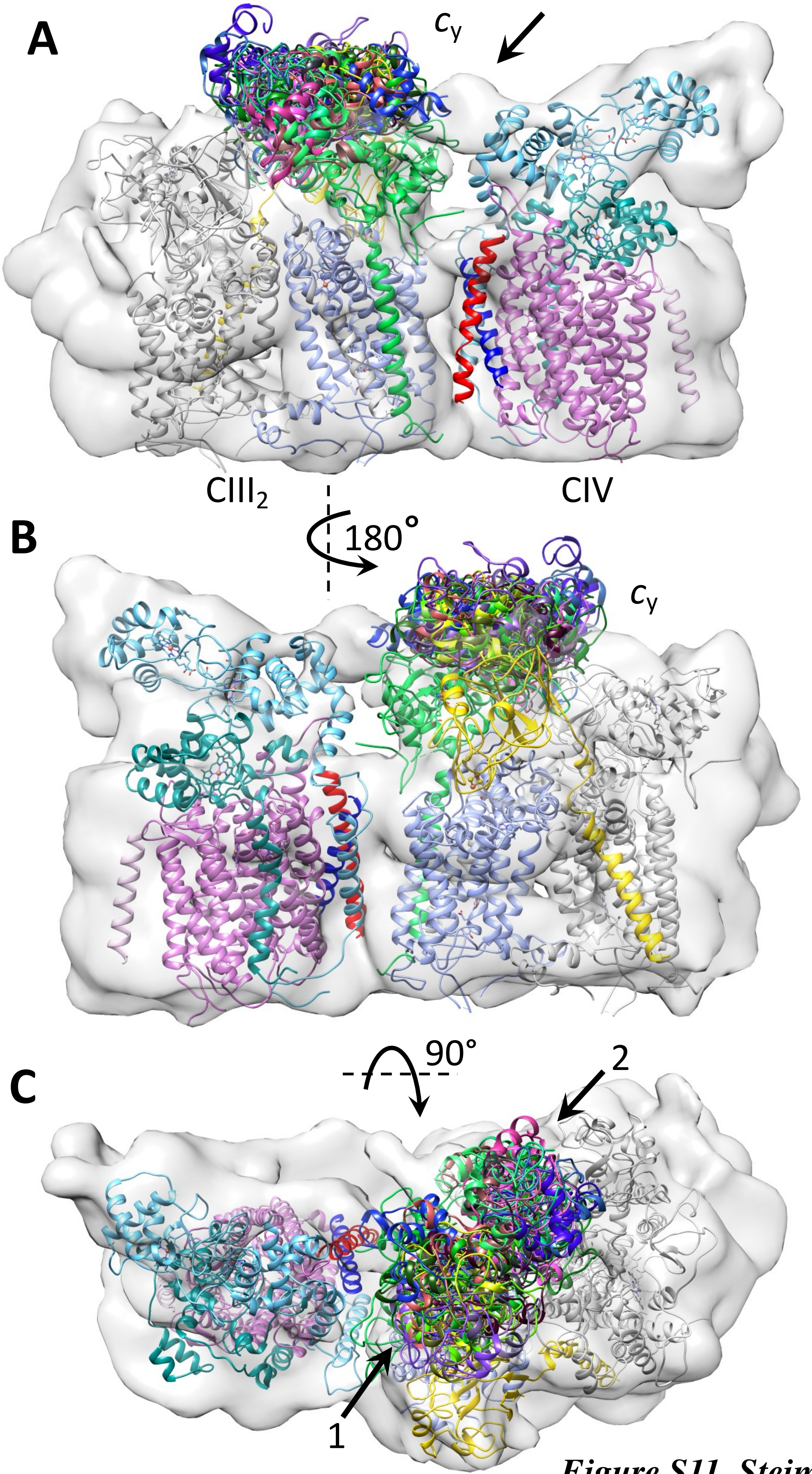

***Figure S11. Steimle et al.,***

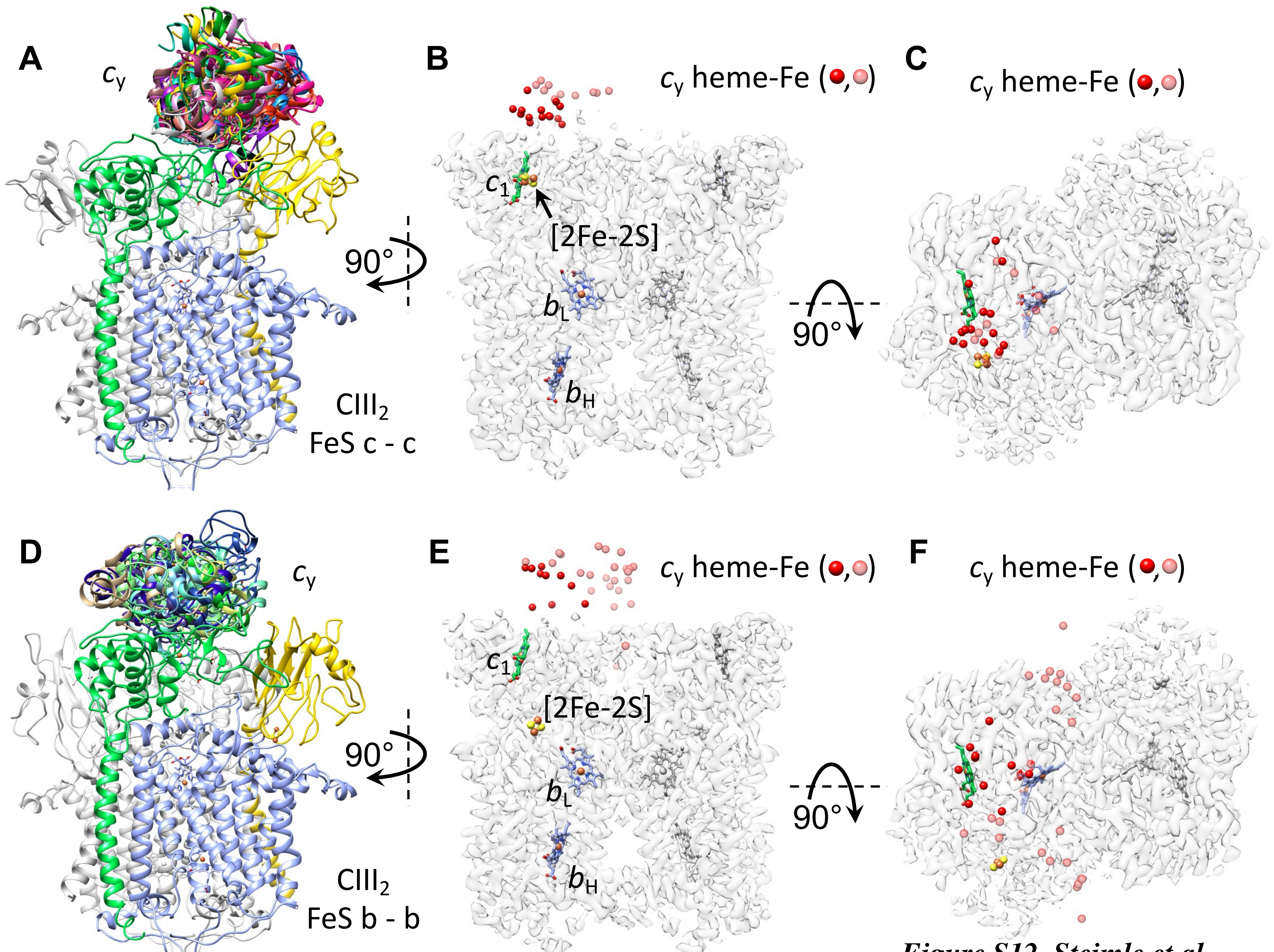

***Figure S12. Steimle et al.,***
